# Characterization of Self-Incompatibility Genes in *Brassica rapa* var. Toria and Yellow sarson

**DOI:** 10.64898/2026.03.25.714316

**Authors:** Hemal Bhalla, Kumari Ankita, Aman Ahlawat, Surabhi S. Rode, Kunwar Harendra Singh, Subramanian Sankaranarayanan

## Abstract

Self-incompatibility (SI), a reproductive mechanism that prevents self-pollen from fertilizing the ovule, is widespread in flowering plants, including the Brassicaceae family, where it promotes outcrossing, genetic diversity, and hybrid vigor. Although prevalent in *Brassica rapa*, an economically vital crop, it remains poorly characterized in widely grown varieties, such as toria and yellow sarson, with prior studies primarily focused on *Brassica napus*. Given its potential for hybrid breeding and crop improvement in rapeseed (*B. rapa*), we characterized key SI-regulatory genes, analyzing their phylogenetic relationships, structure-function dynamics, and expression patterns. Our results indicate sequence, structural, and functional homology as well as conservation with previously known candidates. This study identifies SRK, FER, and ARC1 as essential, while MLPK plays a minor role in SI for the varieties under study. Furthermore, we identified that SRK, FER, and MLPK activate ROS during the SI response, while ARC1 does not. Our findings establish a foundation for harnessing this natural system to integrate agriculturally important traits and sustain them across generations via outcrossing.

## 1. Introduction

Self-incompatibility (SI) is a genetic mechanism in various families of flowering plants that promotes outcrossing, increases genetic diversity and hybrid vigor, and helps avoid inbreeding depression. In the Brassicaceae family, which includes economically important oilseed, vegetable, and leafy green crops, SI is of the Sporophytic type. This system is governed by the polymorphic S-locus in a haplotype-dependent manner, such that matching haplotypes between stigma and pollen trigger an incompatible response (Nasrallah, 1997; Abhinandan et al., 2022; Bhalla et al., 2025a).

At the molecular level, the multi-allelic pollen ligand SP11/SCR (S-locus protein 11/S-cysteine-rich) interacts with the stigmatic receptor kinase SRK (S-locus receptor kinase) to orchestrate an incompatible response when haplotypes are identical (Kusaba et al., 2000; Takayama et al., 2001; Abhinandan et al., 2022; Huang et al., 2023; Bhalla et al., 2025a). This interaction activates downstream components including MLPK (M-locus protein kinase) (Kakita et al., 2007), which, together with SRK, phosphorylates and activates ARC1(Armadillo repeat-containing 1), leading to the ubiquitin-mediated proteasomal degradation of compatibility factors such as Exo70A1 (Exocyst 70A1), GLO1 (Glyoxalase 1), and PLDα1 (Phospholipase D alpha 1) (Figure 1A) (Samuel et al., 2009; Sankaranarayanan et al., 2015; Scandola and Samuel, 2019; Abhinandan et al., 2022; Bhalla et al., 2025a).

**Figure 1.**
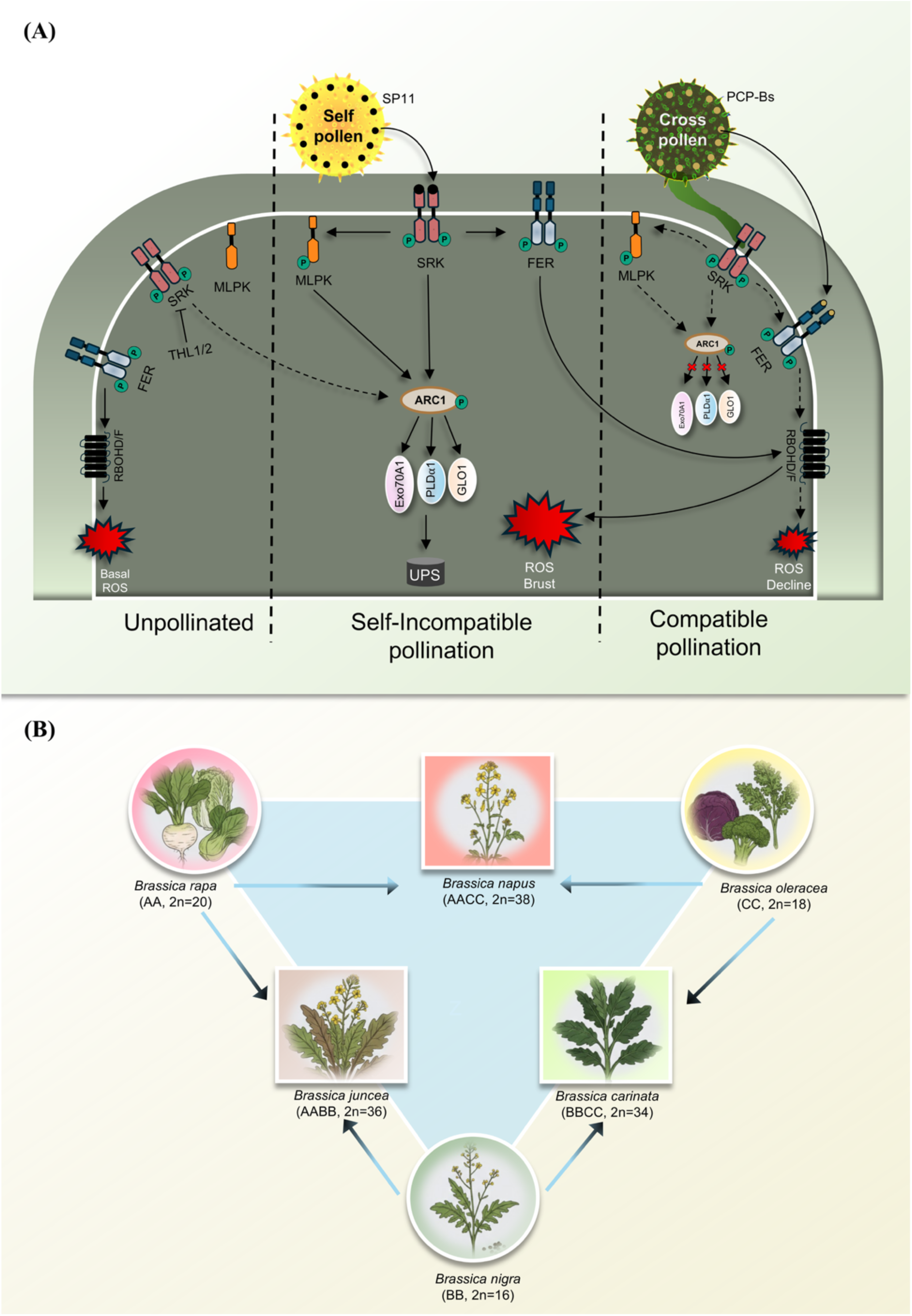
Overview of self-incompatibility in Brassicaceae. (A) Schematic model of Brassicaceae pollen-stigma signaling. In the unpollinated state, the SP11-SRK-ARC1 pathway remains inactive, whereas the FER-RAC/ROP-RBOHD/F pathway maintains basal ROS levels. THL1/2 maintains SRK in its inactive state. Maintaining the balance where the stigma is unfavorable for bacterial or fungal spore germination, while remaining favorable for compatible pollination. Upon self-incompatible pollination, the haplotype-specific interaction between SP11 and SRK phosphorylates MLPK. Together, SRK and MLPK then phosphorylate and activate an E3 ligase, ARC1, leading to the degradation of essential compatible factors required for the compatible pollen response. SRK also simultaneously activates the FER-RAC/ROP-RBOHD/F pathway, leading to a ROS burst and causing self-pollen rejection. During compatible pollination, the SP11-SRK interaction does not trigger ARC1 activation, thereby maintaining compatible factors at optimal levels, while PCP-Bs from compatible pollen suppress the FER-RAC/ROP-RBOHD/F pathway, leading to a decline in ROS and enabling pollen acceptance and tube formation. Note: Solid line indicates a functional pathway, and a dotted lines or red crosses indicates a non-functional pathway. **SP11**: S-locus Protein 11, **SRK**: S-Receptor Kinase (or S-locus Receptor Kinase), THL 1/2: Thioredoxin H-Like proteins, **ARC1**: Armadillo repeat-containing 1, **EXO70A1**: Exocyst 70A1, **GLO1**: Glyoxalase 1, **PLDα-1**: Phospholipase D alpha 1, **UPS**: Ubiquitin Proteasome System, **FER**: FERONIA, **RAC**: RAC GTPase, **ROP**: Rho Of Plants GTPase, **RBOHD/F**: Respiratory Burst Oxidase Homolog D/F, **MLPK**: M-Locus Protein Kinase, **ROS**: Reactive Oxygen Species, **PCP-B**: Pollen Coat Protein B-class, p indicates phosphorylation, (B) U’s triangle for the cultivated *Brassica* species. U’s triangle illustrates the genomic relationships among six cultivated *Brassica* species. Diploid progenitors: *B. rapa* (AA), *B. nigra* (BB), and *B. oleracea* (CC) hybridized naturally and underwent chromosome doubling to form allotetraploids: *B. napus* (AACC), *B. juncea* (AABB), and *B. carinata* (BBCC).

Concurrently, activated SRK also activates the receptor-like kinase FERONIA (FER), which triggers a respiratory burst oxidase homolog (RBOHD/F)-mediated reactive oxygen species (ROS) burst that further restricts pollen acceptance (Figure 1A) (Zhang et al., 2021; Huang et al., 2023). In contrast, during a compatible response, interaction between completely dissimilar haplotypes, these signaling cascades are attenuated, thereby preventing degradation of compatibility factors, limiting ROS accumulation, and permitting pollen acceptance and germination (Figure 1A).

Brassicaceae is a family of major economic importance, encompassing a wide range of food crops such as cabbage, broccoli, cauliflower, kale, turnip, pak choi, and radish; oilseed crops including canola, Indian mustard, and rapeseed mustard (rapeseed); and condiments such as black, brown, and white mustard. The family is also central to basic research, particularly through the model species *Arabidopsis thaliana* (Zhang et al., 2025). Within this family, the genus *Brassica* stands out, comprising six major crop species: three diploids—*Brassica nigra* (BB genome), *Brassica rapa* (AA genome), and *Brassica oleracea* (CC genome)—and three allotetraploids—*Brassica juncea* (AABB genome), *Brassica carinata* (BBCC genome), and *Brassica napus* (AACC genome) (Figure 1B) (Zhang et al., 2025).

Among these six species, five, excluding *B. oleracea,* are commonly referred to as oilseed *Brassica* or rapeseed mustard crops and are of high economic value for oil production worldwide (Thakur et al., 2017; Sharma et al., 2025). One of these, *Brassica rapa* L., widely cultivated as rapeseed, is valued for its oil content, favorable fatty acid profile, quality protein composition, preservative properties, and its use as a vegetable, flavoring agent, and condiment (Abul-Fadl et al., 2011; Cartea et al., 2019; Grygier, 2023). Despite this economic significance, the species often lacks high yield, robust disease resistance, and other agronomically important traits that could be introgressed through hybrid breeding and the exploitation of genetic diversity (Wang et al., 2022).

Previous work on self-incompatibility (SI) systems and their application in hybrid seed production and crop improvement has largely focused on canola. In contrast, research on SI in *Brassica rapa* L. has remained relatively limited, despite the species’ morphological and genetic diversity and its substantial economic importance. This has provided strong impetus to investigate its incompatibility systems and to harness the naturally occurring SI pathways in this species for breeding and crop improvement strategies. Our study has characterized the SI system in rapeseed mustard, providing insights into its similarities to previously known molecular players, their protein structures, and their functions. Our results suggest that the two-variety system, comprising self-incompatible toria and self-compatible yellow sarson lines of *B. rapa,* holds considerable potential for further functional studies and for deployment in breeding applications.

## 2. Materials and methods

### 2.1 Plant material and growth conditions

*Brassica rapa* varieties toria (TOR; self-incompatible) and yellow sarson (YS; self-compatible) seeds were obtained from the Indian Council of Agricultural Research - Directorate of Rapeseed-Mustard Research (ICAR-DRMR), Bharatpur, Rajasthan, India. Seeds were surface-sterilized using 0.05% SDS in 70% ethanol and plated on half-strength Murashige and Skoog (MS) medium, stratified at 4°C for 3-7 days, then grown for 10 days at 22°C with a 16 h:8 h light: dark cycle and 70% relative humidity. Plantlets were subsequently transferred to pots under the same conditions for acclimatization before experiments.

### 2.2 *In-silico* analysis and structure prediction

Sequences of the genes; *SRK* (Supplementary Table S3), *FER1* (Supplementary Table S4), *MLPK* (Supplementary Table S5), and *ARC1* (Supplementary Table S6) were retrieved from the NCBI database (Sayers et al., 2022) by performing nucleotide BLAST searches against species of the *Brassica* U’s triangle (Figure 1B) and *Arabidopsis*, using a 75% identity cutoff relative to the respective query sequences. Multiple sequence alignments of these gene sequences were performed using CLUSTAL W (Thompson et al., 1994). The representation and visualization of the multiple sequence alignment was conducted using N.ESPript (Gouet et al., 1999). Phylogenetic trees were constructed with MEGA11 (Tamura et al., 2021) using the maximum likelihood method with 1000 bootstrap replications, the Tamura–Nei nucleotide substitution model, and default parameters; the resulting trees were visualized using iTOL (Letunic and Bork, 2024).

Secondary structures of the protein sequences SRK, FER1, MLPK, and ARC1 were predicted using the PSIPRED server (McGuffin et al., 2000). The three-dimensional (3D) structural models were predicted using AlphaFold3, an AI model by Google DeepMind and Isomorphic Labs (Abramson et al., 2024). For each query sequence, the template-based modeling mode was selected, with all other parameters set to their defaults. Evolutionarily conserved functional domains in these proteins were identified using the NCBI Conserved Domain Database (Marchler-Bauer et al., 2009).

### 2.3 Cross-compatibility assay

Flowers of *Brassica rapa* var. toria and yellow sarson were emasculated on the day before crosses were conducted. In-planta manual pollination was conducted in both directions. The plants were grown under optimal conditions, and seeds were harvested to determine the number per pod.

Stigmas were pollinated in both directions for 12 h. These stigmas were subjected to an aniline blue assay to determine pollen attachment and pollen tube count.

### 2.4 Cloning and sequencing of genes

Primers were designed from previously known sequences in NCBI GenBank (Supplementary Table S1) (Sayers et al., 2022) using Primer3Plus (Untergasser et al., 2012). Total RNA was extracted using the RNeasy plant mini kit (Qiagen, Catalog: 74904), and first-strand cDNA was synthesized using the Thermo Scientific First Strand cDNA synthesis Kit (Catalog: K1612). The resulting cDNA was used as a template in PCR to optimize amplification of the product (Supplementary Table S2). The amplified products were cloned into the pCR4 -TOPO TA vector using the TOPO TA Cloning Kit for Sequencing (Catalog: 450030). The recombinant constructs were introduced into *E. coli* DH5α cells and selected on LB plates containing either ampicillin or kanamycin. Successful cloning and transformation were confirmed by colony PCR and restriction digestion. At least two independent plasmid clones were subjected to Sanger sequencing to verify the insert sequence.

### 2.5 Designing of oligonucleotides (S-ODNs and AS-ODNs)

Sense and antisense oligodeoxyribonucleotide (S-ODN and AS-ODN) pairs were designed using Sfold (Supplementary Table S8) (Ding, 2001, 2003; Ding et al., 2004). Potential off-target effects were assessed using NCBI BLAST (Camacho et al., 2023). ODNs were synthesized by Sigma-Aldrich with phosphorothioate modifications at the three terminal bases on both the 5’- and 3’-ends to enhance stability and prevent nuclease degradation.

### 2.6 *In-vitro* treatment of stigma and pollination assay

For the *in vitro* oligonucleotide treatment, the top-dripping method previously described was used (Zhang et al., 2024). Briefly, in this method, flowers were placed on Pollen Germination Media (PGM) with agar (5 mM CaCl_2_, 5 mM KCl, 0.01% H_3_BO_3_, 1 mM Mg_2_SO4·7H_2_O, 10% sucrose, and 0.8% agarose at pH 7.5). Further, 2 µL of 30 µM S-ODN, AS-ODN (with 0.0125% Tween 20), or mock (nuclease-free water with 0.0125% Tween 20) was pipetted onto each stigma. Following treatment, stigmas were placed at 22°C with 70% relative humidity for 1.5 h.

These treated stigmas were manually pollinated with self- or cross-pollen and incubated for an additional 12 h under the same conditions as previously described, followed by aniline blue staining to assess pollen attachment and pollen tube count.

### 2.7 Aniline blue staining

Stigmas were kept in Carnoy’s Fixative (acetic acid: ethanol = 1:3) for decolorization for 2 h. Samples were then transferred to 1 M NaOH at 55 °C for 25 min to soften the tissue. Next, the stigmas were stained with aniline blue (0.1% in 108 mM Potassium phosphate tribasic buffer - K_3_PO_4_, pH 11) for 2 h, and pollen grains and tubes were visualized with a Leica epifluorescence microscope and captured with a KI3C digital camera.

### 2.8 Nitro blue tetrazolium (NBT) assay for ROS estimation

Mock-, Sense-, or Antisense ODN-treated stigmas, collected unpollinated or 15, 30, and 60 minutes after self-incompatible pollination, were incubated in NBT staining solution (1mg/ml in 200 mM phosphate buffer, pH 7, and 0.02% Silwet) for 2.5 hours. Samples were then transferred to an ethanol: glycerol: acetic acid solution (3:1:1) and heated at 90°C for 10 minutes to clear the tissues. Finally, the stigmas were mounted in chloral hydrate: glycerol: water (8:1:2) for further clearing and visualized under a Leica microscope with a brightfield setup.

### 2.9 Reverse-transcription PCR

For RT-PCR, primers were designed using Primer3Plus (Untergasser et al., 2012) with default settings (Supplementary Table S9). Primer specificity was assessed using BLAST against *Brassica rapa* Chiifu V 4.0 genome in the BRAD database (Chen et al., 2022). For RT-PCR, stigmas were treated with oligonucleotides as previously described for 1.5 h. After treatment, stigmas were used for RNA isolation with the RNeasy Plant Mini Kit (Qiagen, Catalog: 74904) according to the manufacturer’s protocol. An equal amount of RNA was used to synthesize cDNA using the Thermo Scientific First Strand cDNA synthesis kit (Catalog: K1612). The cDNA was used in an RT-PCR reaction with 2X Takara Master Mix (Catalog: RR310A). All RT-PCR reactions were repeated 3 times, and *BrActin* 7 (NCBI Accession: KU851921) was used as a control.

### 2.10 Data analysis

For the cross-compatibility assay, seeds were counted from 5 pods each, with three biological replicates (n=15) (see Supplementary Data 2), and pollen attachment and tube penetration were measured in 5 independent replicates (n=5) (see Supplementary Data 2). For the AS-ODN-based assay, n = 9 stigmas were analyzed per treatment for pollen attachment and pollen tube penetration (see Supplementary Data 2). To quantify the intensities of the Nitro blue tetrazolium (NBT) assay, fluorescent intensities were randomly measured across the stigmatic papillary cells using a 120×120 μm region of interest (ROI) in ImageJ (see Supplementary Data 2) (Schneider et al., 2012). For RT-PCR analysis, the band intensities were calculated using ImageJ (Schneider et al., 2012) (see Supplementary Data 2).

Graphs were plotted using GraphPad Prism 8.0.2 (www.graphpad.com). One-way ANOVA (P < 0.05) was used to calculate statistics; p < 0.05, * indicates a significant difference, and ns indicates no significant difference.

## 3. Results

### 3.1 Phenotypic and cross-compatibility characterization of self-incompatible toria and self-compatible yellow sarson in rapeseed mustard

To investigate self-incompatibility in rapeseed mustard, we grew two commercially available *Brassica rapa* L. var. toria (self-incompatible) and yellow sarson (compatible). The plants were optimized for growth under a 16-hour light and 8-hour dark cycle at 22 °C and 70% relative humidity. Germination occurred within 7 days post-plating, and flowering was observed around 30 days for toria, whereas yellow sarson flowered approximately 35-40 days post-germination (Figure 2A). The flowers were cruciform and yellow in color, comprising all the four whorls - sepals (four, narrow, spreading, yellow green and about 5-8 mm long), petals (four, obovate, bright yellow, arranged in a cross with a slender claw and about 6-11 mm long), stamens (tetradynamous arrangement - 4 long and 2 short), and pistil (superior elongated ovary, with a short style and capitate stigma) (Figure 2B). Floral development from the bud stage to flowering took about 48 hours, with stage 1 as the bud stage, stage 2 as a slightly opened flower, and stage 3 as a fully opened flower, with the highest receptivity to pollination at stage 2, as observed in various crosses conducted in this study (Figure 2C).

**Figure 2.**
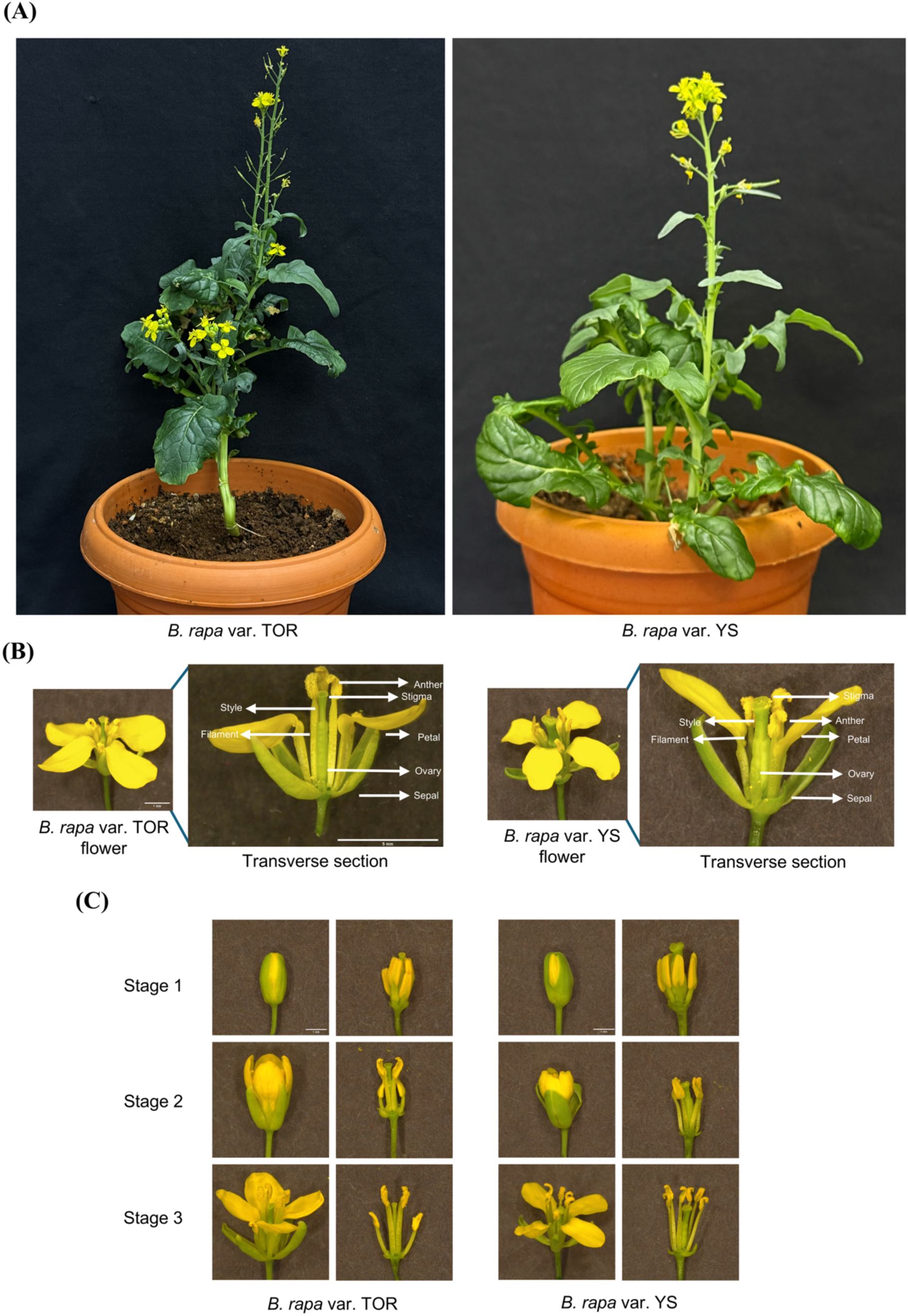
Morphological overview of *Brassica rapa* var. toria (TOR) and yellow sarson (YS). (A) Mature plants from the self-incompatible variety toria (Left) and the compatible variety yellow sarson (Right). (B) Close-up of flowers from each variety, highlighting key floral parts and transverse section. Scale bars: 1 mm and 5mm (C) Sequential floral developmental stages from complete flower and transverse section: stage 1: unopened bud, stage 2: slightly opened flower, and stage 3: opened flower from each variety showing stamen and pistil. Scale bar: 1 mm

Next, we examined cross-compatibility between the two varieties, which is crucial for any system used to dissect and manipulate self-incompatibility. Flowers were emasculated at stage 1 (Figure 2C) and subsequently pollinated with pollen from both self- and cross-compatible donors (Figure 3D-F). Aniline blue assay revealed that pollen attachment was comparable in both cross-pollinations with compatible pollination, while self-pollinated toria showed negligible pollen attachment. Furthermore, the number of pollen tubes penetrating the style was markedly reduced in both cross-pollinations relative to the compatible control (Figure 3D-F). Seed set assays showed that the cross with yellow sarson as the female parent and toria as the pollen donor produced a seed number similar to the compatible cross, whereas the reciprocal combination yielded slightly fewer seeds (Figure 3A and 3B). Seeds from compatible and yellow sarson (♀) × toria (♂) crosses were yellow, whereas seeds from incompatible and toria (♀) × yellow sarson (♂) crosses were brown (Figure 3C). In both crosses, the seed sizes were comparable and smaller than those obtained by self-pollination. Furthermore, the F1 generation seeds showed approximately 100% germination, further supplementing the cross-compatibility trait (Supplementary Figure S1 and Supplementary Data 2).

**Figure 3.**
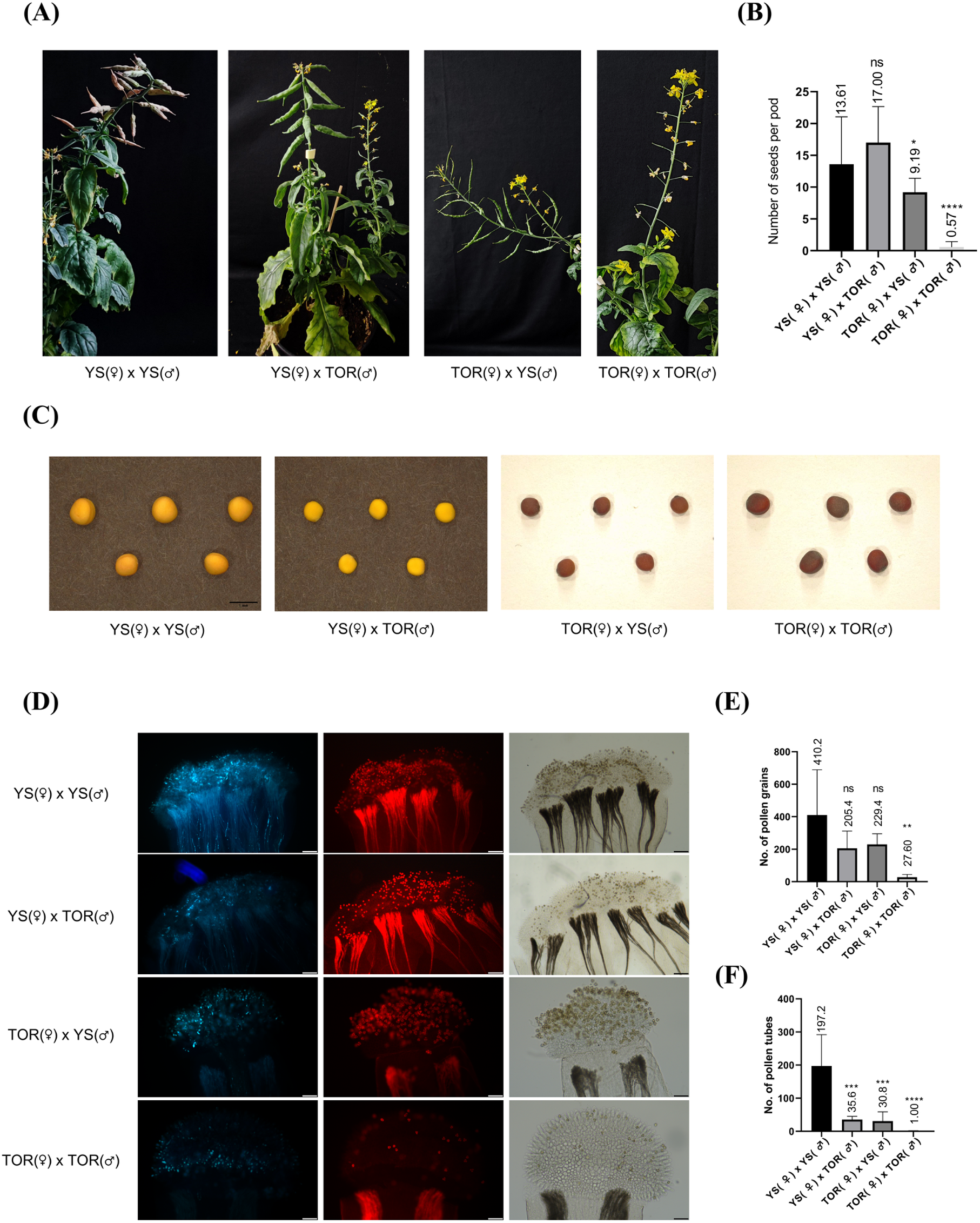
Cross compatibility between *B.rapa* var. toria and yellow sarson. (A) Mature plants showing pods from various in-planta crosses (seed-set assay) between *B.rapa* var. toria and yellow sarson. (B) Graph indicating the number of seeds per pod in various crosses between *B.rapa* var. toria and yellow sarson. Note: the values over the scale bar indicate average seeds per pod; One way ANOVA, *P < 0.05; n = 15, ns = not significant (C) Morphology of seeds collected from various crosses between *B.rapa* var. toria and yellow sarson. (D) Representative images of aniline blue assay indicating the cross-compatibility between *B.rapa* var. toria and yellow sarson. Note: Scale bar: 200 µm (E) Graph indicating the number of pollen grains in various crosses between *B.rapa* var. toria and yellow sarson. Note: One way ANOVA, *P < 0.05; n = 5, ns = not significant (F) Graph indicating the number of pollen tubes in various crosses between *B.rapa* var. toria and yellow sarson. Note: One way ANOVA, *P < 0.05; n = 5, ns = not significant

### 3.2 Conservation and phylogenetic characterization of self-incompatibility regulators in Brassica rapa

To investigate the role of the key self-incompatibility-associated genes *SRK*, *FER1*, *MLPK*, and *ARC1* in Brassicaceae (Figure 1A) (Abhinandan et al., 2022; Bhalla et al., 2025a), we retrieved their sequences from NCBI (Supplementary Table S3-S6) (Sayers et al., 2022). Gene-specific primers were designed from these sequences (Supplementary Table S1), and PCR conditions were optimized for all target genes (Supplementary Table S2). Clear bands of the expected sizes were obtained for all the genes, although non-specific amplification was observed for *MLPK* (Supplementary Figure S2A). The amplified products were cloned into the pCR4-TOPO TA vector using the TOPO TA cloning protocol and transformed into *E. coli* DH5α. Colonies were screened for the presence of the construct, and at least 2 positive colonies were verified by colony PCR (Supplementary Figure S2B). Restriction digestion confirmed successful cloning, yielding the predicted band sizes for all target plasmids (Supplementary Figure S2C). These clones were subjected to Sanger sequencing to analyze their sequences and compare them with previously reported homologs from other *Brassica* species.

To better understand the evolutionary relationships between the genes characterized in this study and their previously reported homologs, we retrieved homologous sequences using NCBI BLAST (Supplementary Table S3-S6) (Sayers et al., 2022) by filtering for 75% identity among species of U’s triangle and *Arabidopsis* (Figure 1B). Multiple sequence alignments were generated for *SRK*, *FER1*, *MLPK*, and *ARC1* (see Supplementary Data 3), and phylogenetic trees were constructed to visualize the evolutionary relationships between the newly characterized sequences and previously known candidates (Figure 4). *Arabidopsis thaliana* and *Arabidopsis lyrata* were utilized as outgroups for the tree construction (Figure 4). The largest number of homologs was obtained for *SRK*, consistent with its known allelic and haplotype diversity in *SP11/SCR-*mediated SI (Abhinandan et al., 2022). In general, two distinct clades were evident, with *Brassica rapa* var. toria *SRK* clustering closely with *BrSRK22* and showing strong bootstrap support (Figure 4A). *FER1* yielded only four sequences above the identity cutoff, yet it grouped more closely with *AtFER1* (Figure 4B). *MLPK* showed greater relatedness to *BrMLPK* from *B. rapa* subsp. *chinensis* and *Brassica napus* (Figure 4C), whereas *ARC1* exhibited higher similarity to *BrARC1* from yellow sarson, underscoring conservation between these two *B. rapa* varieties (Figure 4D). Together, these phylogenetic analyses indicate broad conservation of *SRK*, *FER1*, *MLPK*, and *ARC1* across the species of U’s triangle and reinforce their functional importance in the self-incompatibility response.

**Figure 4.**
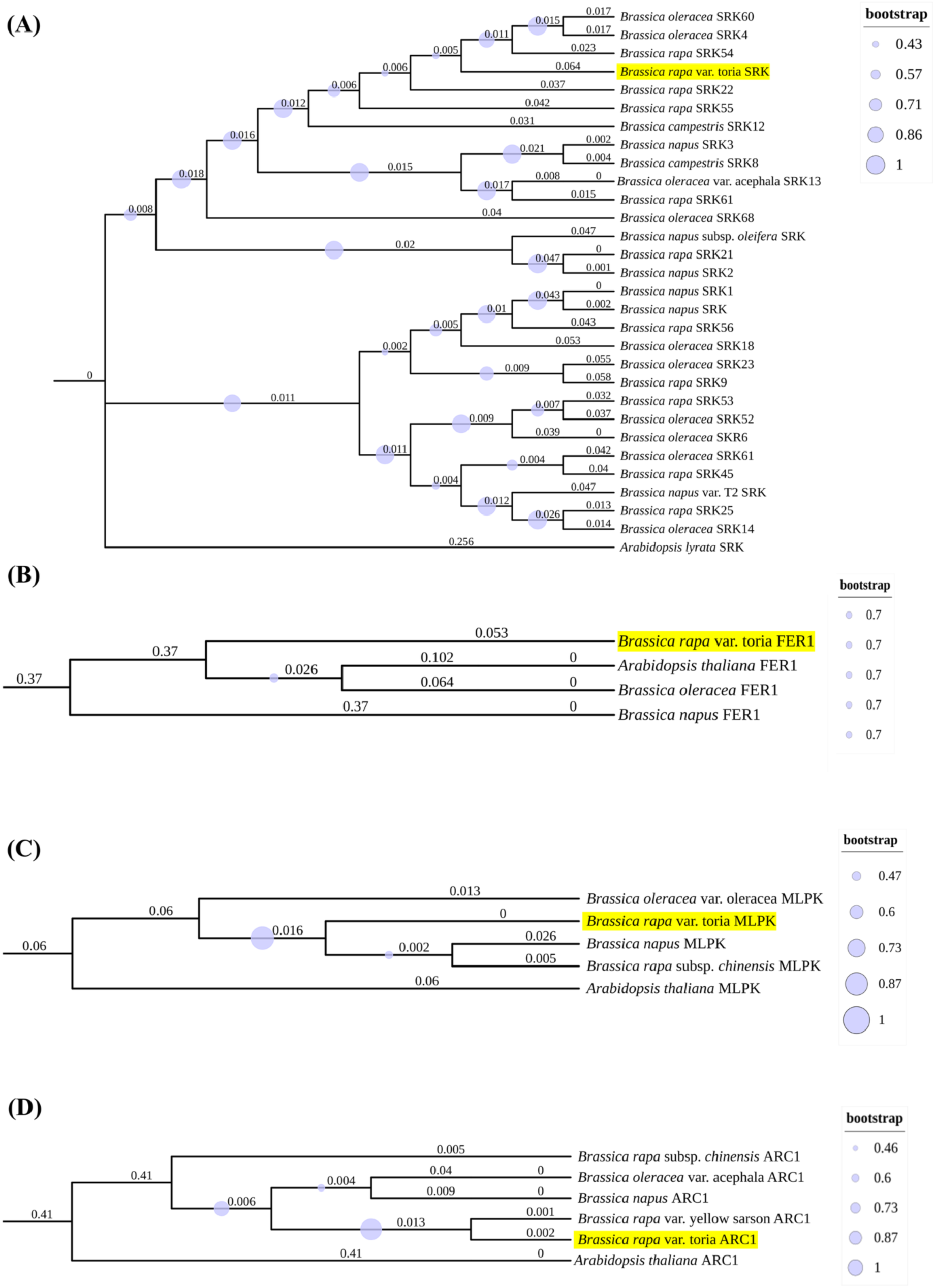
Phylogenetic analysis of genes under study. Phylogenetic tree of (A) *SRK*; (B) *FER1*; (C) *MLPK*; and (D) *ARC1* genes from *Brassica spp.* and *Arabidopsis* (Supplementary Table S3-S6) inferred from multiple sequence alignment (see Supplementary Data 3), using the maximum likelihood method. Note: Bootstraps: 1000 replicates

### 3.3 Structure-function, and *in silico* characterization of self-incompatibility regulators in Brassica rapa

To understand the functional properties of SRK, FER1, MLPK, and ARC1, it is essential to examine their structural organization, which provides key insights into the structure-function relationship. We first used PSIPRED to predict local spatial conformation and secondary structure (McGuffin et al., 2000). For SRK (829 amino acid residues; Supplementary Data 1), secondary structure analysis revealed a distribution of helices, coils, and strands across the protein, with high prediction confidence (Supplementary Figure S4A). The catalytic serine/threonine kinase domain at the C-terminal of SRK was predicted to have a higher helical and coiled structure, while the N-terminal domain was mainly composed of β-strands (Supplementary Figure S4A and Supplementary Table S7). The *Catharanthus roseus* Receptor-like Kinase 1-like (CrRLK1L) family protein FER1 (896 amino acid residues; Supplementary Data 1) predictions again yielded β-strands, α-helices, and coiled structures with a high confidence value (Supplementary Figure S4B). The extracellular malectin-like domain in the C-terminal consisted of coils and stretches of β-strands, while the cytosolic serine/threonine kinase catalytic domain at the N-terminus combined short β-strands with a predominance of α-helices and coils (Supplementary Figure S4B and Supplementary Table S7). For MLPK (418 residues), secondary structure prediction indicated a few short β-strands interspersed with higher helical content and coiled segments (Supplementary Figure S4C). The N-terminal membrane-anchored catalytic domain was predicted to contain β-strands, helices, and coiled regions with a high confidence value of secondary structure prediction, while the C-terminal kinase domain was mainly composed of helices and coiled regions (Supplementary Figure S4C and Supplementary Table S7). The ARC1 (661 amino acid residues; see Supplementary Data 1) showed a predominantly α-helical and coiled architecture (Supplementary Figure S4D), with the N-terminal U box region known to underlie SI function (Stone et al., 2003), comprising random coils and helices (Supplementary Figure S4D and Supplementary Table S7).

Next, we modelled 3D structures using AlphaFold 3, which predicted atomic-level details of the protein backbone, topology, and folding (Figure 5). The SRK model had a pTM score of 0.6, indicating moderate confidence that the predicted fold resembles the native structure, and exhibited a high confidence pLDDT profile across most of the protein (Figure 5A). The Ramachandran plot analysis of the SRK structure indicated 89.8% residues in the most favored region, 8% in the additionally allowed region, 1% in the generally allowed region, and 1.1% in the disallowed region (Supplementary Figure S5A). The SRK structure revealed a multidomain architecture, with an N-terminal β-sheet-rich region corresponding to a mannose-binding lectin-type domain (B-lectin superfamily), followed by an S-locus glycoprotein domain implicated in SI, and a C-terminal PAN domain of the PAN_APPLE superfamily (domains were verified using the conserved domain database; Supplementary Figure S3A). Six conserved cysteine residues in each of the S locus glycoprotein and PAN domains likely contribute to structural stabilization via disulfide bonds. The C-terminal serine/threonine kinase domain adopted a helix and strand-rich fold interconnected by loops (Figure 5A and Supplementary Figure S3A). Domain assignments based on the NCBI Conserved Domain Database (Marchler-Bauer et al., 2009) further delineated the amino acid spans of these domains (Supplementary Table S7). These features mirror the experimentally resolved SRK9:SCR9 complex, which also contains two lectin-like domains, an EGF-like domain, and a PAN_APPLE domain stabilized by conserved cysteines, with the PAN region mediating ligand-independent homodimerization and also interaction with SCR/SP11 to drive self-incompatibility (Naithani et al., 2007; Ma et al., 2016). The *in-silico* study of SRK revealed strong structural and domain-level similarities with experimentally derived SRK9 features, further supporting its role in self-pollen rejection.

**Figure 5.**
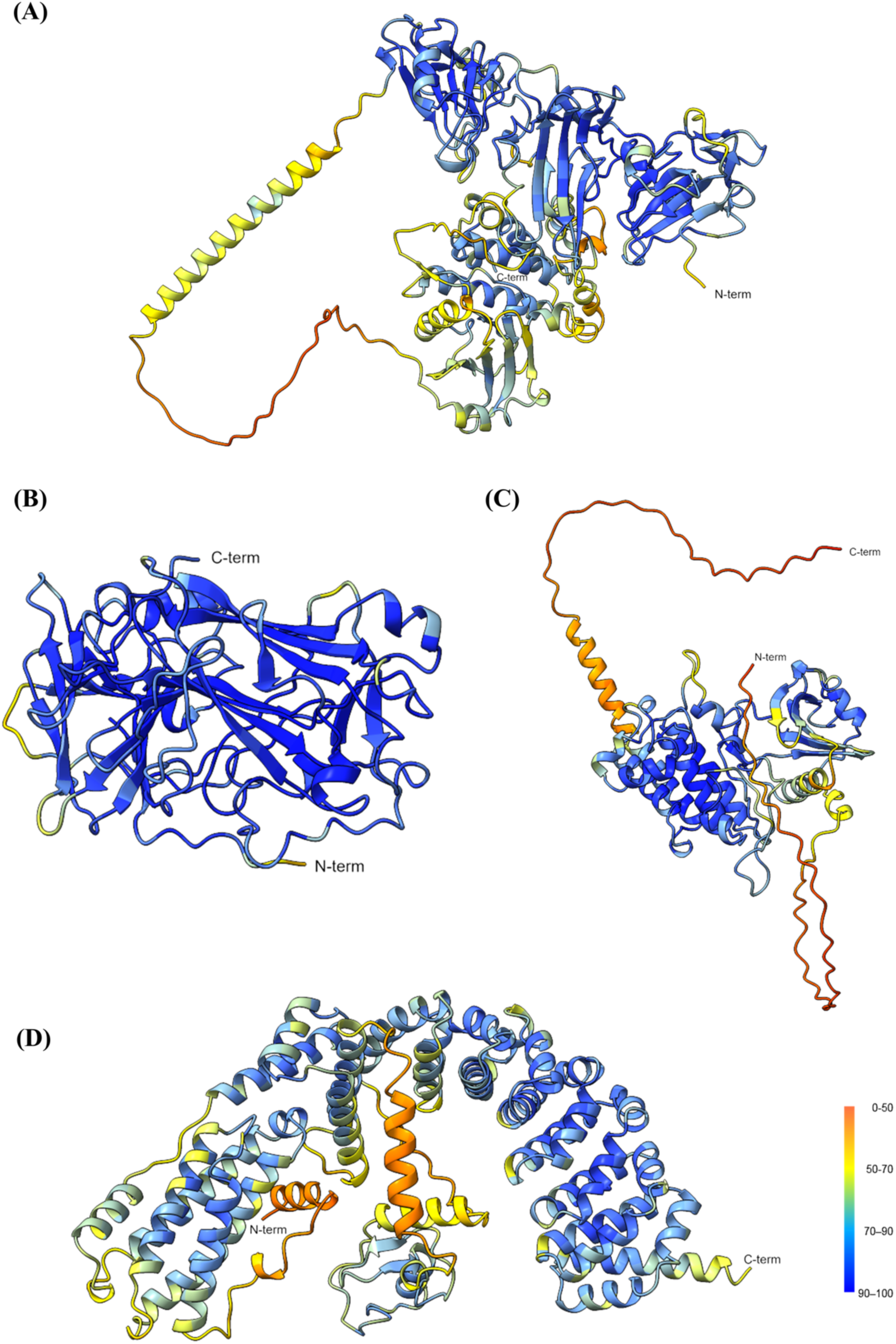
Structure prediction of various proteins regulating self-incompatibility in rapeseed mustard. Three-dimensional (3-D) structure predictions of (A) SRK; (B) FER1; (C) MLPK; and (D) ARC1 proteins using AlphaFold3. The structures are colored as per pLDDT scores. Dark blue color indicates very high accuracy (pLDDT >90), light blue indicates high confidence (90 > pLDDT > 70), yellow orange indicates low confidence (70 > pLDDT > 50), and red represents very low confidence (pLDDT < 50).

The AlphaFold3-predicted extracellular domain of FER1, which mediates pollen selectivity, showed a pTM score of 0.96 and a high pLDDT profile, indicating high prediction accuracy (Figure 5B). The structure conformed to the malectin-like superfamily surface receptor architecture annotated by the Conserved Domain Database (Supplementary Figure S3B and Supplementary Table S7) (Marchler-Bauer et al., 2009). The Ramachandran plot analysis of the FER1 extracellular domain placed 92.4% residues in the most favored region, 7.4% in the additionally allowed region, and 0.3% in the generally allowed region, indicating good quality of the predicted structure (Supplementary Figure S5B). The functional domains of the FER1 full-length protein were identified through the Conserved Domain Database (Supplementary Figure S3B and Supplementary Table S7). Full-length FER1 comprises three major functional modules: an extracellular malectin-like domain involved in pollen selectivity, a transmembrane segment anchoring the protein in the plasma membrane, and a cytosolic serine/threonine kinase (STK) domain responsible for downstream signaling (Supplementary Figure S3B and Supplementary Table S7) (Cheng et al., 2025). The STK domain includes an evolutionarily conserved ATP-binding site that transfers the γ-phosphoryl group from ATP to serine/threonine residues on substrate proteins, as inferred from domain-profile analysis. The experimentally determined structure of the *Arabidopsis thaliana* FERONIA extracellular domain reveals two tandem malectin-like regions linked by a β-hairpin that forms a ligand-binding site for pollen selection (Cheung, 2024), whereas the N-terminal extracellular part of FER1 in complex with the stigmatic ligands LLG2 (LORELEI-like GPI-anchored protein 2) and RALF23 (Rapid Alkalinization Factor 23) provides a molecular basis for blocking incompatible pollen attachment (Xiao et al., 2019; Bhalla et al., 2025b). However, the structural details of the cytosolic domain of FERONIA remain unknown due to the unavailability of experimental data. The presence of malectin-like extracellular domain at the N-terminal of FER1, with analogous topology to *At*FER1 crystal structure, directs the functionality of the FER1 extracellular domain in *Brassica rapa* (Figure 5B).

The AlphaFold3-predicted MLPK structure yielded a pTM score of 0.77 and a high-confidence core central domain flanked by intrinsically disordered N- and C-terminal segments (Figure 5C). The Ramachandran plot for MLPK placed 91.4% residues in the most favored region, 6.9% in the additionally allowed, 0.8% in the generally allowed, and 0.8% in the disallowed region (Supplementary Figure S5C). Conserved-domain analysis identified an N-terminal serine/threonine kinase domain characteristic of protein kinases, along with ATP- and substrate-binding sites (Supplementary Figure S3C and Supplementary Table S7).

Finally, AlphaFold3-predicted ARC1 adopted a folded, curved architecture with good overall quality (Figure 5D), with a pTM score of 0.72 and a Ramachandran profile placing 95.7% residues in the most favored region, 4.1% in the additionally allowed, and 0.2% in the generally allowed region (Supplementary Figure S5D). The 3D model revealed two main α-helical bundles: an N-terminal region harboring the U-box domain, which mediates ubiquitin ligase activity, and a C-terminal armadillo (ARM) repeat-containing domain implicated in substrate recognition (Figure 5D, Supplementary Figure S3D, and Supplementary Table S7). The NMR solution structure of the *Arabidopsis* U-box-containing PUB14 further supports this architecture, demonstrating that U-box plus ARM repeats form a compact, elongated fold suitable for E3-ligase function (Andersen et al., 2004). Although no experimental crystal structure of *Brassica* ARC1 has yet been reported, our *in-silico* analysis establishes a robust structural framework for future biochemical and structural studies of ARC1 in SI-related signaling.

### 3.4 Functional characterization of genes responsible for SI response

To functionally validate the roles of *SRK*, *FER1*, *MLPK*, and *ARC1* in the self-incompatibility-(SI) response of *Brassica rapa* var. toria, we suppressed these genes using antisense oligonucleotides (AS-ODNs). Chemically modified AS-ODNs bind to target mRNAs and thereby inhibit gene expression (Huang et al., 2020; Zhang et al., 2024). Based on previous studies, we adopted the top-dripping method (Zhang et al., 2024), in which stigma of self-incompatible toria flowers was treated with mock solution, sense, or anti-sense ODNs delivered dropwise from above (Supplementary Figure S6 and Supplementary Table S8). Following treatment, stigmas were pollinated with either self- or cross -compatible pollen for 12 h, after which aniline blue staining was performed. In self-pollinated toria stigmas, AS-ODN treatment significantly increased pollen attachment for all four genes compared with mock and sense controls (Figure 6 and Supplementary Figure S7). In contrast, under cross-pollination conditions, where pollen acceptance and tube growth occur naturally in the absence of SI, AS-ODN treatments had no significant effect on pollen attachment or tube penetration (Figure 6 and Supplementary Figure S7). The number of pollen tubes penetrating the style likewise increased in AS-ODN-treated-self-incompatible stigmas relative to mock and sense treatments for *SRK*, *FER1*, and *ARC1*, whereas AS-*MLPK*-treated stigmas showed no significant increase in pollen tube penetration (Figure 6 and Supplementary Figure S7).

**Figure 6.**
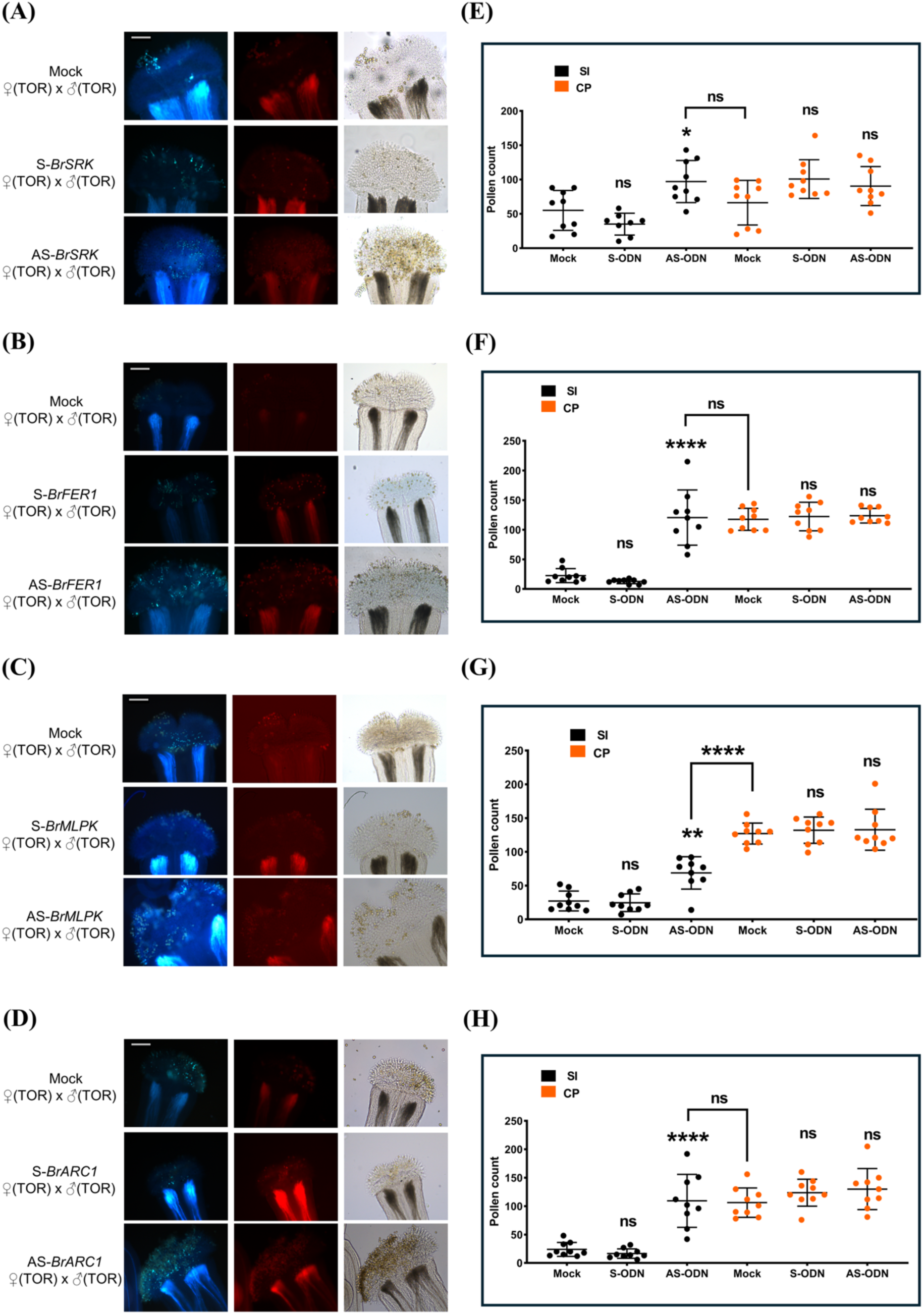
Functional characterization of genes during self-incompatibility response. (A-D) Representative aniline blue images of pollinated stigmas treated with Mock, Sense, and Anti-Sense ODNs of (A) *SRK*, (B) *FER1*, (C) *MLPK*, and (D) *ARC1*, showing pollen attachment and tube formation. Note: Aniline blue (left); Red channel (middle); Bright field (right). Scale bars: 300 μm (E-H) Graph indicating the number of pollen grains per stigma observed in (Figure 6A-D) & (Supplementary Figure S7A-D) treated with Mock, Sense, and Anti-Sense ODNs of (E) *SRK*, (F) *FER1*, (G) *MLPK*, and (H) *ARC1*. Note: Dots represent individual data points. One way ANOVA, *P < 0.05; n = 9, ns = not significant

Next, to evaluate the impact of ODN-mediated gene suppression on reactive oxygen species (ROS) levels, which are highly dynamic during the SI response (Figure 1A), particularly superoxide, we performed NBT staining on pollinated stigmas as previously described (Sankaranarayanan et al., 2025). We first optimized the time window for ROS analysis by monitoring ROS dynamics in unpollinated stigmas and at 15, 30, and 60 minutes after pollination (MAP) (Supplementary Figure S8). ROS levels rose at 15 and 30 MAP and then declined at 60 MAP, indicating a peak at 30 MAP (Supplementary Figure S8). Using this optimized time point, we assessed the effects of mock, S-ODN, and AS-ODN treatments on ROS accumulation in stigmas at 30 MAP (Figure 7). ROS levels were significantly reduced after AS-ODN treatment for *SRK*, *FER1*, and *MLPK*, whereas *ARC1* knockdown had no detectable effect on ROS (Figure 7).

**Figure 7.**
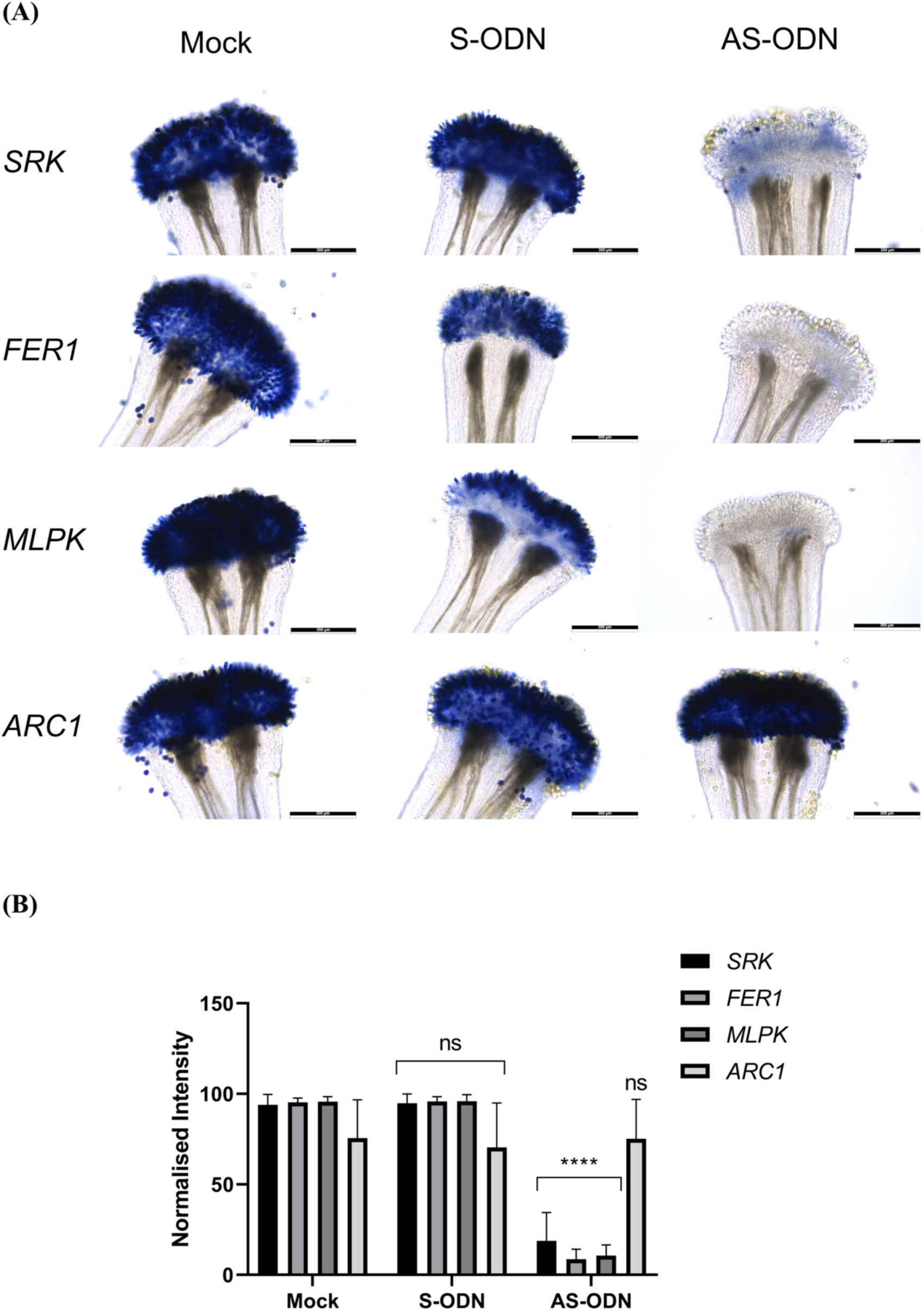
NBT staining assay for genes responsible for regulating self-incompatibility. (A) Representative images for various stigmas after 30 minutes of pollination, treated with Mock, Sense, and Anti-Sense ODNs for various genes under study, and stained using NBT. (B) Graph indicating the normalized intensity for various stigmas, pollinated for 30 minutes, treated with Mock, Sense, and Anti-Sense ODNs for various genes under study, and stained with NBT. Note: One-way ANOVA, *p<0.05, n=15

To confirm that ODN treatment indeed downregulated target gene expression, we performed RT-PCR on stigma tissue following mock, sense, and anti-sense ODN treatments. Gene-specific primers together with the reference gene *BrActin7* (NCBI Accession: KU851921) (Supplementary Table S9) were used for amplification. The RT-PCR analysis revealed a clear reduction in transcript levels of all four target genes in AS-ODN-treated stigmas, while expression levels remained comparable between mock and sense-treated controls (Figure 8 and Supplementary Figure S9).

**Figure 8.**
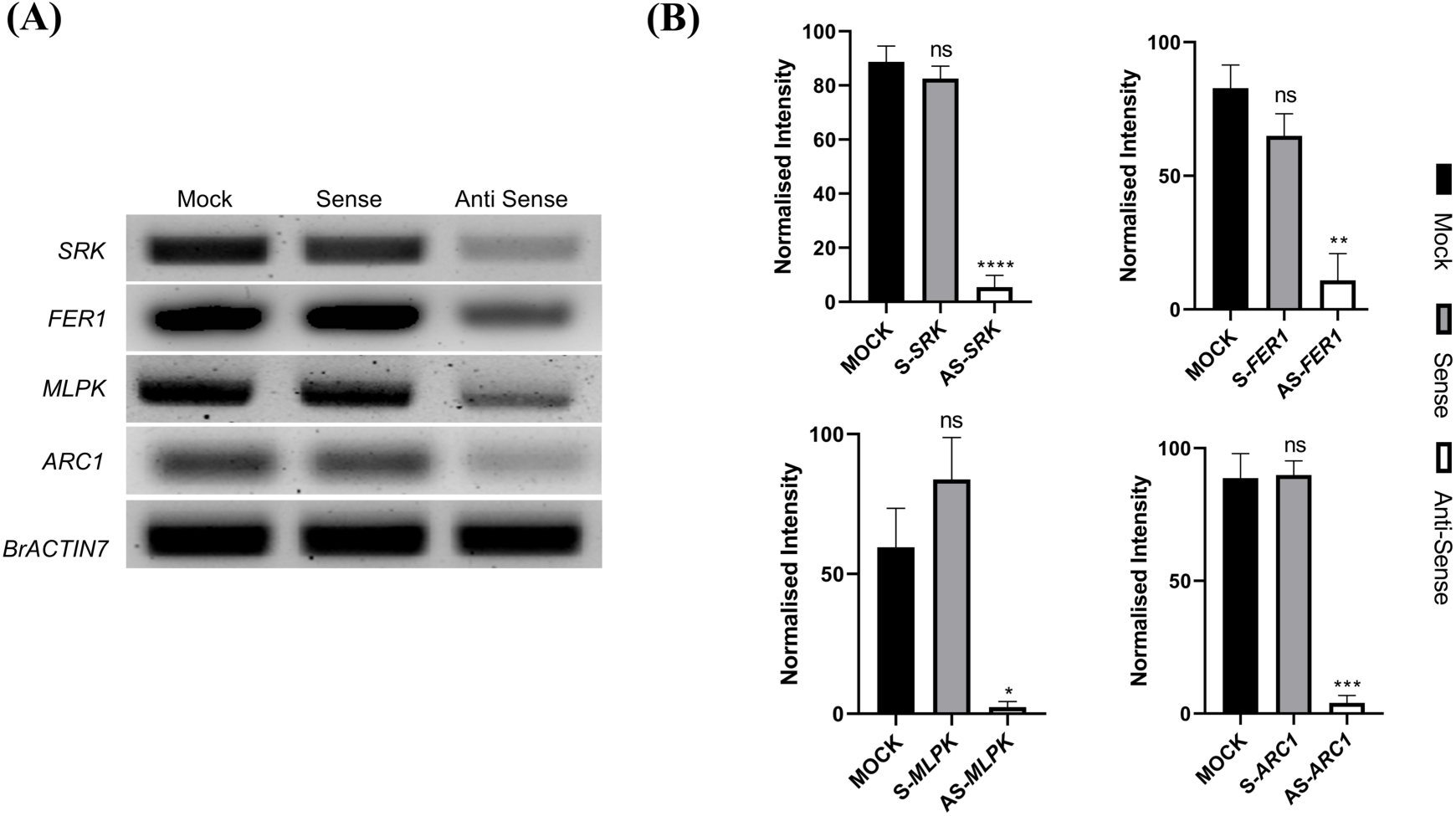
Expression analysis of ODN-treated stigmas. (A) Representative gel image for the *SRK*, *FER1*, *MLPK*, and *ARC1* treated with ODNs - Mock, Sense, and Anti-Sense. Note: *BrActin 7* (NCBI Accession: KU851921) was used as an internal control (B) Graph indicating the normalized intensity for various bands of *SRK*, *FER1*, *MLPK*, and *ARC1* treated with ODNs - Mock, Sense, and Anti-Sense. Note: *BrActin 7* (NCBI Accession: KU851921) was used for normalization and as an internal control; one-way ANOVA, *P < 0.05; n = 3, ns = not significant.

## 4. Discussion

Understanding self-incompatibility (SI) systems and exploiting them for crop improvement, hybrid breeding, and enhancement of genetic diversity is of central importance and has been successfully applied in several crops worldwide. The Brassicaceae family, encompassing economically vital vegetable, oilseed, and condiment species, has become a key model for dissecting incompatibility mechanisms and developing strategies to introduce traits such as high yield, disease resistance, and shatter tolerance. Despite this, the application of SI-based approaches in rapeseed mustard (*Brassica rapa* L.) remains limited, even though the species is well adapted to both irrigated and rainfed environments, amenable to sole or mixed cropping, and profitable at relatively low input and water requirements. Under these conditions, characterizing and harnessing the SI system for integrating major agronomic traits becomes particularly imperative.

Our study investigates the SI trait in *Brassica rapa* L. (rapeseed mustard), focusing on two commercially available and widely cultivated varieties: toria and yellow sarson. Toria is a self-incompatible, early-maturing form selected under breeding pressure, whereas yellow sarson represents a quality-type mutant selected for seed and oil characteristics (Thakur et al., 2017). Building on prior work, we optimized growth conditions for these varieties and report detailed observations on their morphology and reproductive development, within the context of elucidating self-incompatibility in rapeseed mustard. We further evaluated cross-compatibility between the two varieties, a prerequisite for utilizing SI in breeding schemes. Although the self-incompatible variety shows partial resistance to pollen tube penetration and seed set, it still permits sufficient fertilization and seed production to be integrated into hybrid-breeding programs.

The seed color and size differences observed in these crosses provide simple but informative phenotypic signatures of the underlying genetic behavior and physiological interactions. Uniformly yellow seeds from compatible crosses and from the yellow sarson (♀) × toria (♂) cross, in contrast to brown seeds from incompatible and toria (♀) × yellow sarson (♂) crosses, suggest a non-Mendelian, parent-of-origin effect more consistent with maternal or cytoplasmic control than with a simple nuclear dominant/recessive system. The reduced seed size in both yellow sarson (♀) × toria (♂) and toria (♀) × yellow sarson (♂) crosses, compared with fully compatible combinations, further indicates that the incompatibility status and the direction of the cross modulate resource allocation and seed development, likely reflecting disturbed pollen tube guidance, fertilization efficiency, or early post-fertilization interactions.

Furthermore, in this study, we sequenced all the major genes previously known to regulate self-incompatibility in these two varieties, which were submitted to NCBI (Supplementary Table S1) (Sayers et al., 2022) and provided insights into their conservation and relatedness to other species in the *Brassica* genus. Phylogenetic analysis of the newly characterized *Brassica rapa* var. toria sequences for *SRK*, *FER1*, *MLPK*, and *ARC1*, together with homologues from species of the U’s triangle and *Arabidopsis* as an outgroup, revealed close evolutionary relationships and shared ancestral origins. By examining these genes in *B. rapa* var. toria, we highlight their conservation both within the U’s triangle species and across closely related Brassicaceae, underscoring their functional relevance in SI.

Understanding a protein’s structural details, three-dimensional conformation, and catalytic domains is essential for interpreting its functional role in the biological system. *In silico* analyses using PSIPRED and AlphaFold3 predicted the secondary and tertiary structures of SRK, FER1, MLPK, and ARC1, while conserved-domain searches identified motifs characteristic of their roles in *Brassica* SI. The structural similarity between our SRK model and the experimentally resolved SRK9:SCR9 complex—particularly the presence of N-terminal α-helices, lectin-type domains, an S locus glycoprotein domain, and a PAN-like domain—highlights the mechanistic basis of SRK-mediated self-pollen rejection (Ma et al., 2016). Likewise, *in silico* analysis of FER1 identified an extracellular malectin-like receptor domain involved in pollen selection, a transmembrane region with a hydrophobic segment, and a cytosolic serine/threonine kinase domain that mediates downstream signaling. The predicted extracellular domain structure closely resembles the topology of the *At*FER1 crystal structure, supporting functional conservation across Brassicaceae (Cheung, 2024). Detailed structural studies of the cytosolic domain of FER1 in this family will further clarify its role in the signaling cascade. MLPK has previously been shown to bind SRK and phosphorylate downstream effectors to reinforce the rejection of self-pollen (Kakita et al., 2007), and functional analyses in *Brassica napus,* a functional analogue of *Brassica rapa* MLPK, have demonstrated its positive regulation of SI by modulating expression of *SRK* and *ARC1* (Chen et al., 2019). Our *in-silico* model of MLPK revealed conserved serine/threonine kinase, ATP-binding, and substrate-binding sites, consistent with a membrane-anchored kinase function. In contrast, the ARC1 structural model shows a conserved N-terminal ubiquitin ligase domain and a C-terminal armadillo (ARM) repeat-containing domain, supporting earlier reports of its E3 ligase activity and a positive role in SI (Stone et al., 2003). This bioinformatic analysis thus defines the structure-function relationship of ARC1, whose experimental structure determination by crystallography or other methods would deepen understanding of its mechanistic role in the Brassicaceae SI pathway.

Further, ODN-based transient suppression has previously been utilized and characterized in Brassicaceae (Zhang et al., 2024). Using the previously optimized technique, we suppressed major self-incompatibility genes, including *SRK*, *FER1*, *MLPK*, and *ARC1*. AS-ODN treatment increased pollen attachment and tube penetration, resulting in complete breakdown of SI for three genes, whereas *MLPK* suppression induced only a modest increase in pollen attachment relative to the compatible control. This breakdown underscores the functional importance of these genes in regulating the SI response in rapeseed mustard. In contrast, MLPK’s limited effect suggests a minor or redundant role in pollen acceptance/rejection, without compromising the overall SI response. Although previously the essential nature of MLPK has been illustrated (Murase et al., 2004; Chen et al., 2019), certain studies characterizing MLPK have also identified its non-essential role in SI response in certain *Brassica rapa* haplotypes and in SI-restored *Arabidopsis thaliana* (Murase et al., 2004; Kakita et al., 2008; Kitashiba et al., 2011; Ohata et al., 2023). Contrastingly, MLPK, previously characterized in yellow sarson by positional cloning (Murase et al., 2004), has been shown to be mutated and one of the integral reasons for its compatible nature, apart from insertion in SRK and deletion of a 89bp region in SP11/SCR (Murase et al., 2004; Fujimoto et al., 2006; Kakita et al., 2008). Furthermore, AS-ODN treatment did not alter compatible pollination, indicating that the previously optimized concentration and time point are also suitable for rapeseed mustard (Zhang et al., 2024). In alignment with previous studies, AS-ODN treatment of genes *SRK*, *FER1*, and *MLPK* inhibited ROS bursts at 30 MAP, whereas *ARC1* AS-ODN did not affect ROS levels (Huang et al., 2023), indicating the parallel nature of the ARC1- and FER-mediated pathways as well as the interactions between SRK, MLPK, and FER1 directly or indirectly with the ROS machinery. Interestingly, although *MLPK* AS-ODNs reduced ROS, we did not observe a corresponding complete breakdown of SI, suggesting a more intricate, layered regulatory network underlying self-incompatibility in Brassicaceae (Bhalla et al., 2025a). RT-PCR analysis confirmed downregulation of all four target genes after AS-ODN treatment; however, *MLPK*’s inability to break SI despite transcriptional suppression further supports the interpretation of its secondary or redundant contribution to the SI pathway, similar to the previously studied S29 haplotype of *B. rapa* (Ohata et al., 2023).

Overall, our results establish a robust platform for harnessing SI to improve major economic traits, enable hybrid seed production, and accelerate crop improvement in rapeseed mustard. The study reveals a well-developed, naturally occurring system in this species that can be exploited to enhance oil and seed yield, develop hybrids, build resistance to both biotic and abiotic stresses, and adapt varieties to non-traditional environments. The characterized system could be deployed in self-compatible *Brassica rapa* lines, as previously validated in *Arabidopsis thaliana* (Nasrallah et al., 2002; Zhang et al., 2019; Fujii et al., 2020). To our knowledge, no prior work has comprehensively characterized this system in rapeseed mustard, despite its traditional use in agriculture. A deeper understanding of the molecular players and their manipulation through breeding or emerging biotechnology approaches will provide valuable tools for breeders and farmers seeking to maximize the genetic potential of this important oilseed crop.

## Statements

### Data availability statement

All data that support the findings of this study are included in the manuscript and supplementary data of this article.

### Author contributions

HB, KA, and AA performed experiments and investigations. HB and KA were involved in characterization, cloning, sequencing, and functional investigation. AA performed the RT-PCR analysis. SSR performed the in-silico analysis. HB, KA, AA, and SSR were involved in writing the first draft of the manuscript. HB and KA contributed equally and share first authorship. KHS was involved in manuscript editing and writing. SS conceived the project, planned, and supervised the research and was involved in data analysis, manuscript writing, and editing.

## Acknowledgments

We would like to acknowledge that the seeds were provided by the ICAR-Directorate of Rapeseed Mustard Research, Bharatpur, Rajasthan, India. We also acknowledge the Ministry of Education for the Prime Minister Research Fellowship to HB, the University Grants Commission (UGC) for the fellowship to KA, the Department of Biotechnology for the fellowship to AA, and the Department of Biotechnology Research Associate Fellowship to SSR. We acknowledge support from DBT for the Ramalingaswami Re-entry fellowship and IITGN start-up grants to SS.

