## Supplementary Data 1 for "Characterization of Self-Incompatibility Genes in *Brassica rapa* var. Toria and Yellow sarson"

### Supplemental Material - 1

#### Gene sequences of molecular players regulating SI:

##### ***SRK***

ATGAAAGGTGTACGAAACATCTATCACCATTCTTACACCTCCTTTTTGCTCGTCTTCGTTATCATGATTCTATTTCATCCTACTCTTT
CGATCTATATTAACGCTTTGTCGTCTACAGAATCTCTTACCATCTCAGGCAACAGAACACTTGTATCTCCCGGTGATGTCTTCGAGC
TCGGTTTCTTCAAGACCACTTCAAGTTCTCGTTGGTATCTCGGTATATGGTACAAGAAACTCTCCGAAAGAACCTACGTATGGGTT
GCCAACAGAGATAGCCCTCTCTCAAATGCCGTTGGAACCCTTAAAATCTCTAACATGAACCTGGTCCTCCTTGATCACTCTAATAA
ATCTGTTTGGTCGACGAATGCAACTAAAGGAAATGATAGATCTCCGGTGGTGGCAGAGCTTCTCGCTAACGGAAACTTCGTGATA
CGATACTCCAATAACAACGACGCAAGTAAATTTTTGTGGCAAAGTTTCGATTACCCTACAGATACTTTGCTTCCAGAGATGAAACT
AGGTTACGACCTCAAAAAAGGGCTGAACAGATACCTTACATCATGGAGAAATTCAGATGATCCGTCAAGCGGGGAAATCTCATAC
CAAATAGACAATCAAACGGGAATCCCTGAGTTCTATCTATTGCAAACCGGCATACGAGTGCATCGGAGCGGTCCATGGAATGGAG
TCCGATTTAGTGGCATACCAGGGGACCAAGAGTTAAGTTACATGGTGTACAATTTCACAGAGAATAGTGAAGAAGTCGCTTATAC
ATTCCGAGTGACCGACAACAGCATATACTCGATATTGAAAACAAGTTCCGAAGGGTTTTTGGAGCGACAGACGTGGACCCTGAAC
TCAATTACATGGACCTTGTTCTGGTATTTACCATTGGAAAACCAGTGCGATATGTACATGATTTGTGGGCGTTATGCTTACTGTGAT
GTGAACACATCACCGTTGTGTAACTGTATCCATGGGTTCATACCCTGGAATAAGCAGCAGTGGGAGATGATAAATCCGGCAGGTG
GGTGTATAAGGAGGACGCGGCTGAGCTGCAGTGGTGATGGTTTACCAGGATGAAGAAGATGAAGTTGCCAGAGACGAAGATGG
CGATTGTCGACAGGAGTATTGGTGTGAAAGAATGTGAGAAAAGGTGCCTTAGCAATTGTAATTGTACAGCTTTTGCAAATGCGGA
TATCCGGAATGGCGGGACGGGTTGTGTGATTTGGACAGGAGACCTCGAGGATCTTCGAAATTACTATGCTGACGGTCAAGATCTT
TATGTCAGATTGGCTGCCGCTGATCTTGTTAAAAAGAAAAACGCGAATTGGAAAATCATAAGTTTGATTGTTGGAGTTAGTGTTGT
TCTGCTTCTGCTTCTTCTGATCATGTTCTGCCTTTGGAAAAGGAAACAAAATCGAGCAAAAGCAATGGCAACATCTATTGTCAATC
AACAGAGAAACCAAAATGTACTTATGAACACGATGACACAATCAGACAAGAGACAGTTGTCTAGAGAGAACAAAGCTGATGAAG
TCCAACTTCCATTGATACAGGTGGAAGCTGTTGCCAAAGCCACCGCAAATTACTCCCATTGTAACGAACTTGGACGAGGTGGTTTC
GGTATTGTTTACAAGGGGATGCTTGATGGGCCAAATGTTGCGGTAAAAAGGCTATCAAAGACATCCCTTCAAGGCATTGATGAGT
TTATGAATGAGGTGAGATTGATCGCAAGGCTTCAGCATATAAACCTTGTCCGAAGTCTTGGCTGTTGCAATGAAGCGGACGAGAA
GATTCTGATATATGAGTATTTGGAAAAATCAAGCCTGGATTATTTTCTCTTTGGAACAAAACAAAGCTCTAACTTAAGTTGGAAGG
ACAGATTCGCCATTACCAATGGTGTTGCTCGAGGGCTTTTATATCTACATCATGACTCACGGTTTAGGATAATCCACCGGGATTTG
AAACCAGGTAACATTACGCTTGATAAATATAGGATCCCACGGATCTCGGATTTTGGGATGGCCAGAAACATAGCCAGGGATGAAA
CTCAAGTTAGGACAGACAATGCGGTCGGAACCAACGGCTACTGGTCTCCGGAAAACGCAATGTATGGGGTAATCTCGGGAAAAA
CAGATGTTTTCAGTTTTGGAGTCATAGTTCTTGAAATTGTTATTGGAAAAAGAAATAGAGGATTCTACCAGGTGAACCCTGAAAAC
AATCTTCCAAGCTATGCATGGACTCATTGGGCGGAGGGAAGAGCGCTAGAAATCGTAGATCCAGTCATCTTAGATTCATTGTCAT
CTCTGCCATCAACATTTAAACCAAAAGAAGTCCTAAAATGCATACAAATTGGTCTATTGTGTATTCAAGAACGTGCGGAGCACAG
ACCAACGATGTCGTCGGTGGTTTGGATGCTTGGAAGTGAAGCAACAGAGATTCCTCAGCCTAAACCGCCAGTTTATTGTCTCATAG
CAAGTTATTATGCAAATAATCCTTCCTCAAGTAAGCAATTCGATGACGATGAATCCTGGACAGTGAACAAGTACACCTGCTCAGT
CATCGATGCCCGGTAA

***FER1***

ATGAAGATAACTGAGGGACAATCACGTCTCTCCCTCCTCCTCCTCCTCCTTCTCTTATCCTTATCTTCATCAACCTCAGCTGCTGAC
TACACTCCCACTGACAAGATCCTCTTAAACTGTGGTGGCTCCTCTGATCGTACCGACACAGATAACCGGACATGGATCCCCGATGT
CAAATCCAAGTTCTTGTCTTCCTCCGGAGACTCCAAAACCTCTCCCGCCGCCACACAAGACCCTTCCGTCCCCGAGGTTCCTTACA
TGACAGCTAGAATCTTCCGGTCTCCCTTCACTTACTCTTTCCCCGTCGCTTCAGGTCGTAAGTTCGTGCGTCTCTACTTCCACCCCA
ACTCATACGACGGCCTCAACGCCACGACCTCCCTCTTCTCCGTCACCTTAGGCTCCTCCTACACTCTCCTCAAGAACTTCAGCGCT
GCTCAAACAGCTCAGGCCTTGTCTTACTCCTCCATCGTTAAAGAGTTTATCGTGAACGTGGAAGGTGGAGCTTCTTTGAACATAAC
GTTCACACCTGAATCAACACCAAAGGCTTATGCCTTTGTGAACGGTATTGAGGTGACTTCAATGCCTGATCTATACAGCAACACTG
ATGGGACTTTGTCCATCGTGGGATCTTCTACTGCGGTCGATATCGATAACAGCACTGCTCTTGAGAATGTTTACAGGCTTAACGTT
GGAGGGAATGATATCTCTCCTTCTGAAGATACAGGTCTTTACAGGTCATGGTACGACGACTCGCCTTACATTTTCACTGCGGGGAT
TGGAGTCGTTGAGACTGTTGATCCCAACATGACCATTAAGTATCCCACGGACACACCTACATACATTGCTCCTGTTGATGTTTACT
CAACTGCTAGGTCTATGACTCCCACAGCTCAAATCAACCTCAACTTCAACCTGACTTGGGTTTTCAGCATTGACTCTGGCTTCACTT
ATCTTGTTAGGCTTCATTTCTGCGAGGTTCTTCCCGACATCACTAAGATTAACCAGCGTGTGTTTACAATCTACCTCAACAACCAA
ACAGCTGAGTCTGAAGCTGATGTTGCTGGCTGGACGGGTGGTAATGGGATTCCTATATATAAAGACTACGTTGTGAATCCTCCTGA
TGGTAAGGGACAGCAAGATCTTTGGCTTGCTCTTCATCCAAACACGAGGGGCAAGCCGGAGTACTACGATGCTATTCTTAATGGA
GTTGAGATTTTCAAGATGAATGGTTCTGATGGTAATCTTGCTGGTCCTAATCCTATACCTGGTCCGCAAGTGACTGCAGATCCATC
CAGAGTGTTACGCCCTCGCACTGGTTCATCTAAGAGCCATACAGCTATTGTTGCAGGTGTAATCAGTGGTGCAGTTGTTTTAGGTC
TTATTGTTGGTTTATGTGTAATGGTTGCTTACCGTAGACGTAAGGCTGGTGAATACCAGCCTGCAAGTGATGCAACATCAGGGTGG
CTTCCACTGTCTTTGTATGGAAACTCACATTCTGGTGGCTCGGGTAAGACAAACACTACAGGAAGCTACGCCTCGTCCCTTCCTTC
AAACCTGTGTCGTCACTTCTCCTTTGCTGAGATCAAAGCAGCTACTAAGAACTTTGATGAGTCTCGAGTGCTCGGTGTTGGAGGTT
TTGGTAAGGTGTACAGAGGAGAGATTGATGGTGGAACTACAAAGGTAGCCATCAAGAGAGGCAACCCTATGTCTGAGCAAGGTG
TGCACGAGTTTCAGACAGAGATTGAGATGCTTTCGAAGCTTAGACACCGTCACCTTGTGTCTTTGATTGGATACTGTGAAGAGAAC
TGCGAGATGATACTTGTGTATGATTACATGGCTCATGGGACAATGAGAGAGCATCTCTACAAGACTCAGAACTCTCCTCTTCCTTG
GAAGCAACGTCTTGAGATATGCATTGGGGCAGCAAGAGGGTTGCATTATCTACACACCGGTGCGAAACACACGATCATCCACAGA
GATGTGAAGACGACAAACATTCTGTTGGATGAGAAGTGGGTGGCTAAGGTCTCTGACTTCGGTCTGTCAAAGACTGGTCCTACAC
TTGACCACACACATGTTAGCACGGTGGTGAAAGGAAGCTTCGGTTATCTCGACCCAGAGTACTTCAGACGTCAGCAACTGACTGA
TAAATCAGATGTCTACTCCTTTGGTGTCGTTCTTTTCGAAGCTCTATGCGCACGGCCTGCCTTGAACCCGACGCTAGCAAAAGAAC
AAGTGAGCTTAGCTGAGTGGGCACCATACTGCTACAAGAAAGGCATGCTTGACCAGATCGTTGATCCGCATCTCAAGGGCAAGAT
CACACCGGAATGCTTCAAGAAGTTTGCTGAAACCGCGATGAAGTGTGTACTAGACCAGGGCATTGAGAGACCGTCGATGGGAGA
TGTTCTCTGGAACTTAGAGTTCGCGTTGCAGCTTCAGGAAAGCGCTGAGGAGAGCGGGAAAGGGATATGCAGTGAGATGGACAT
GGGTGAGATTAAGTACGATGATGATAACTGTAAAGGGAAGAGCAACAACGACAAGGGCTCTGATGTGTATGAAGGGAATGTTAG
TGACTCGAGGAGCAGTGGTATAGACATGAGTATTGGTGGTAGGAGTTTGGTCAGTGAAGATTCAGATGGACTCACTCCAAGTGCT
GTGTTTTCTCAGATCATGAATCCTAAGGGACGTTAG

***MLPK***

ATGGGGATTTGCTTGAGTGCTCAGATTAAAGCTGAGAGTCCAAGTAACACAGGTGCGAGTCCGAAGTATATGAGCTCAGAGGCAA
ATGATACACAGAGCATGGGAAGCAAAGGCTCTTCTGTGTCGATCAGAACAAACCCTCGAACCGAAGGAGAGATCTTGCAATCTCC
AAACCTCAAAAGTTTTAGCTTCGCTGAGGTCAAATCAGCAACTAGGAATTTCAGACCAGACAGTGTTCTTGGTGAAGGTGGATTC
GGTTGTGTGTTTAAAGGATGGATTGATGAGCAATCTCTCACTGCGTCTAAACCGGGAACCGGTATGGTTATTGCTGTCAAAAGACT
TAACCAAGATGGTTGGCAAGGTCATCAAGAATGGCTGGCGGAAGTGGATTACTTGGGGAAGTTCTCTCATCCTAATCTTGTGAAA
CTCATCGGTTATTGTTTAGAGGATGAGCAACGTCTTCTTGTGTATGAGTTCATGCCACGTGGAAGCTTAGAGAATCATTTATTCAG
AAGAGGTTCTTACTTTGAACCATTATCTTGGACTCTCAGATTGAAAGTTGCACTTGGCGCTGCAAAAGGCCTAGCTTTTCTTCACA
ACGCGGAGACTCAAGTCATATACCGGGACTTCAAAACTTCTAACATACTTATTGATTCGGACTACAACCCCAAGCTTTCTGATTTT
GGGTTGGCTAAAGACGGTCCAACAGGTGATAAAAGCCATGTCTCCACAAGAATCATGGGTACTTATGGATACGCAGCTCCTGAGT
ATCTTATGACAGGTCATTTAACAACCAAGAGTGATGTCTATAGCTACGGTGTTGTGCTTTTGGAGATACTCTCTGGACGTAGAGTT
GTAGACAAGAACCGTCCACCGGGAGAGCAAAAACTGGTGGATTGGGCAAAACCGTTGCTTGCAAACAAGAGGAAGATCTTTAGA
GTTATCGATAACCGTCTACAAGATCAGTACTCAATGGAAGAAGCGTGTAAAGTAGCTACTCTAGCGCTGAGATGCCTGACGACAG
AGATAAAGCTGAGACCAAACATGACTGAGGTTGTTGCTCACCTCGAACACATACAAACTTTGCATGAAACAGGAGGAGGAAGAA
ACATTGATAAGTTGGAGAGGAGAACGCGTAGGAGAAGTGATAGTGTTGTGGTGAGCCAAAAACCAAATGCTGGTTTTGCTAGAC
AAAGTGCTGTGGGTGGAATAGCAGCTGCGTATCCACGTCCCTCTGCTTCGCCTCTGTTTGTCTGA

***ARC1***

ATGGCCACTGATTCAGCAATGTTCGCATCCTCACGTCGGAGGCAATCTCCGTCGCTTGAGGCGTTTCTATCACCCGTTGATCTCTCCGACGTCCCTCTCCTCCAAACACTATCTTCCATCTCATCAGAGATCGTCTCCTGCTTCAGCAACGCACGTTTCTCCTTCCAACGTAGAAACACCCGTTCCCTGATACGTAAAGTCCAAGTCTTCGCCGTCTTACTCCAACACCTCGCACCCGAGTCAAGCTTGGATCCCACGGCGGTGCTCTGCTTCAAGGAGCTCTATCTCCTCCTCCACCACTCCAAGTTCCTCCTCCGCTACTGCGCTCACTCCTCCAAGCTATGGCTCTTGCTTCAAAGCCCCTCGCTCTCGAGCTTCTTCCATGATCTGAGTAAAGACTATTCCACCCTCTTAGATGTCCTCCTCCCTGCTGAGAGTCTCTGCCTAAACGACGACGTTAGAGAGCAAGTCCAGCTCTTGCACATGCAGCACTACATTGACGATAACAGCGACGAGACGCTGCGTAACAAACTCTATTCGTTTCTAGACGAGTTCGAGAACGGGAGTGTACCAAACTCTGAAGAGCTACGCTTCTTCTTCTTTGAGAAACTCGCTATTAAAGATCCAACAAGTTACAGAGAAGAGATCGAGTTTCTTGAAGAGCAGATCAAAAGCCACGGGTGTGACTTAGACCCTACGAGGTCAGTGATCAACGGGTTTATAGATATCACACGGTACGTTATGTTTCTCTTATTCAAGATTGAAGATGGTAACGAGATTAACAAACAGAAGAAACGTTTGATCTCTGAGGAGATTGAGAACACGTTTACAACAACGCTTCCAAAGGATTTCATCTGCTCCATCTCTCTCAACCTCATGAACGATCCTGTGATCATCTCCACGGGACAGACTTACGATAGAACCTCCATCGCTAGGTGGATTCATCAAGAAGGTCGCTCTACTTGTCCCAAAACAGGACAGAAGCTCGTGGACTTGAGTTTCGTTCCCAACCTAGCTTTGAGACACTTGACAAAGCTTTGGTGCCAAGTCACTGGTCTGTCTCATGACTCGCCTAAAGAGTCTCTCCCAAAGGTGTTTCAAACAAGAGCTTCCACGGAAGCAAACAAAGCAACGTTATCGATTCTTGTACAGAACCTAGCACACGGCTCAGAGTTGGCTGCAGGAGAGATCCGTGTTCTCACTAGAACAGTAACGGAAACGCGTACGTTGATCGTGGAAGCAGGTGCGATCCCGTATCTGCGTAGTCTTCTCAAATCCCAAAACGCTGTTGCGCAGGAGAACGCAGTTGCATCGATCTTTAACTTATCTATAGACGAAGAAAACAGGAGTCTGATCATGGAGGAACACTCTTGTCTCGAGCCGATGATGAGCGTTCTCGTCTCTGGTCTTACGATGAGAGCTAAGGAGATAGCAGCAGCCACGTTGCACACTCTTTCCAGCGTACATGATTACAAGAAAACGATCGCTAACGCTGATGGATGCATCGAGGCGCTTGCACTGGTGCTGCGAAACGGAACCGTGAGAGGGAAGAAAGATGCTGTCTACGCTTTGCATAGCTTATGGCTGCATCCGGATAACTACAGCTTGATGGTAAAAAGGGGAGGAGTGTCTGCTCTCGTTGGAGCTTTAGGGGAAGAGGCTGTGGCGGAGAAAGTTGCGTGGGTGTTGGGTGTGATGGCTACTGAGACTTTAGGAGCTGAGAGTATAGGGAGAGAGGAATCAGTTGTGACGGGGCTCATGGAACTAATGAGATGTGGAAGACCTAGAGGCAAAGAAAAAGCTATTGCGACTTTGTTACAACTCTGCACAGCAGGTGGAGCGGTTGTGACGGAGAAGGTTGTGAAAACACCCGCTCTTGCGGTCTTGACGCGTAAGCTTTTGCTCACGGGTACAGACCGAGCTAAGAGGAAAGCGGTTTCACTCTCTAAGGTATGTAAGGGGTGCGACCAGAAAACACAGAGATAA

**Supplementary Figure legends**

**Supplementary Figure S1.** Germination assay of F1 seeds obtained from various crosses between *B.rapa* var. toria and yellow sarson.

**Supplementary Figure S2.** PCR optimization, Colony PCR, and Restriction Digestion-based analysis of various genes regulating self-incompatibility in *B.rapa* var. toria.

(A) Gel images showing PCR products of various genes under study. Note: The size of the PCR products is indicated.

(B) Gel images showing colony PCR of various DH5α colonies transformed by TOPO cloning-based products of various genes under study. Note: The size of the PCR band is indicated, and red asterisks indicate positive colonies.

(C) Gel images showing Restriction digestion-based products of various TOPO-gene constructs using EcoR1. Note: The size of the Restriction digestion products is indicated.

**Supplementary Figure S3.** Graphical representation of proteins (A) SRK, (B**)** FER1, (C) MLPK, and (D) ARC1 and their conserved domains.

**Supplementary Figure S4.** Secondary structure prediction of (A) SRK; (B) FER1; (C) MLPK, and (D) ARC1 proteins from *Brassica rapa* var. toria predicted through PSIPRED.

**Supplementary Figure S5.** Ramachandran plots of AlphaFold3 predicted 3-D structures (A) SRK; (B) FER1; (C) MLPK; and (D) ARC1, indicating the accuracy and reliability of the predicted structures.

**Supplementary Figure S6.** Illustration of the top-dripping method for *in vitro* ODN treatment.

**Supplementary Figure S7.** Functional characterization of genes during compatibility response.

(A-D) Representative aniline blue images of pollinated stigmas treated with Mock, Sense, and Anti-Sense ODNs of (A) *SRK*, (B) *FER1*, (C) *MLPK*, and (D) *ARC1*, showing pollen attachment and tube formation. Note: Aniline blue (left); Red channel (middle); Bright field (right). Scale bars: 300 µm

(E-H) Graph indicating the number of pollen tubes per stigma observed in (Figure 6A-D) & (Supplementary Figure S7A-D) treated with Mock, Sense, and Anti-Sense ODNs of (E) *SRK*, (F)

*FER1*, (G) *MLPK*, and (H) *ARC1*. Note: Dots represent individual data points. One way ANOVA, *$P < 0.05$; n = 9, ns= not significant

**Supplementary Figure S8.** Optimization of Nitro Blue Tetrazolium (NBT) assay.

(A) Representative images of stigma stained with NBT at unpollinated or 15, 30, and 60 MAP during SI response.

(B) Graph representing the normalized intensity of ROS for various time points after the self-incompatibility response. Note: One-way ANOVA, *$p<0.05$, n=15

**Supplementary Figure S9.** Gel image for the *SRK*, *FER1*, *MLPK*, and *ARC1* treated with ODNs - Mock, Sense, and Anti-Sense. Note: *BrActin 7* (NCBI accession: KU851921) was used as an internal control

#### Supplementary Tables

**Supplementary Table S1.** List of primers used for cloning in this study.

| S.No. | Gene | Primers | Template Size | NCBI Accession |
|---|---|---|---|---|
| 1 | *SRK* | FP: ATGAAAGGTGTACGAAACATC | 2580 bp | OR887605 |
| | | RP: TTACCGGGCATCGATGACTGA | | |
| 2 | *FER1* | FP: ATGAAGATAACTGAGG | 2616 bp | PX355005 |
| | | RP: CTAACGTCCCTTAGGA | | |
| 3 | *MLPK* | FP: ATGGGGATTTGCTTGAGTGC | 1257 bp | PV420907 |
| | | RP: TCAGACAAACAGAGGCGAAG | | |
| 4 | *ARC1* | FP: ATGGCCACTGATTCAGCAATG | 1983 bp | PX058862 |
| | | RP: TTATCTCTGTGTTTTCTGGTC | | |

**Supplementary Table S2.** PCR conditions for amplification of genes used in this study.

<table>
<tr><th>Gene</th><th>Initial denaturation</th><th>Denaturation</th><th>Annealing</th><th>Extension</th><th>Final extension</th></tr>
<tr><td>No. of cycles</td><td>1x</td><td colspan="3">35 Cycles</td><td>1x</td></tr>
<tr><td>SRK</td><td rowspan="4">98ºC; 2 min</td><td rowspan="4">98ºC; 30 sec</td><td>61ºC; 30 sec</td><td>72 ºC; 1 min and 15 secs</td><td rowspan="4">72 ºC; 10 min</td></tr>
<tr><td>FER1</td><td>49ºC; 30 sec</td><td>72 ºC; 1 min and 30 secs</td></tr>
<tr><td>MLPK</td><td>63ºC; 30 sec</td><td>72 ºC; 45 secs</td></tr>
<tr><td>ARC1</td><td>59ºC; 30 sec</td><td>72 ºC;1 min</td></tr>
</table>

**Supplementary Table S3.** List of the *SRK* gene sequences retrieved from the NCBI database for phylogenetic analysis.

| S.No. | Accession number | Gene Description |
|---|---|---|
| 1 | OR887605 | *Brassica rapa* cultivar toria S receptor kinase |
| 2 | AB219163.1 | *Brassica rapa* BrSRK-54f S receptor kinase |
| 3 | AB032474.1 | *Brassica oleracea* SRK60 S60 S-locus receptor kinase |
| 4 | AB298890.1 | *Brassica oleracea* SRK-4 mRNA S-locus receptor kinase |
| 5 | AB054061.1 | *Brassica rapa* SRK22 mRNA S locus receptor kinase |
| 6 | AB270777.1 | *Brassica napus* BnSRK-3 pseudogene S receptor kinase |
| 7 | D38564.2 | *Brassica campestris* SRK12 receptor protein kinase |
| 8 | D38563.1 | *Brassica campestris* SRK8 receptor protein kinase |
| 9 | EU180597.1 | *Brassica oleracea* var. acephala SRK13-b receptor kinase |
| 10 | AB298885.1 | *Brassica rapa* SRK-55 S-locus receptor kinase |
| 11 | AB298887.1 | *Brassica rapa* SRK-61 S-locus receptor kinase |
| 12 | AB270775.1 | *Brassica rapa* BrSRK-21 S receptor kinase |
| 13 | M97667.1 | *Brassica napus* ssp. oleifera serine/threonine kinase receptor |
| 14 | AB032473.1 | *Brassica oleracea* SRK18 S-locus receptor kinase |
| 15 | M76647.1 | *Brassica oleracea* SKR6 receptor protein kinase |
| 16 | AB270776.1 | *Brassica napus* BnSRK-2 pseudogene S receptor kinase |
| 17 | AB298875.1 | *Brassica rapa* SRK-25 S-locus receptor kinase |
| 18 | AB270767.1 | *Brassica napus* BnSRK-1 S receptor kinase |
| 19 | AB013720.1 | *Brassica oleracea* SRK23Bol S receptor kinase |
| 20 | AB298891.1 | *Brassica oleracea* SRK-14 S-locus receptor kinase |
| 21 | AB298884.1 | *Brassica rapa* SRK-53 S-locus receptor kinase |
| 22 | LC556298.1 | *Brassica rapa* BrSRK-9 S-locus receptor kinase |
| 23 | AB298902.1 | *Brassica oleracea* SRK-61 S-locus receptor kinase |
| 24 | U00443.1 | *Brassica napus* cultivar T2 S-receptor kinase |
| 25 | AB298901.1 | *Brassica oleracea* SRK-52 S-locus receptor kinase |
| 26 | AB298886.1 | *Brassica rapa* SRK-56 S-locus receptor kinase |

| | | |
|---|---|---|
| 27 | AB012106.1 | *Brassica rapa* SRK45 S-locus receptor kinase |
| 28 | AB298905.1 | *Brassica oleracea* SRK-68 S-locus receptor kinase |
| 29 | AB052756.1 | *Arabidopsis lyrata* SRKb S-locus receptor kinase |

**Supplementary Table S4.** List of the *FER1* gene sequences retrieved from the NCBI database for phylogenetic analysis.

| **S.No.** | **Accession number** | **Gene Description** |
|---|---|---|
| 1 | PX355005.1 | *Brassica rapa* cultivar toria FER1 |
| 2 | EF681131.1 | *Brassica oleracea* FER1 |
| 3 | XM_048743008.1 | *Brassica napus* FER1 |
| 4 | EF681137.1 | *Arabidopsis thaliana* FER1 |

**Supplementary Table S5.** List of the *MLPK* gene sequences retrieved from the NCBI database for phylogenetic analysis.

| S.No. | Accession number | Gene Description |
|---|---|---|
| 1 | PV420907.1 | *Brassica rapa* M locus protein kinase |
| 2 | XM_013888729.3 | *Brassica napus* probable serine/threonine-protein kinase (predicted) |
| 3 | KC576522.1 | *Brassica rapa* subsp. chinensis M locus protein kinase |
| 4 | XM_013780614.1 | *Brassica oleracea* var. oleracea protein kinase chloroplastic (predicted) |
| 5 | NM_001036363.2 | *Arabidopsis thaliana* protein kinase 1B PK1B |

**Supplementary Table S6.** List of the *ARC1* gene sequences retrieved from the NCBI database for phylogenetic analysis.

| **S.No.** | **Accession number** | **Gene Description** |
|---|---|---|
| 1 | PX058862.1 | *Brassica rapa* cultivar toria armadillo repeat containing 1 |
| 2 | PX058861.1 | *Brassica rapa* cultivar yellow sarson armadillo repeat containing 1 |
| 3 | KC576518.1 | *Brassica rapa* subsp. chinensis armadillo repeat containing 1 |
| 4 | AF024625.1 | *Brassica napus* arm repeat containing 1 |
| 5 | EU344909.1 | *Brassica oleracea* var. acephala arm repeat containing protein 1 |

**Supplementary Table S7.** List of conserved domains and their location in various proteins under study.

| Protein | Name of domains | Accession | Description | Internal | E-value |
|---|---|---|---|---|---|
| SRK | STKc_IRAK | cd14066 | Catalytic domain of the Serine/Threonine kinases, | 537-808 | $6.54 \times 10^{-79}$ |
| | B_lectin | pfam01453 | D-mannose binding lectin | 83-189 | $2.94 \times 10^{-47}$ |
| | S_locus_glycop | pfam00954 | S-locus glycoprotein domain | 223-330 | $1.39 \times 10^{-40}$ |
| | PAN_2 | pfam08276 | PAN-like domain | 354-418 | $3.20 \times 10^{-29}$ |
| | DUF3403 | pfam11883 | Domain of unknown function (DUF3403) | 810-859 | $5.75 \times 10^{-10}$ |
| | DUF3660 | pfam12398 | Receptor serine/threonine kinase | 483-520 | $1.98 \times 10^{-7}$ |
| FER1 | STKc_IRAK | cd14066 | Catalytic domain of the Serine/Threonine kinases, Interleukin-1 Receptor Associated Kinases and related STKs | 541-807 | $1.21 \times 10^{-88}$ |
| | PK_Tyr_Ser-Thr | pfam07714 | Protein tyrosine and serine/threonine kinase | 539-735 | $2.09 \times 10^{-49}$ |
| | STYKc | smart00221 | Protein kinase; unclassified specificity | 539-735 | $5.32 \times 10^{-49}$ |
| | SPS1 | COG0515 | Serine/threonine protein kinase Signal | 539-735 | $6.49 \times 10^{-46}$ |

| | | | | | |
|---|---|---|---|---|---|
| | | | transduction mechanisms | | |
| | Malectin_like | pfam12819 | Malectin-like domain | 38-408 | $1.81 \times 10^{-40}$ |
| MLPK | STKc_IRAK | cd14066 | Catalytic domain of the Serine/Threonine kinases, Interleukin-1 Receptor Associated Kinases and related STKs | 80-359 | $1.08 \times 10^{-96}$ |
| | TyrKc | smart00219 | Tyrosine kinase, catalytic domain | 77-356 | $7.89 \times 10^{-54}$ |
| | PK_Tyr_Ser-Thr | pfam07714 | Protein tyrosine and serine/threonine kinase | 79-356 | $2.57 \times 10^{-53}$ |
| | SPS1 | COG0515 | Serine/threonine protein kinase [Signal transduction mechanisms] | 73-416 | $2.51 \times 10^{-49}$ |
| | PLN00113 | PLN00113 | leucine-rich repeat receptor-like protein kinase | 50-357 | $4.93 \times 10^{-24}$ |
| | knB_PASTA_kin | NF033483 | Stk1 family PASTA domain-containing Ser/Thr kinase | 178-283 | $2.59 \times 10^{-11}$ |
| ARC1 | RING-Ubox_PUB | cd16664 | U-box domain, a modified RING finger, found in *Arabidopsis* plant U-box proteins (AtPUB) and similar proteins | 281-332 | $5.00 \times 10^{-24}$ |

| | | | | | |
|---|---|---|---|---|---|
| | U-box | smart00504 | Modified RING finger domain | 283-347 | $9.36 \times 10^{-24}$ |
| | U-box | pfam04564 | U-box domain | 281-341 | $3.24 \times 10^{-14}$ |
| | PLN03200 | PLN03200 | Cellulose synthase-interactive protein | 374-624 | $9.75 \times 10^{-8}$ |
| | Arm | pfam00514 | Armadillo/beta-catenin-like repeat | 406-443 | $2.04 \times 10^{-4}$ |

**Supplementary Table S8.** List of oligonucleotides used in this study.

| **S.No.** | **Gene** | **Sense/Antisense** | **Sequence** |
|---|---|---|---|
| 1 | *SRK* | Sense | T*A*G*CCCTCTCTCAAATG*C*C*G |
| | | Antisense | C*G*G*CATTTGAGAGAGGG*C*T*A |
| 2 | *FER1* | Sense | A*T*G*AAGATAACTGAGGG*A*C*A |
| | | Antisense | T*G*T*CCCTCAGTTATCTT*C*A*T |
| 3 | *MLPK* | Sense | A*T*G*GGAAGCAAAGGCTC*T*T*C |
| | | Antisense | G*A*A*GAGCCTTTGCTTCC*C*A*T |
| 4 | *ARC1* | Sense | G*A*C*GTCCCTCTCCTCCA*A*A*C |
| | | Antisense | G*T*T*TGGAGGAGAGGGAC*G*T*C |

Note- Asterisks (*) indicate phosphorothioate modifications at the three terminal bases on both the 5’ and 3’ ends.

**Supplementary Table S9.** List of primers used to perform RT-PCR in this study.

| S.No. | Gene | Primers | Amplicon size (bp) |
|---|---|---|---|
| 1 | *SRK* | FP: CAGACGTGGACCCTGAACTC | 209 |
| | | RP: CGCGTCCTCCTTATACACCC | |
| 2 | *FER1* | FP: ACCTACATACATTGCTCCTGTTGA | 187 |
| | | RP: TGTAAACACACGCTGGTTAATCTT | |
| 3 | *MLPK* | FP: GGTTGGCTAAAGACGGTCCA | 223 |
| | | RP: GGTTTTGCCCAATCCACCAG | |
| 4 | *ARC1* | FP: AGTCAAGCTTGGATCCCACG | 219 |
| | | RP: CGTCGTCGTTTAGGCAGAGA | |
| 5 | *BrActin 7* | FP: TGGTTCGACCATGTTCCCTG | 213 |
| | | RP: CTGTGGACGATGGATGGACC | |

**Supplementary Figures**

**Supplementary Figure S1**

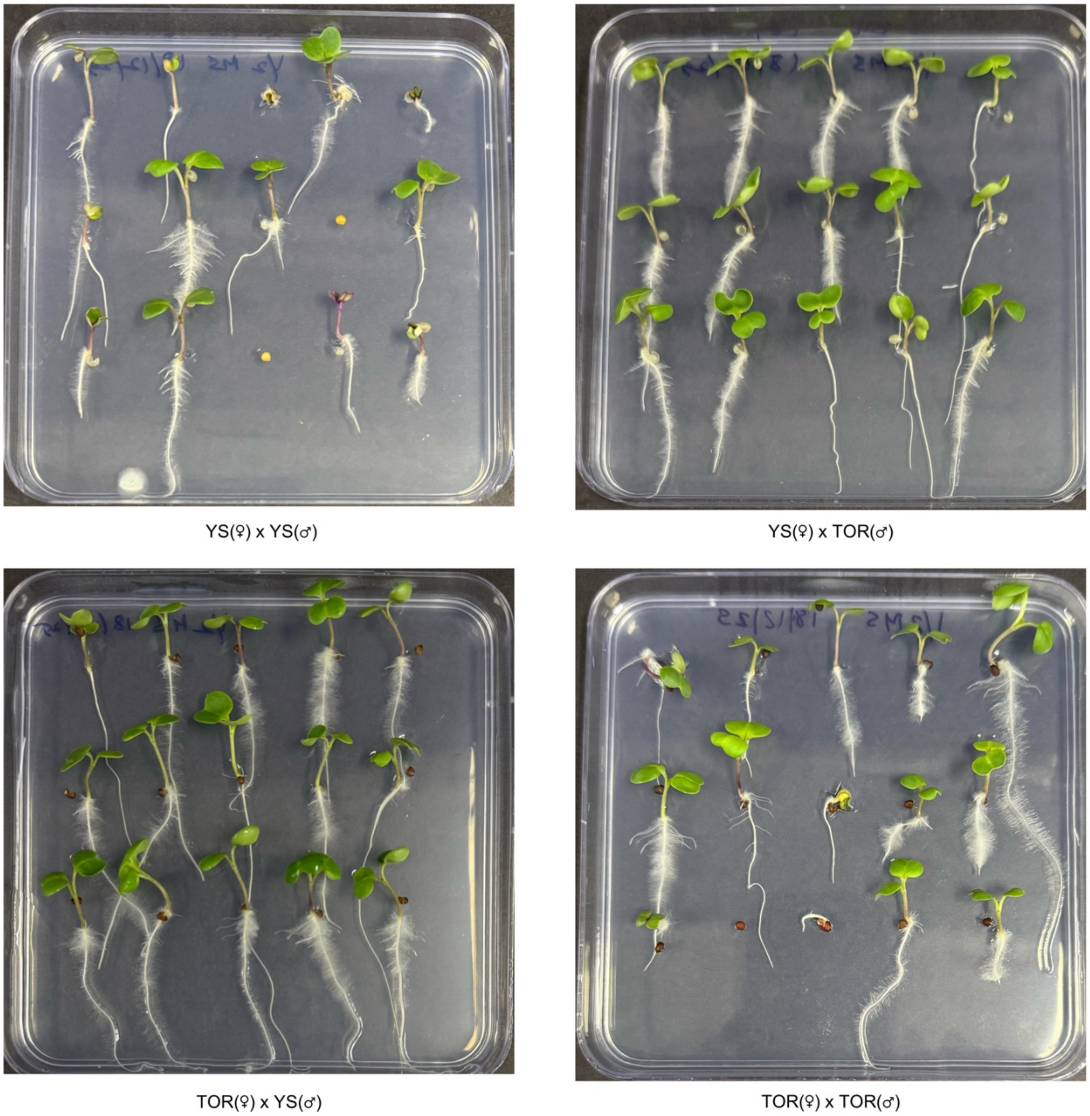

**Supplementary Figure S2**

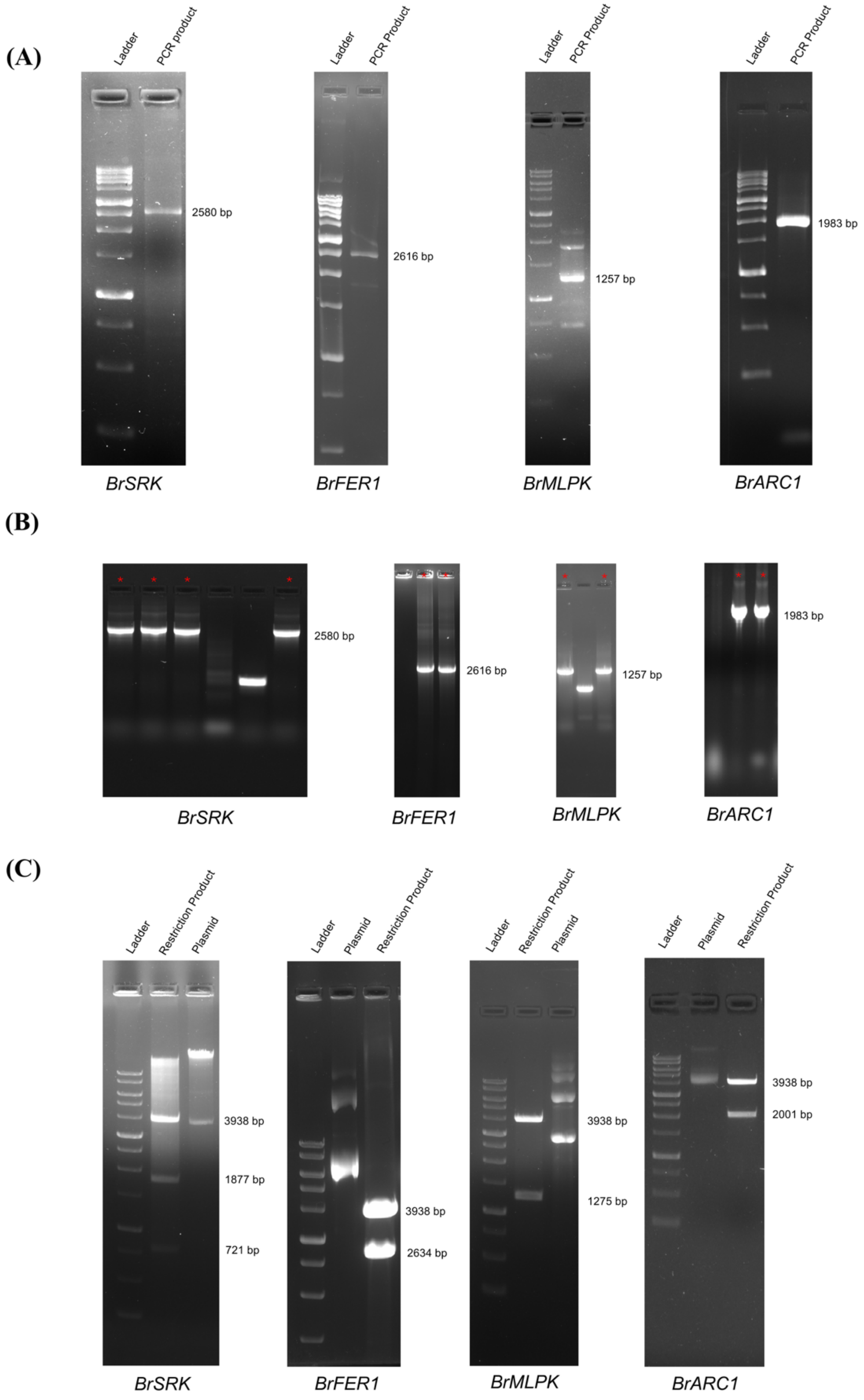

358 **Supplementary Figure S3**

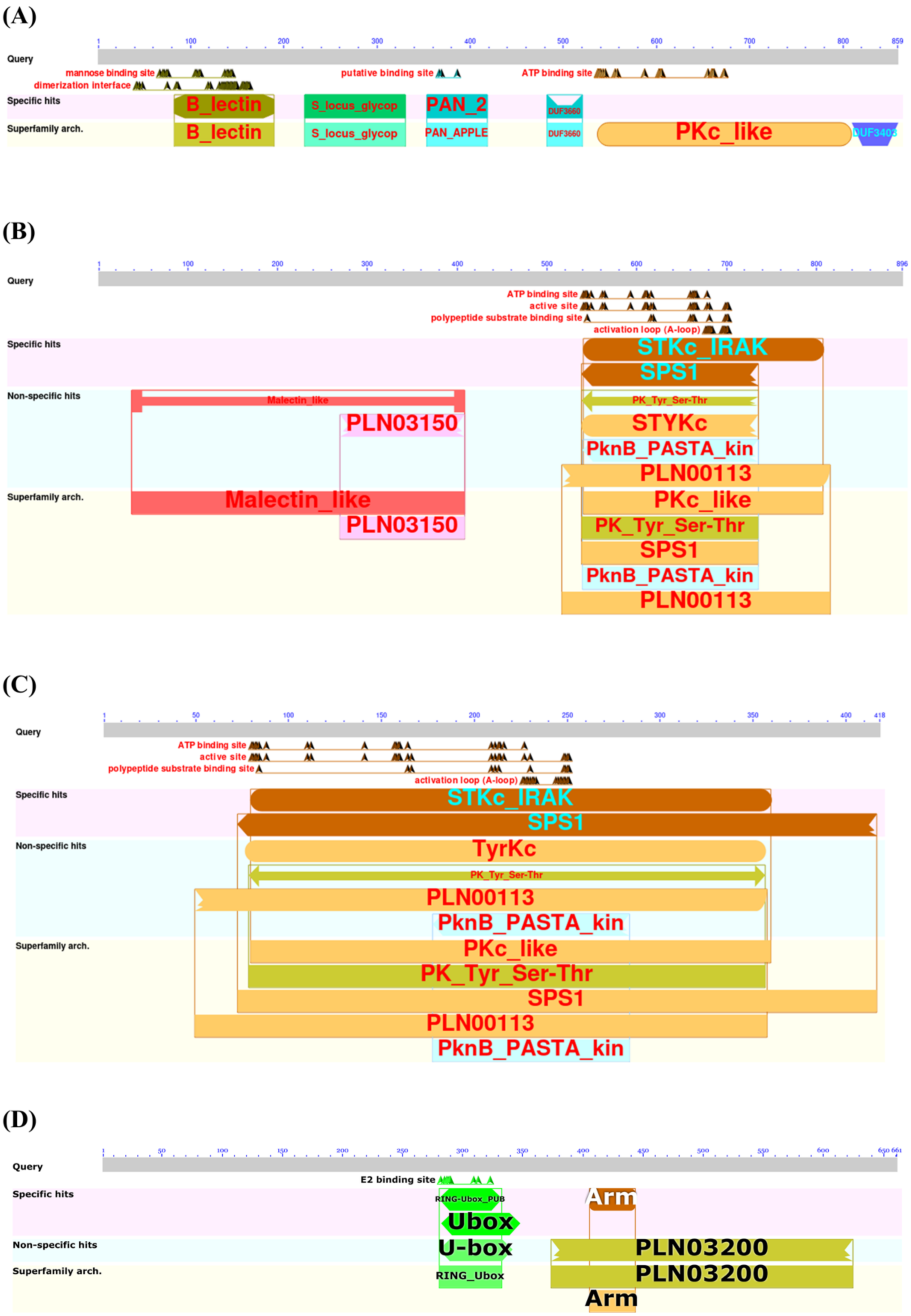


359

#### Supplementary Figure S4

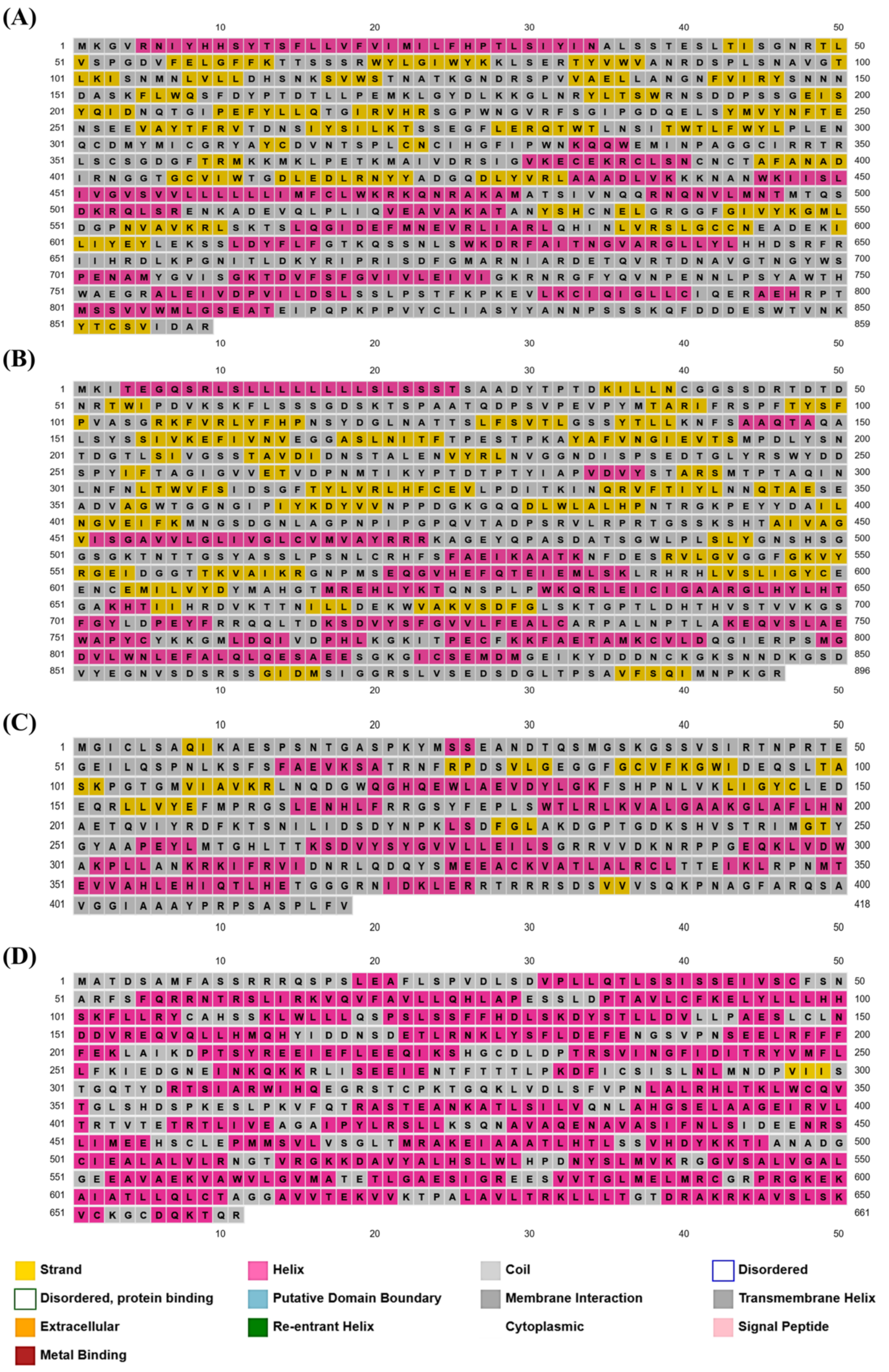

#### Supplementary Figure S5

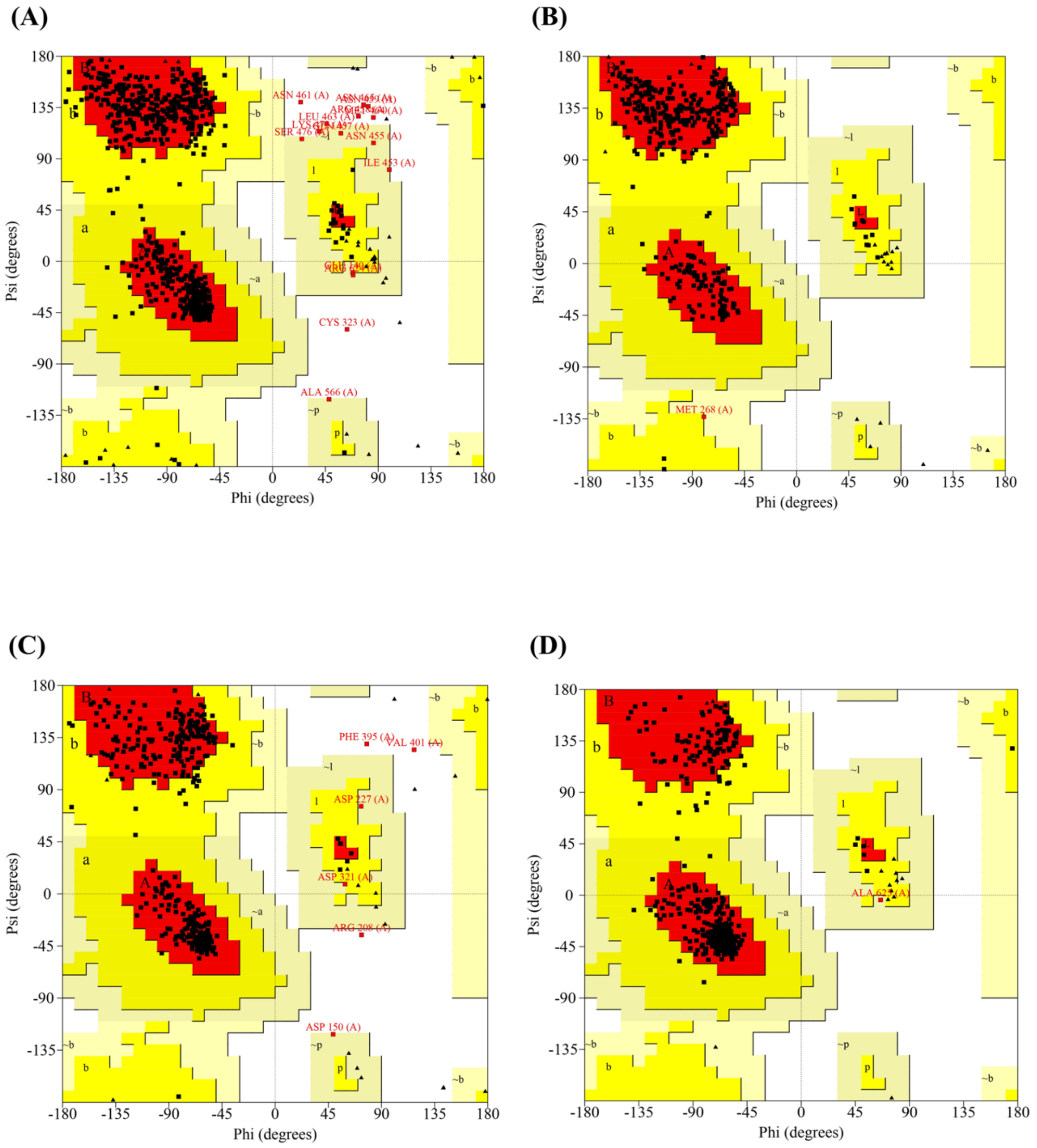

**Supplementary Figure S6**

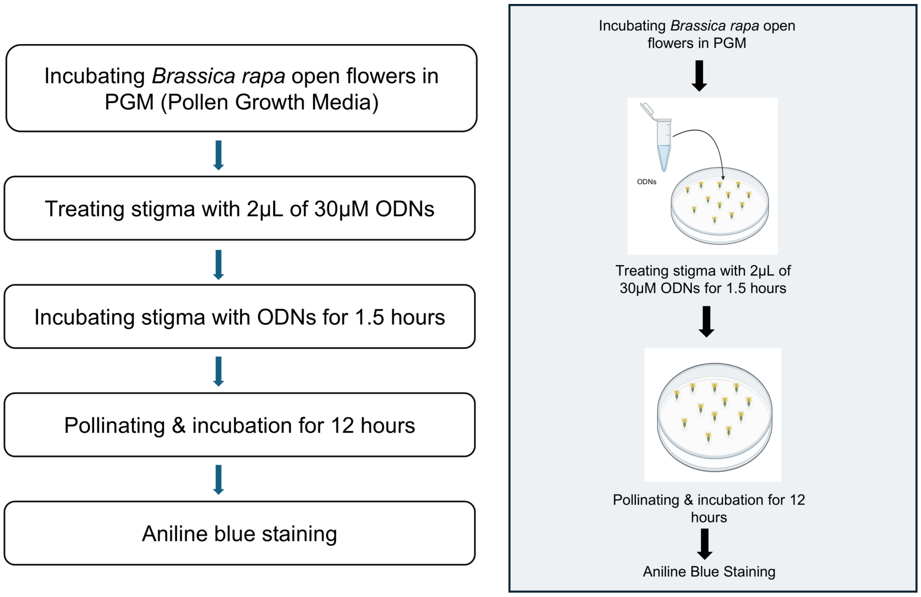

#### Supplementary Figure S7

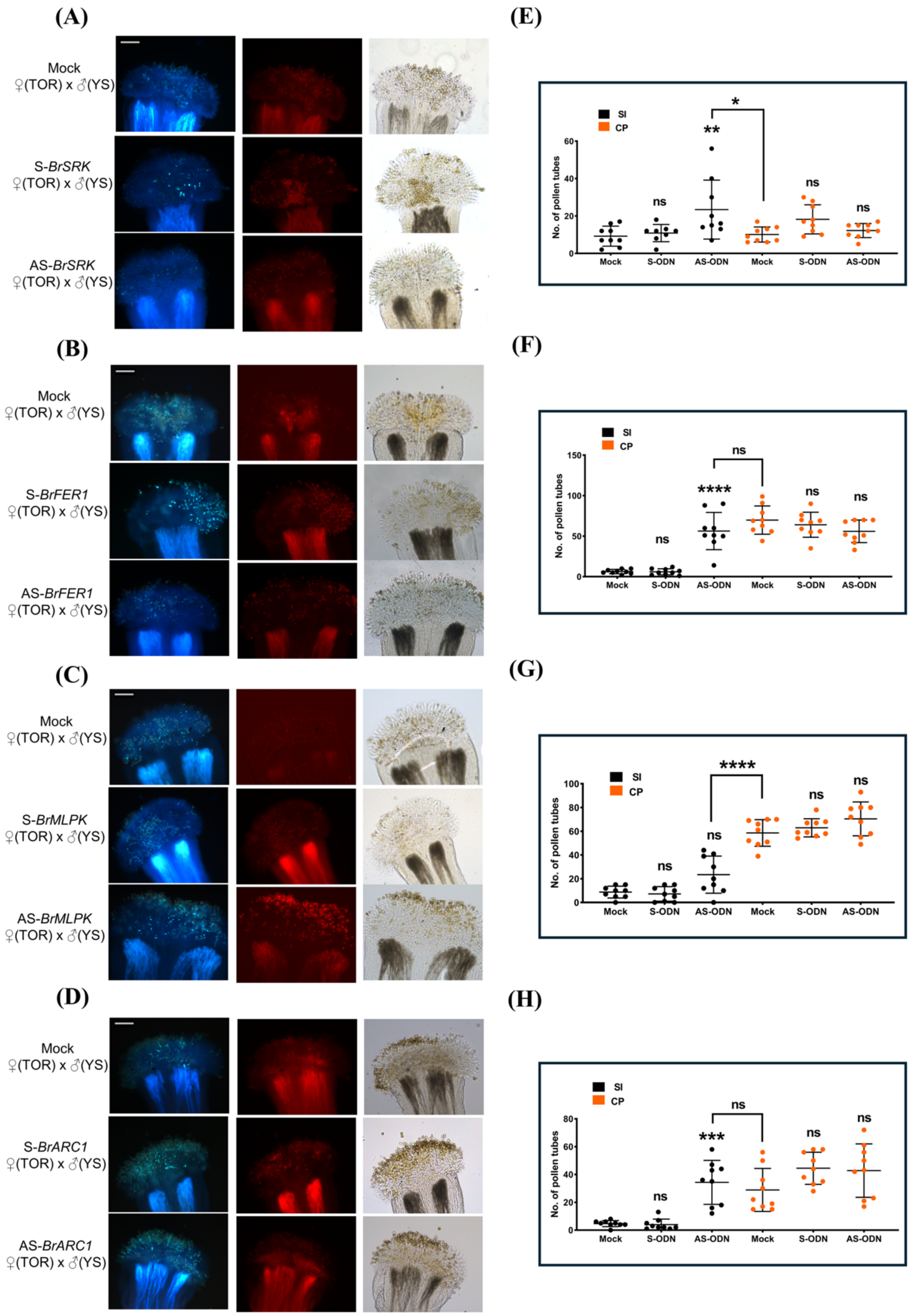

**Supplementary Figure S8**

**(A)**

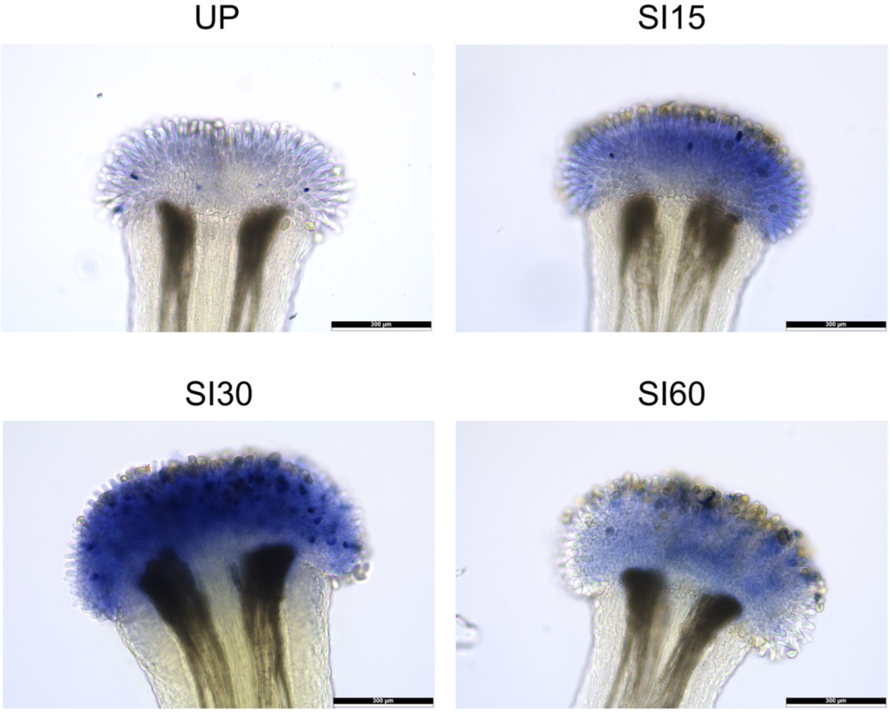


**(B)**

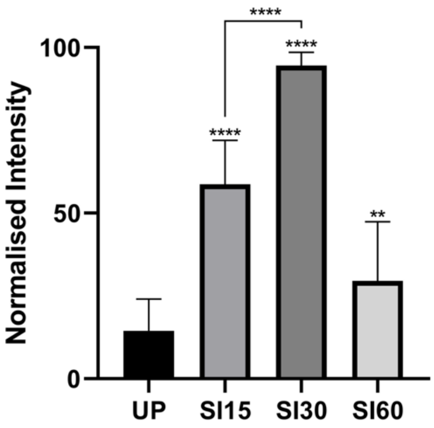

**Supplementary Figure S9**

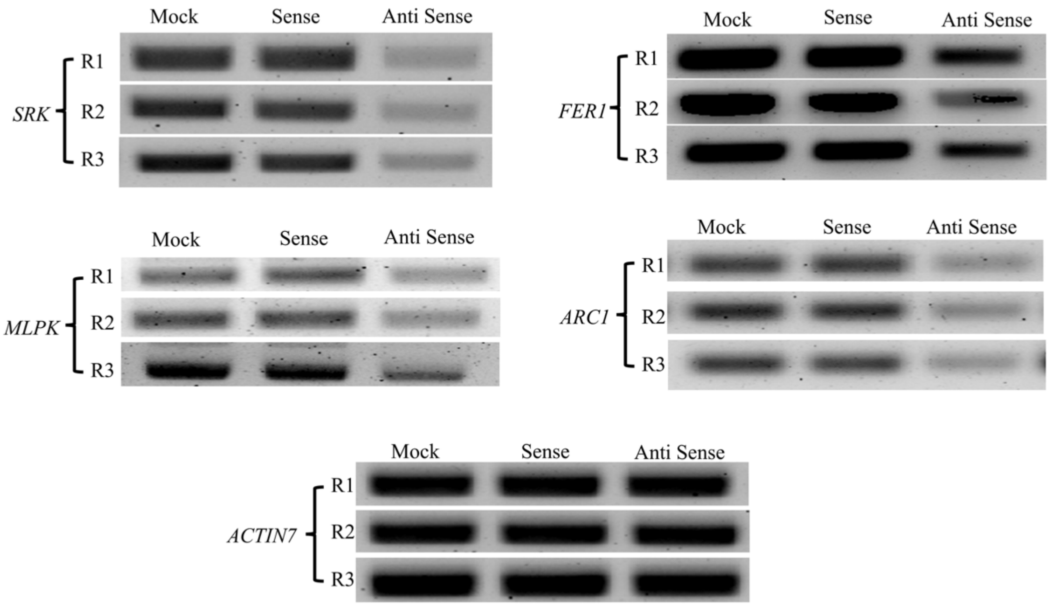
