## Supplementary Data 3 for "Characterization of Self-Incompatibility Genes in *Brassica rapa* var. Toria and Yellow sarson"

**Supplemental Material - 3**

**Multiple sequence alignment of gene sequences of *SRK* selected for phylogenetic study**

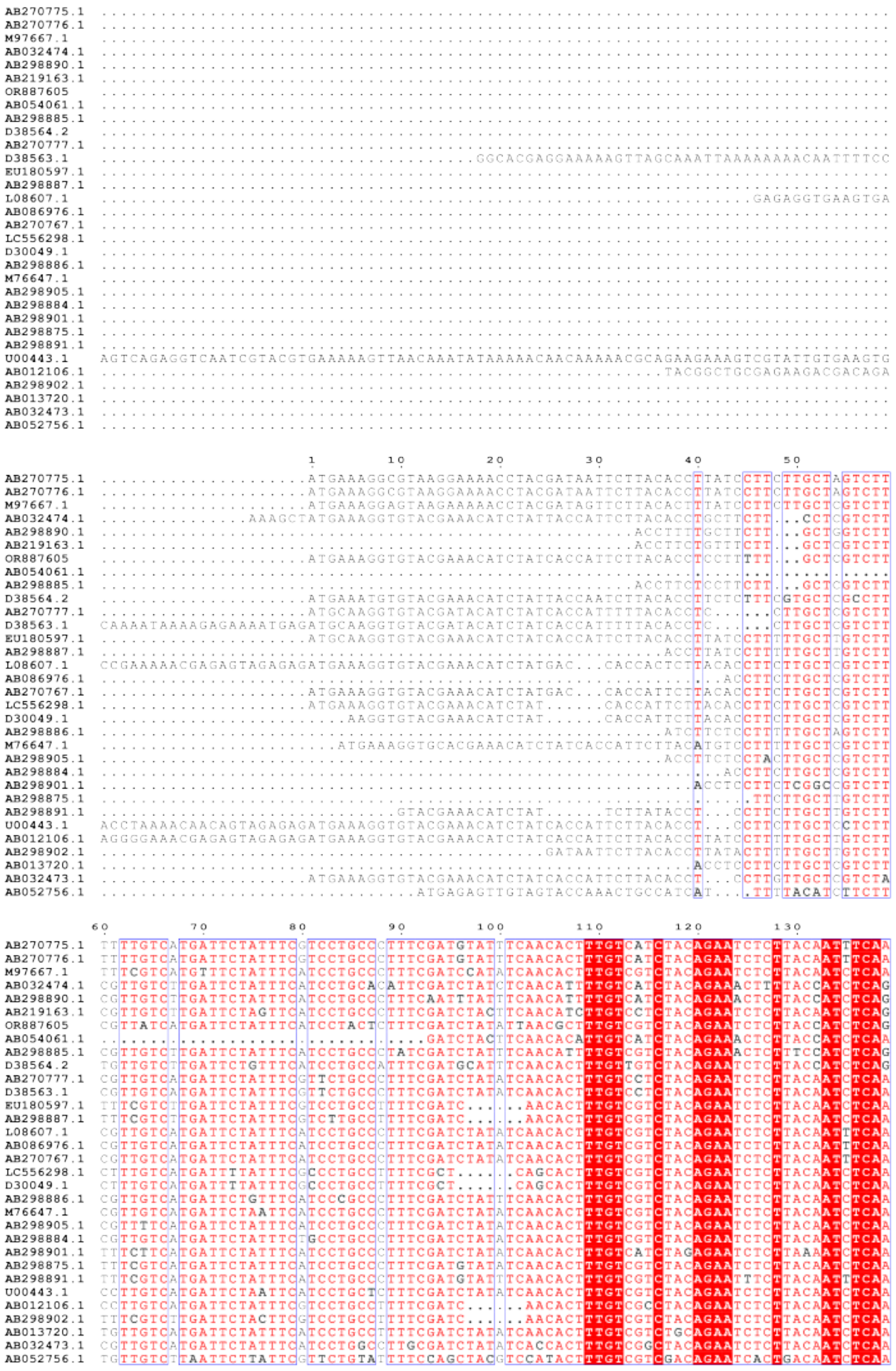

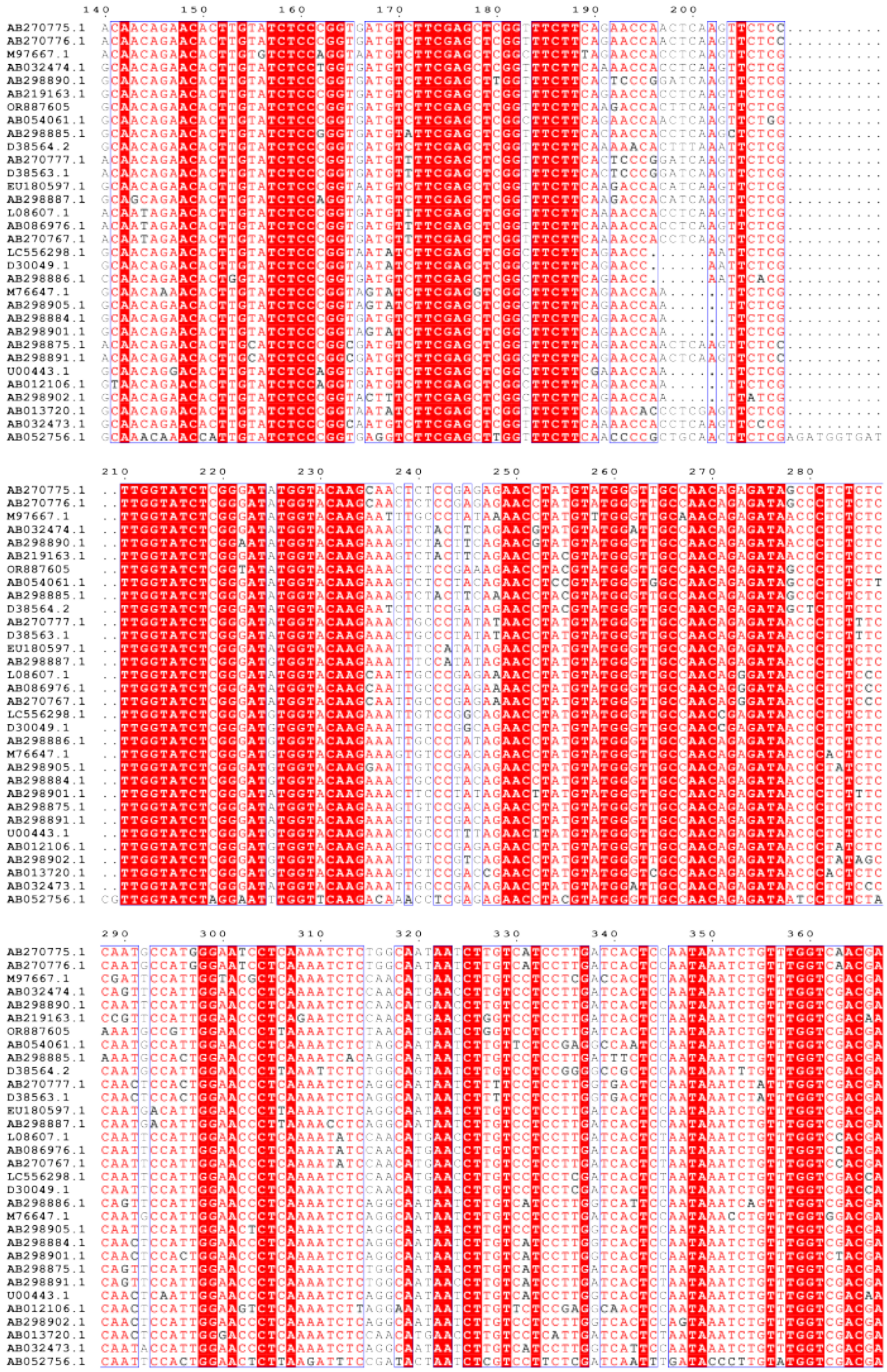

24

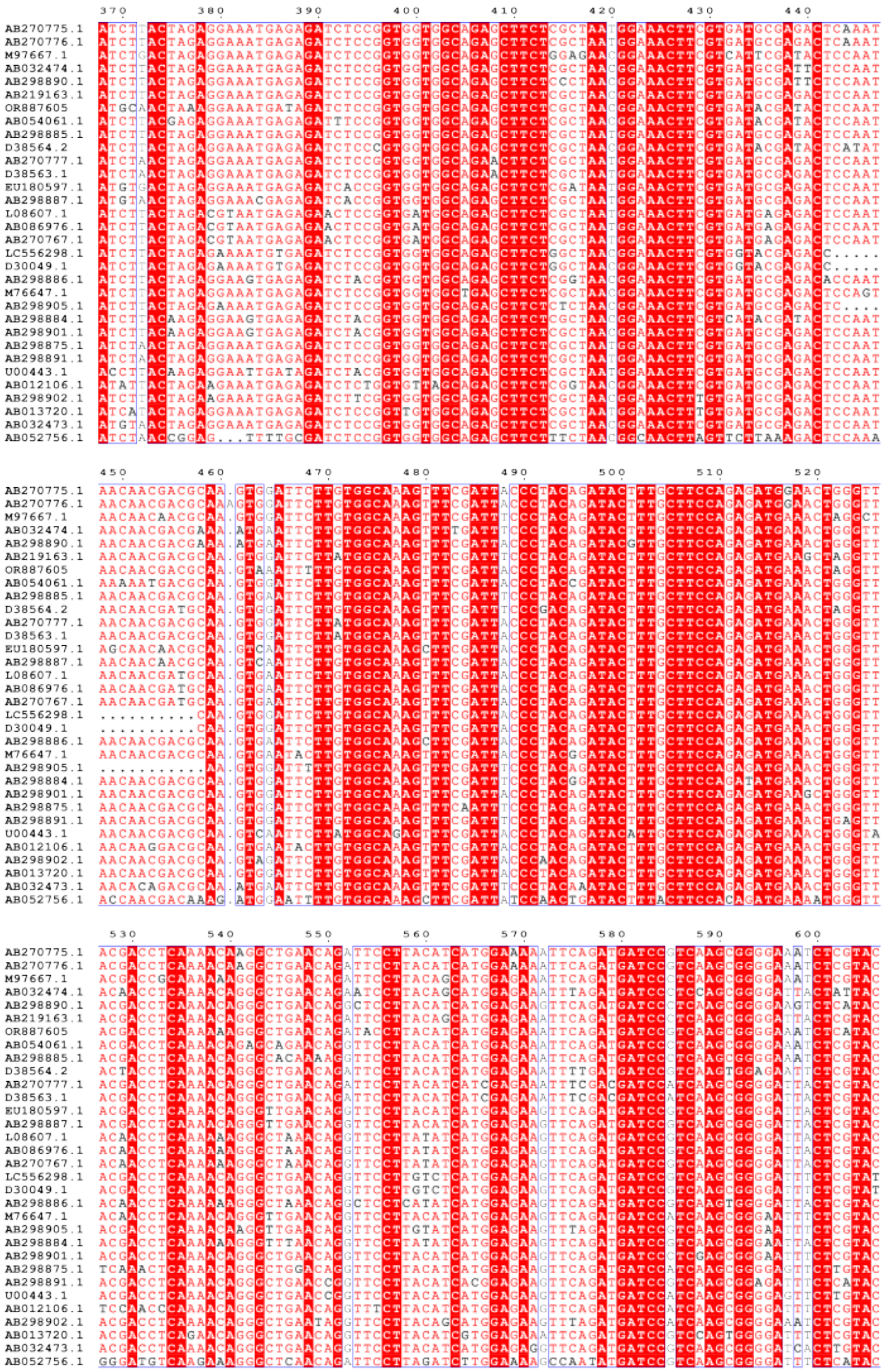

25

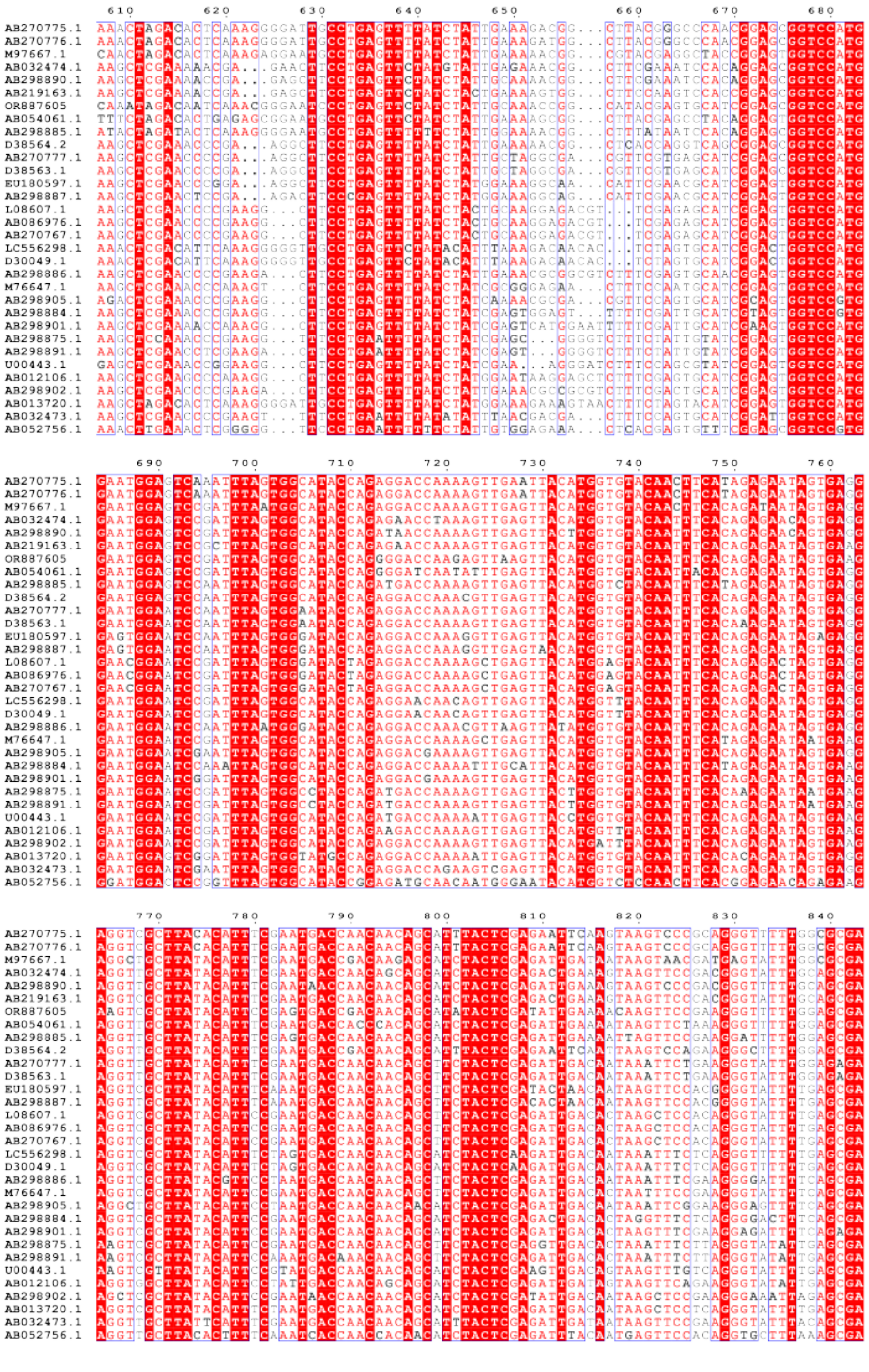
AB270775.1
AB270776.1
M97667.1
AB032474.1
AB298890.1
AB219163.1
OR887605
AB054061.1
AB298885.1
D38564.2
AB270777.1
D38563.1
EU180597.1
AB298887.1
L08607.1
AB086976.1
AB270767.1
LC556298.1
D30049.1
AB298886.1
M76647.1
AB298905.1
AB298884.1
AB298901.1
AB298875.1
AB298891.1
U00443.1
AB012106.1
AB298902.1
AB013720.1
AB032473.1
AB052756.1

26

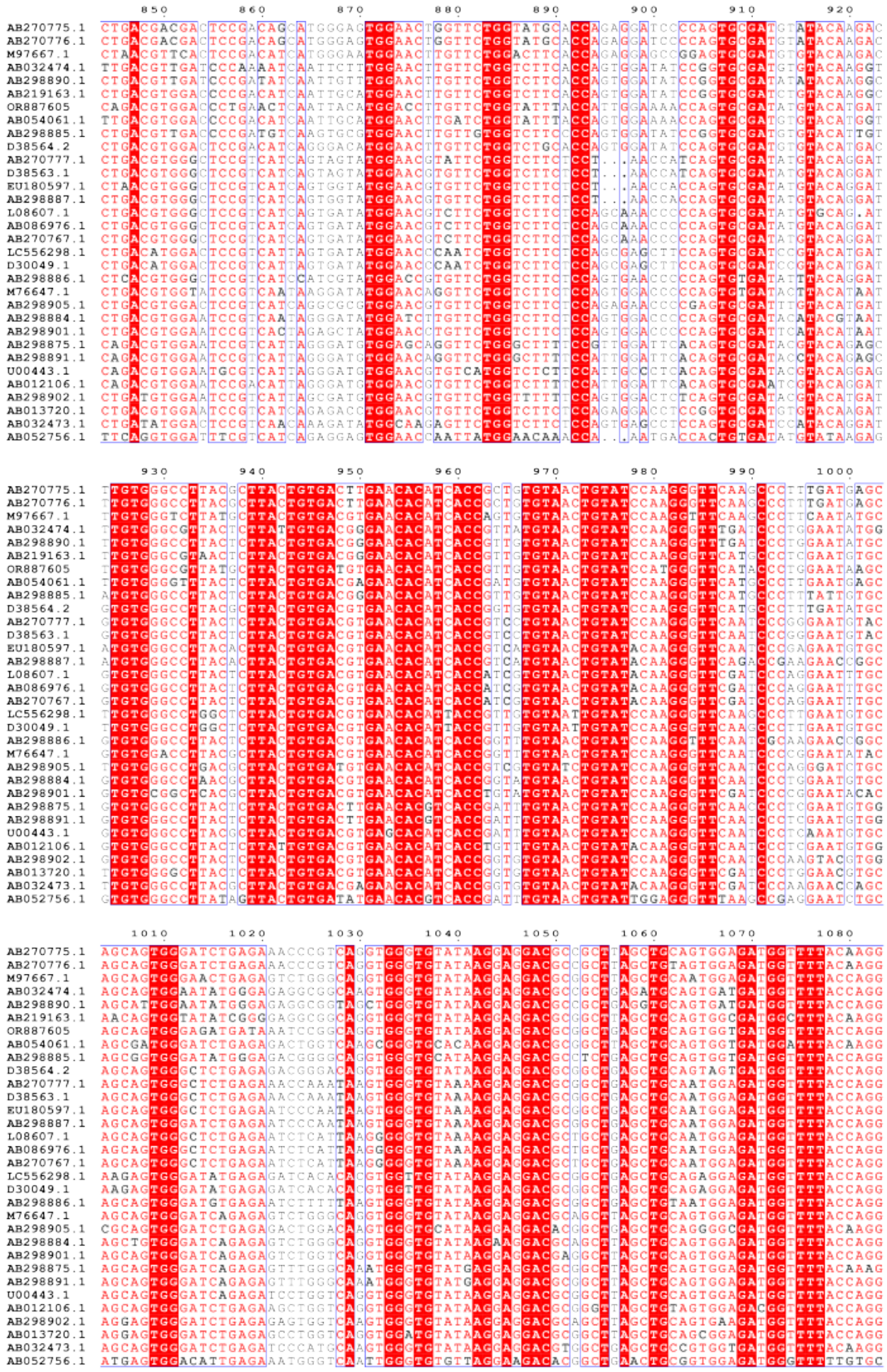

27

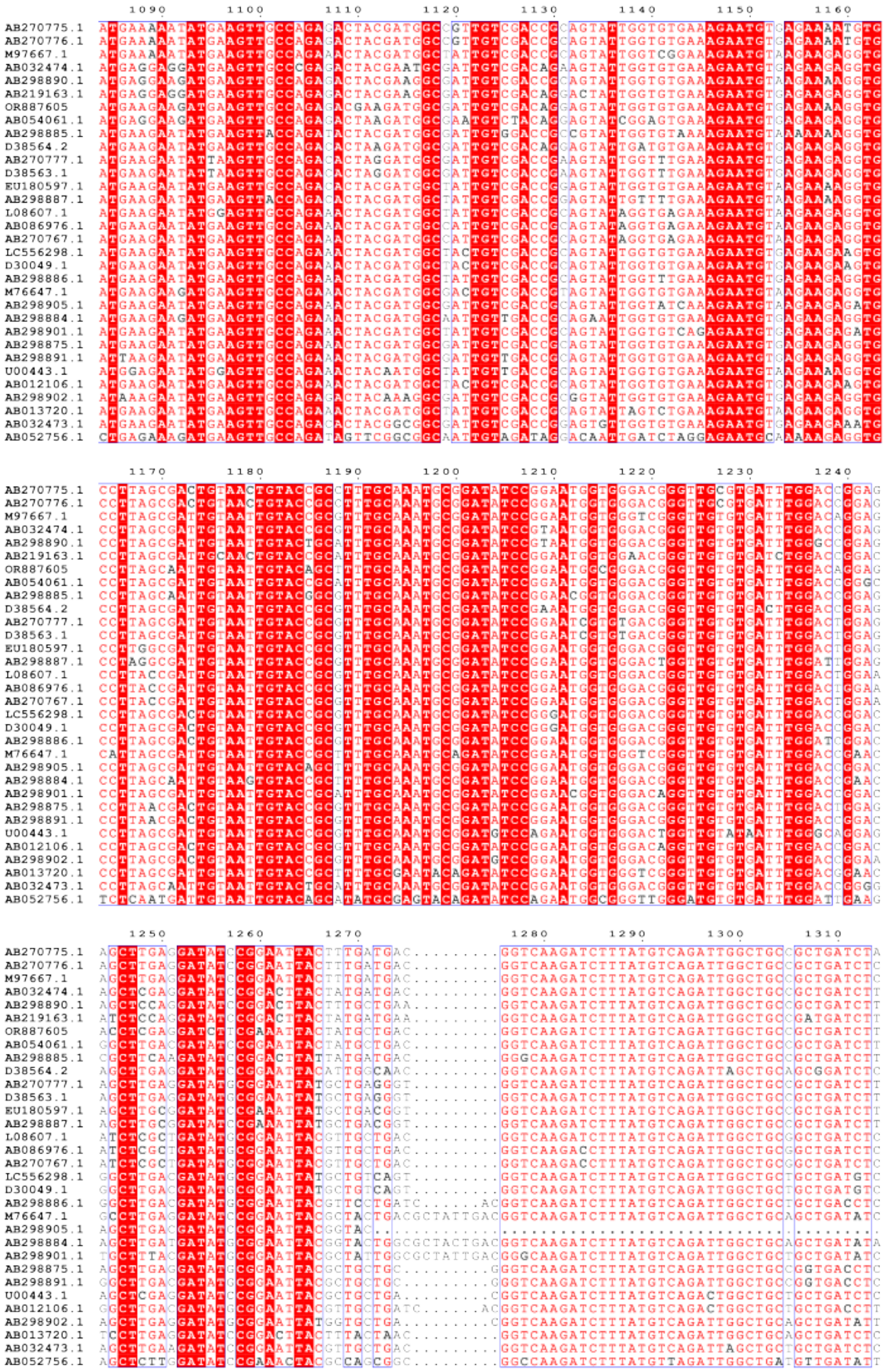

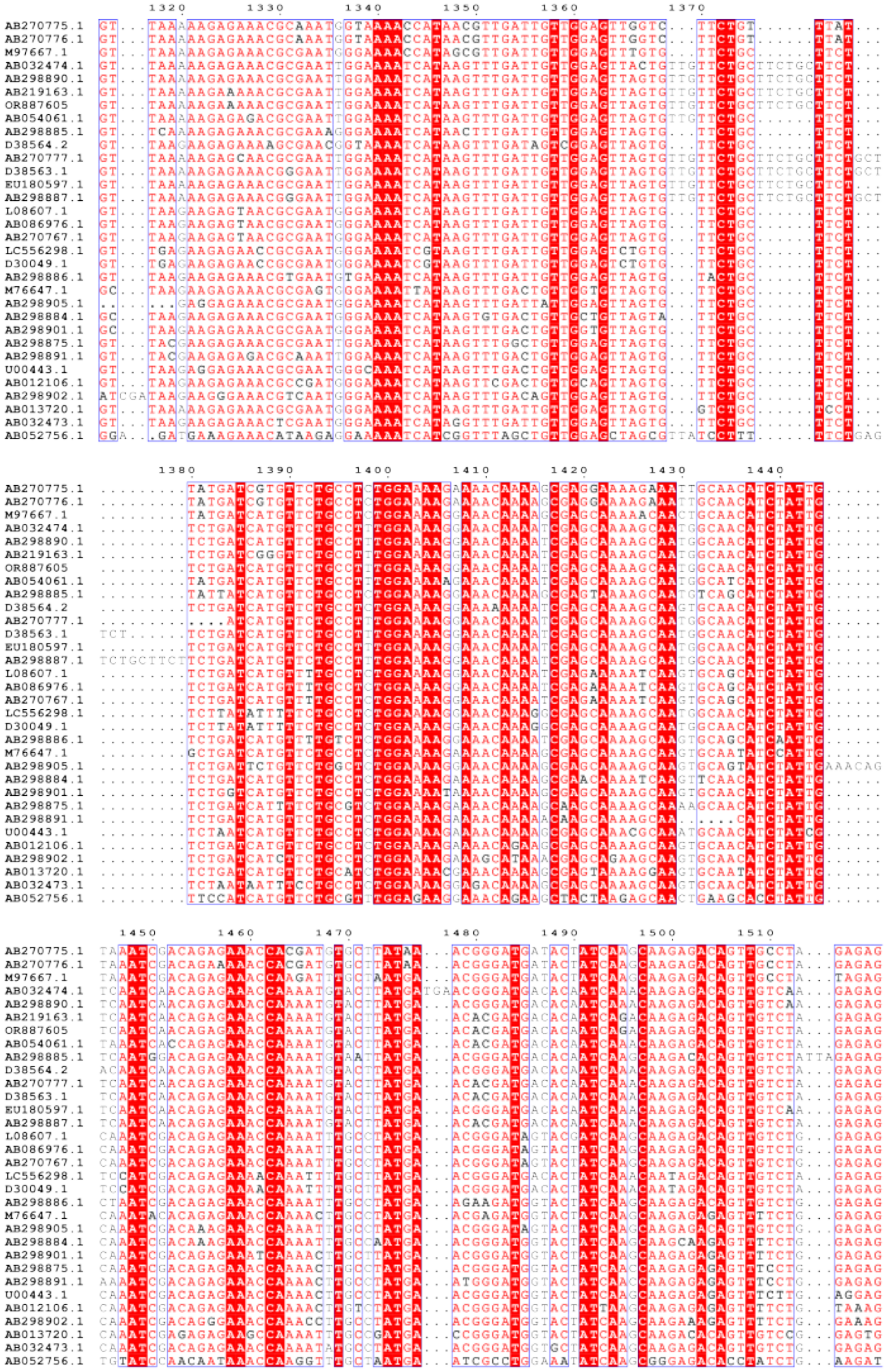

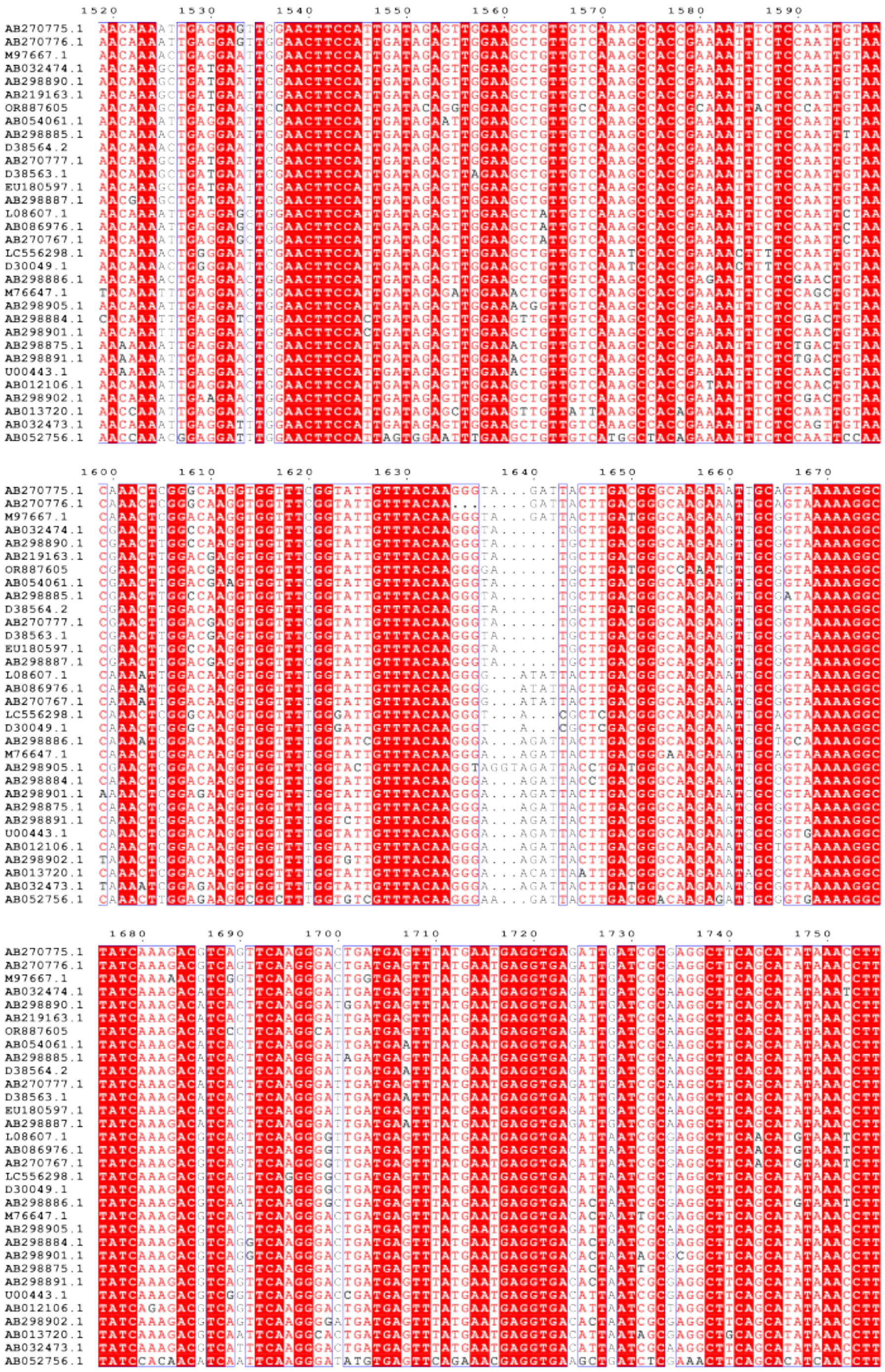
1520 1530 1540 1550 1560 1570 1580 1590
1600 1610 1620 1630 1640 1650 1660 1670
1680 1690 1700 1710 1720 1730 1740 1750
AB270775.1
AB270776.1
M97667.1
AB032474.1
AB298890.1
AB219163.1
OR887605
AB054061.1
AB298885.1
D38564.2
AB270777.1
D38563.1
EU180597.1
AB298887.1
L08607.1
AB086976.1
AB270767.1
LC556298.1
D30049.1
AB298886.1
M76647.1
AB298905.1
AB298884.1
AB298901.1
AB298875.1
AB298891.1
U00443.1
AB012106.1
AB298902.1
AB013720.1
AB032473.1
AB052756.1

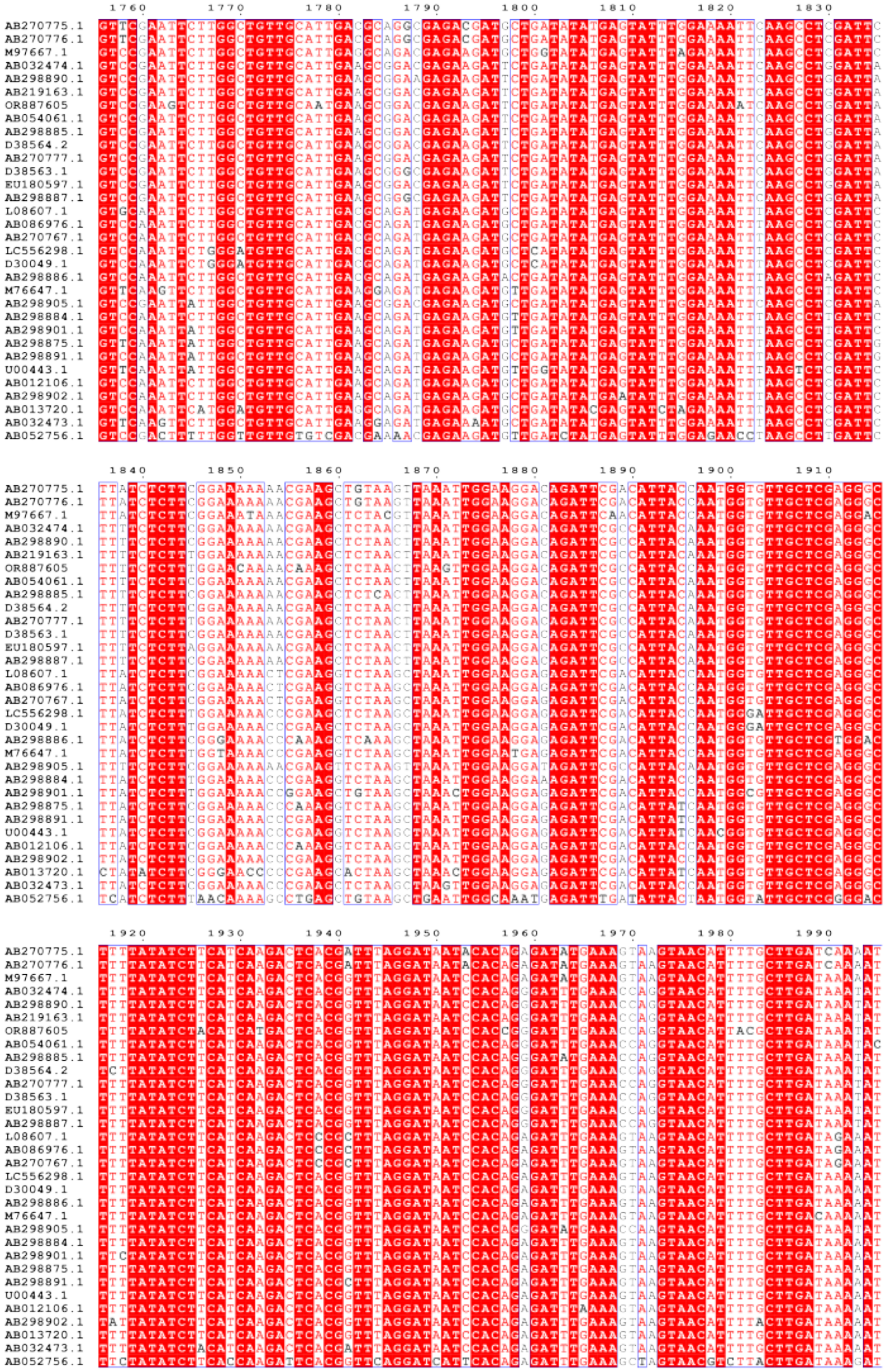
1760 1770 1780 1790 1800 1810 1820 1830
1840 1850 1860 1870 1880 1890 1900 1910
1920 1930 1940 1950 1960 1970 1980 1990
AB270775.1
AB270776.1
M97667.1
AB032474.1
AB298890.1
AB219163.1
OR887605
AB054061.1
AB298885.1
D38564.2
AB270777.1
D38563.1
EU180597.1
AB298887.1
L08607.1
AB086976.1
AB270767.1
LC556298.1
D30049.1
AB298886.1
M76647.1
AB298905.1
AB298884.1
AB298901.1
AB298875.1
AB298891.1
U00443.1
AB012106.1
AB298902.1
AB013720.1
AB032473.1
AB052756.1

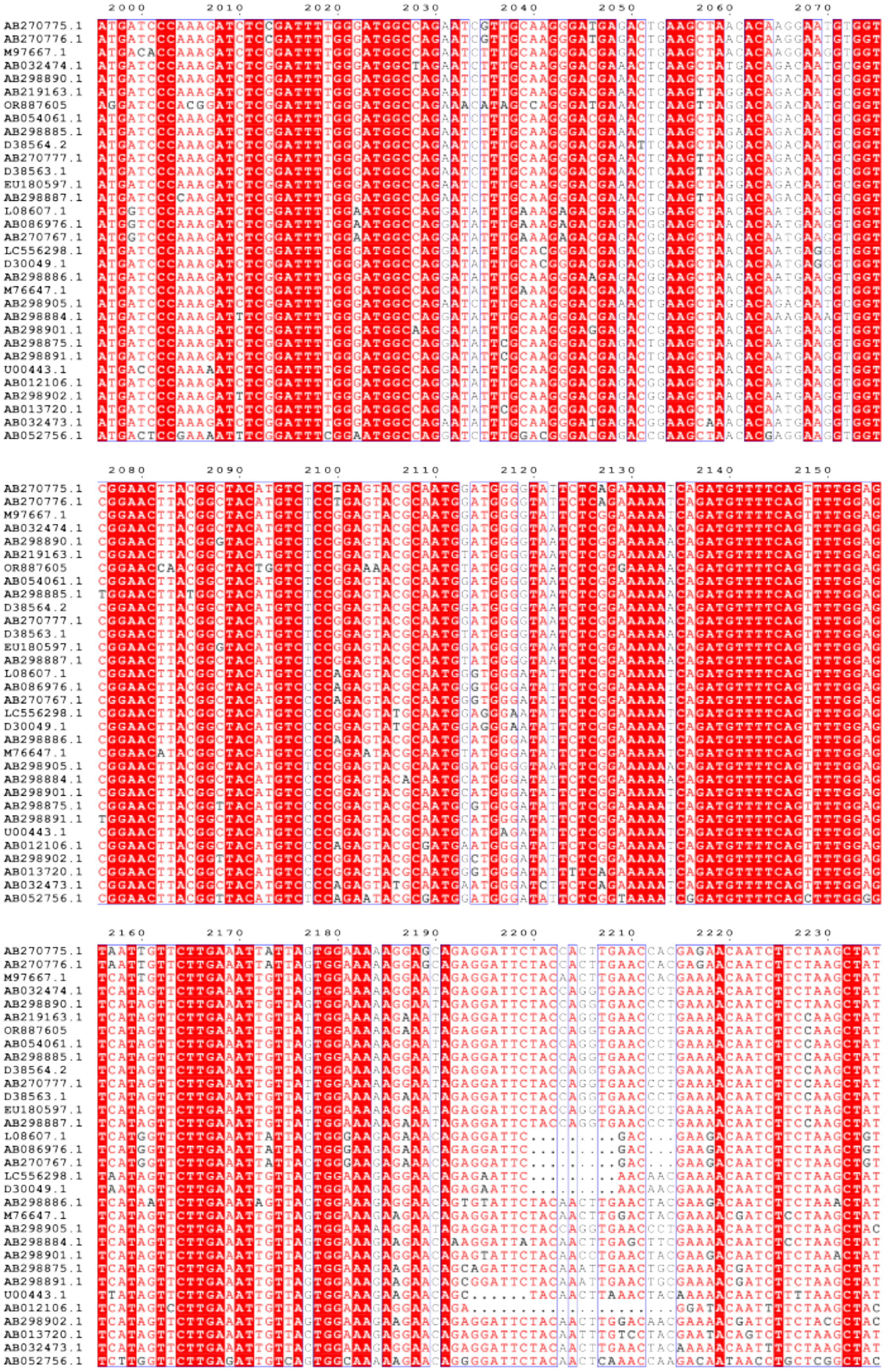
2000
2010
2020
2030
2040
2050
2060
2070
2080
2090
2100
2110
2120
2130
2140
2150
2160
2170
2180
2190
2200
2210
2220
2230
AB270775.1
AB270776.1
M97667.1
AB032474.1
AB298890.1
AB219163.1
OR887605
AB054061.1
AB298885.1
D38564.2
AB270777.1
D38563.1
EU180597.1
AB298887.1
L08607.1
AB086976.1
AB270767.1
LC556298.1
D30049.1
AB298886.1
M76647.1
AB298905.1
AB298884.1
AB298901.1
AB298875.1
AB298891.1
U00443.1
AB012106.1
AB298902.1
AB013720.1
AB032473.1
AB052756.1

32

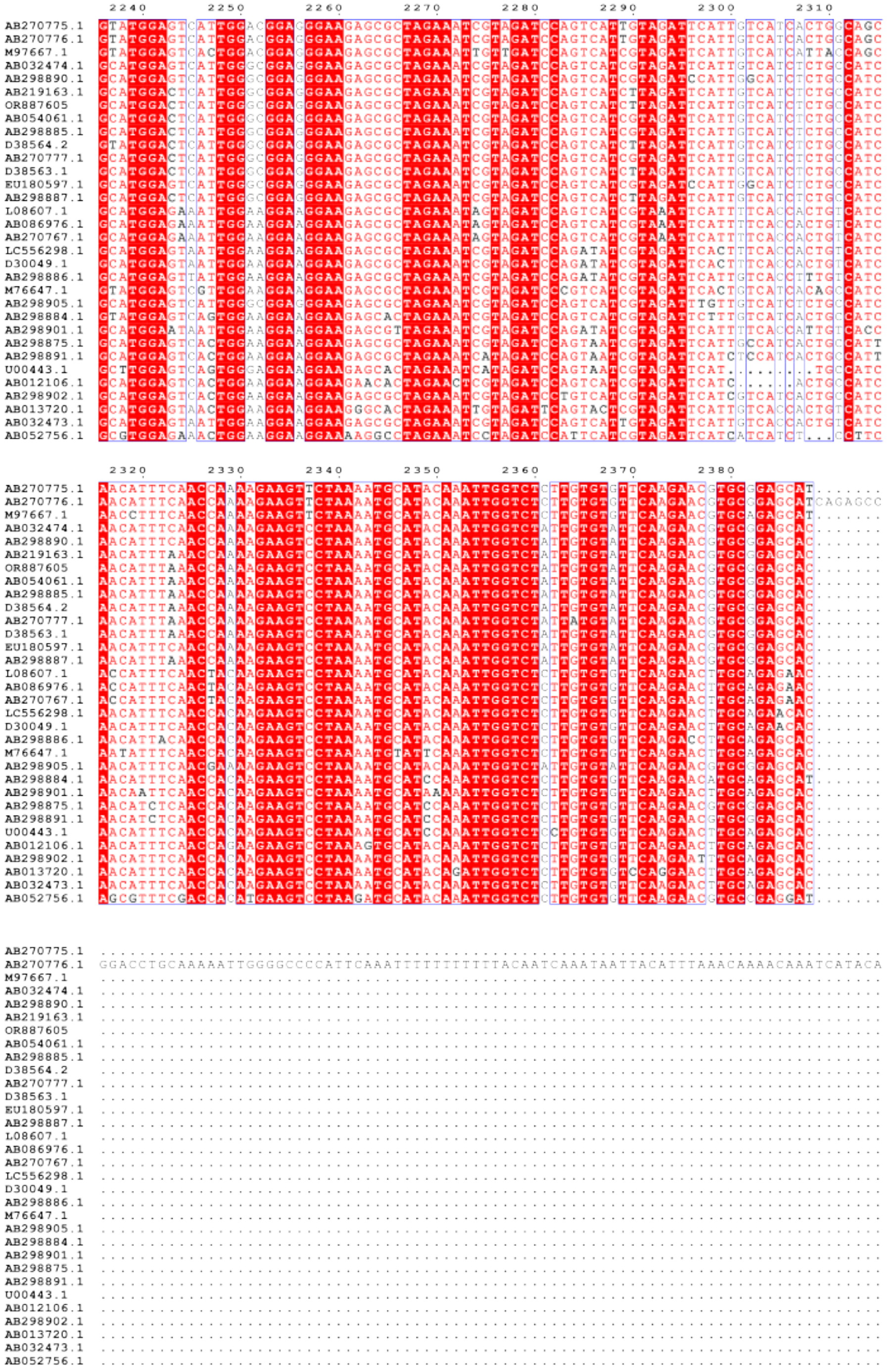

33

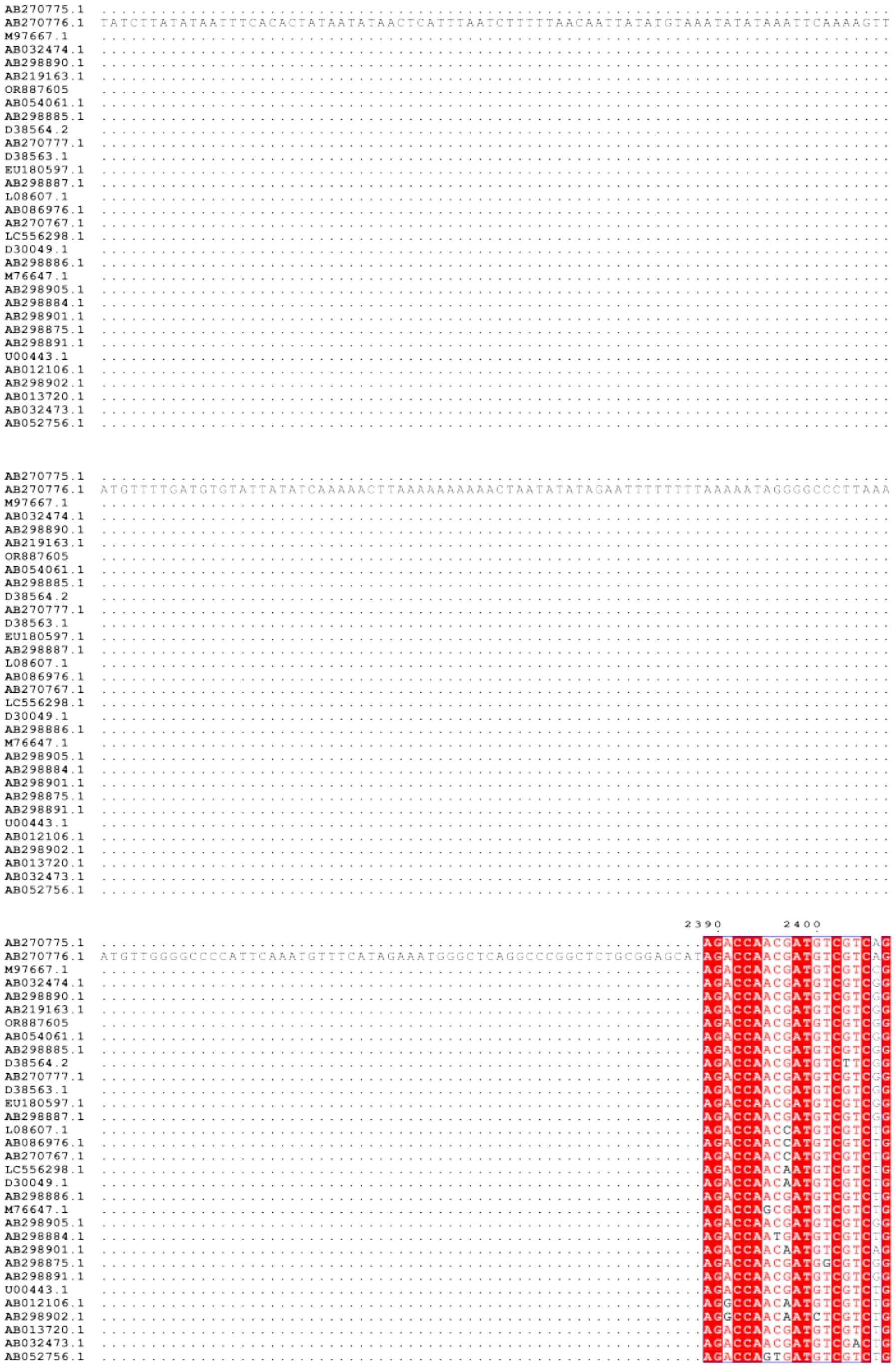
AB270775.1
AB270776.1
M97667.1
AB032474.1
AB298890.1
AB219163.1
OR887605
AB054061.1
AB298885.1
D38564.2
AB270777.1
D38563.1
EU180597.1
AB298887.1
L08607.1
AB086976.1
AB270767.1
LC556298.1
D30049.1
AB298886.1
M76647.1
AB298905.1
AB298884.1
AB298901.1
AB298875.1
AB298891.1
U00443.1
AB012106.1
AB298902.1
AB013720.1
AB032473.1
AB052756.1
2390
2400

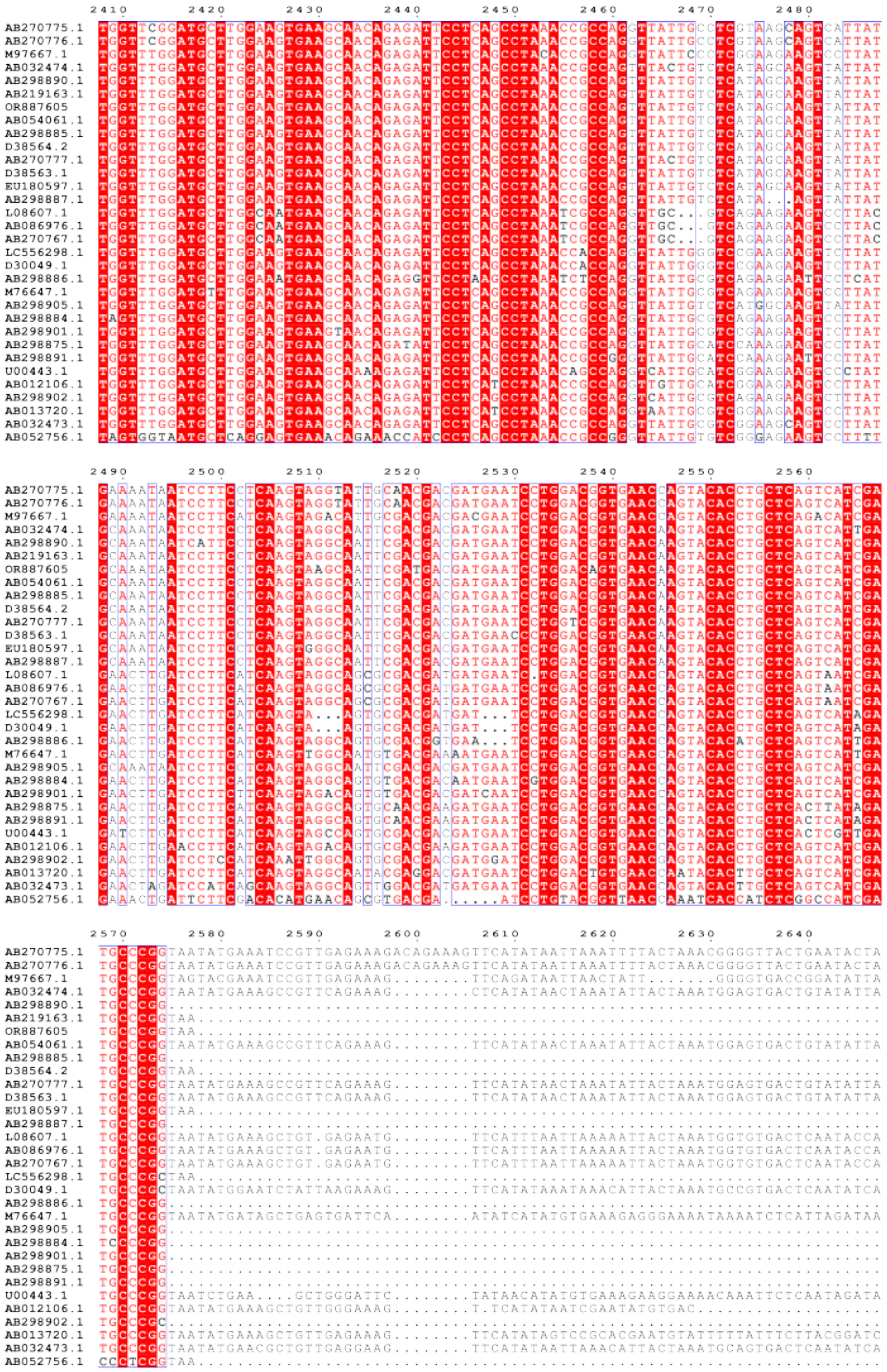

35

```
               2650      2660      2670      2680      2690      2700      2710      2720
AB270775.1  TGTGTGAAAGAAT.TAATATTTCAATAGATAAATTTCTTGTTATTTTGGGAAAAAGAATTCTTATTTTCATAACCAATTC
AB270776.1  TGTGTGAAAGAAT.TAATATTTCAATAGATAAATTTCTTGTTATTTTGGGAAAAAGAATTCTTATTTTCATAACCAATTC
M97667.1    TAAGTGAAAGAAAATAAAATTTCAATAGTTAAGT...TTGTTATTTGATAACCAAATCTTGTTATTTCCTGGTGGTGTTG
AB032474.1  TAAGTGATCGAAGGACAATAAATTCTCAGTAGATAAGTTTGTTATATTGATAACCAATTCATGTTATTTTCTGGTGAAGT
AB298890.1  ................................................................................
AB219163.1  ................................................................................
OR887605    ................................................................................
AB054061.1  TAAGTGATCGAAGGAAAATAAATTCTCAGCAGATAAGTTTGTTATATTGATAACCAATT.............GGTGAAGT
AB298885.1  ................................................................................
D38564.2    ................................................................................
AB270777.1  TAAGTGAA........................................................................
D38563.1    TAAGTGATCGAAGGAAAATAAATTCTCAGTAGATAAGTTTGTTATATTGATAACCAATTCTTGTTATTTTCTGGTGAAGT
EU180597.1  ................................................................................
AB298887.1  ................................................................................
L08607.1    TATGTGAAAGAAGTAAAATAAATTCTCAATAGAAA.............................................
AB086976.1  CATGTGAAAGAA....................................................................
AB270767.1  TATGTGAAAGAAGTAAAATAAATTCTCAATAGATAAAGTATGTTATTTTGATAACAAATTCTTGCGTCTTTTTTCTGGCG
LC556298.1  ................................................................................
D30049.1    CATGTGAAAGAAGGAAAATAAATTCTTAATAGTTAA.GTATGTTATTTTGATAACAAATTCTTGTTTTTTCTAGCTGTAT
AB298886.1  ................................................................................
M76647.1    GTAGGTTATTTCGATAACCACTTCTTGTTATTTTCTGGCGGTGTTGTCATTATCCCCTTTATATTAAAAAGAAGCATTTG
AB298905.1  ................................................................................
AB298884.1  ................................................................................
AB298901.1  ................................................................................
AB298875.1  ................................................................................
AB298891.1  ................................................................................
U00443.1    AGTATGTTATTTTGATGACCTATTTTTGTTATTTTCTTGCGGTGTTGTCATTATTCAAAATCTATAATAACACATTATTG
AB012106.1  ................................................................................
AB298902.1  ................................................................................
AB013720.1  AAAATTTAATGCATGATAAAATTGTTATATACTATTTAAAAATTGATGATATATATTTTATACTGA..............
AB032473.1  TATGTGAAAGAAGGAAATAAATTCTCAAAATATAAGTATGTTATTTTGTAAC............................
AB052756.1  ................................................................................

                2730      2740      2750      2760      2770      2780      2790
AB270775.1  TTGTTATTTTCTGGTGGTGTTCATATTCAAAGTACCATATTAAATGATTCCGGTTCCCCTATTA................
AB270776.1  TTGTTATTTTCTGGTGGTGTTCATATTCAAAGTACCA...........................................
M97667.1    TCATATTCGTTTTTCTGAATGAATGTTAAAGTTATTATTC........................................
AB032474.1  TGTCATTATTCATAGTACAATAATACATGCTGGAGCGCCTTGTCGGCACAAATGCGTGTTAGCTTTTTGTTTTGTGTGAT
AB298890.1  ................................................................................
AB219163.1  ................................................................................
OR887605    ................................................................................
AB054061.1  TGTCATTATTCATAGTACAATAATACATGCTGGAGCGTCTTGTCGGCACAAAAAAA........................
AB298885.1  ................................................................................
D38564.2    ................................................................................
AB270777.1  ................................................................................
D38563.1    TGTCATTATTCAAAGTACAATAATAAATGCTGGAGCGTCTTGTCGGC.................................
EU180597.1  ................................................................................
AB298887.1  ................................................................................
L08607.1    ................................................................................
AB086976.1  ................................................................................
AB270767.1  GTTCAAAGTTTTTTTTTTTCCTCGATGATATTAAACCAAATTGAAAGGACTACAAGGTCGGATACGTTTTTAGGACACCA
LC556298.1  ................................................................................
D30049.1    CTATCTTATTAAAATATAATTACACTTAT...................................................
AB298886.1  ................................................................................
M76647.1    TATTAAATCCCCTTGCCTCAAGAGATATTCACAAGAATACTATTGTGACGTGACAGCCTCACTATCGTTTAAACATTACA
AB298905.1  ................................................................................
AB298884.1  ................................................................................
AB298901.1  ................................................................................
AB298875.1  ................................................................................
AB298891.1  ................................................................................
U00443.1    C...............................................................................
AB012106.1  ................................................................................
AB298902.1  ................................................................................
AB013720.1  ................................................................................
AB032473.1  ................................................................................
AB052756.1  ................................................................................

AB270775.1  .....................................................
AB270776.1  .....................................................
M97667.1    .....................................................
AB032474.1  ACAGATAATTGTGAGATATTCAAACTGCAACCGCTCATTTTTTAAGTTGTTTG
AB298890.1  .....................................................
AB219163.1  .....................................................
OR887605    .....................................................
AB054061.1  .....................................................
AB298885.1  .....................................................
D38564.2    .....................................................
AB270777.1  .....................................................
D38563.1    .....................................................
EU180597.1  .....................................................
AB298887.1  .....................................................
L08607.1    .....................................................
AB086976.1  .....................................................
AB270767.1  AAACCAGCTTTCAAACAGATGAAACTGGAGACATAAGCTCCTA..........
LC556298.1  .....................................................
D30049.1    .....................................................
AB298886.1  .....................................................
M76647.1    ATGCTGACGTGTGGCTTGTAAATAGCTTCTCAGACC.................
AB298905.1  .....................................................
AB298884.1  .....................................................
AB298901.1  .....................................................
AB298875.1  .....................................................
AB298891.1  .....................................................
U00443.1    .....................................................
AB012106.1  .....................................................
AB298902.1  .....................................................
AB013720.1  .....................................................
AB032473.1  .....................................................
AB052756.1  .....................................................
```

**Multiple sequence alignment of gene sequences of *FER1* selected for phylogenetic study**

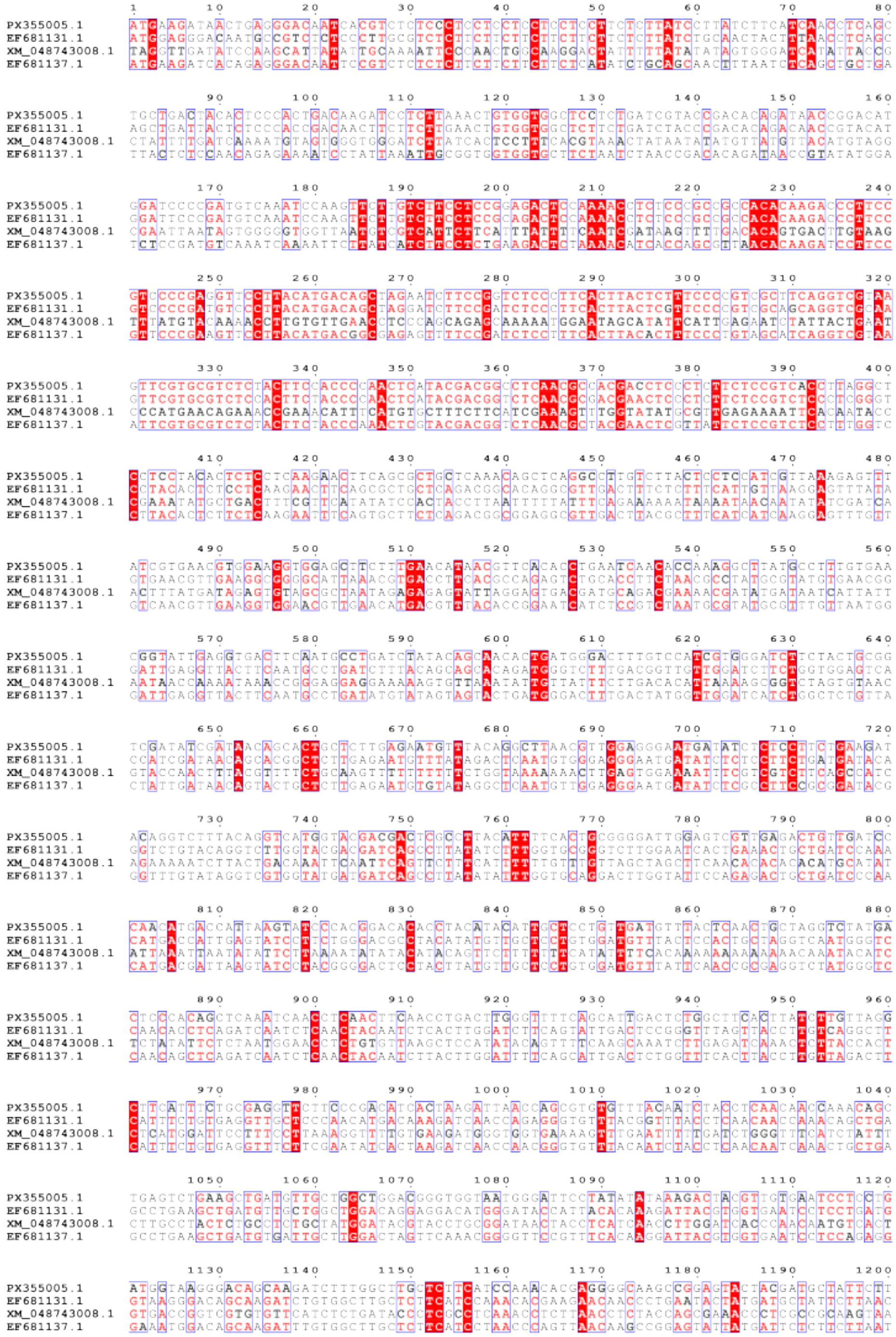

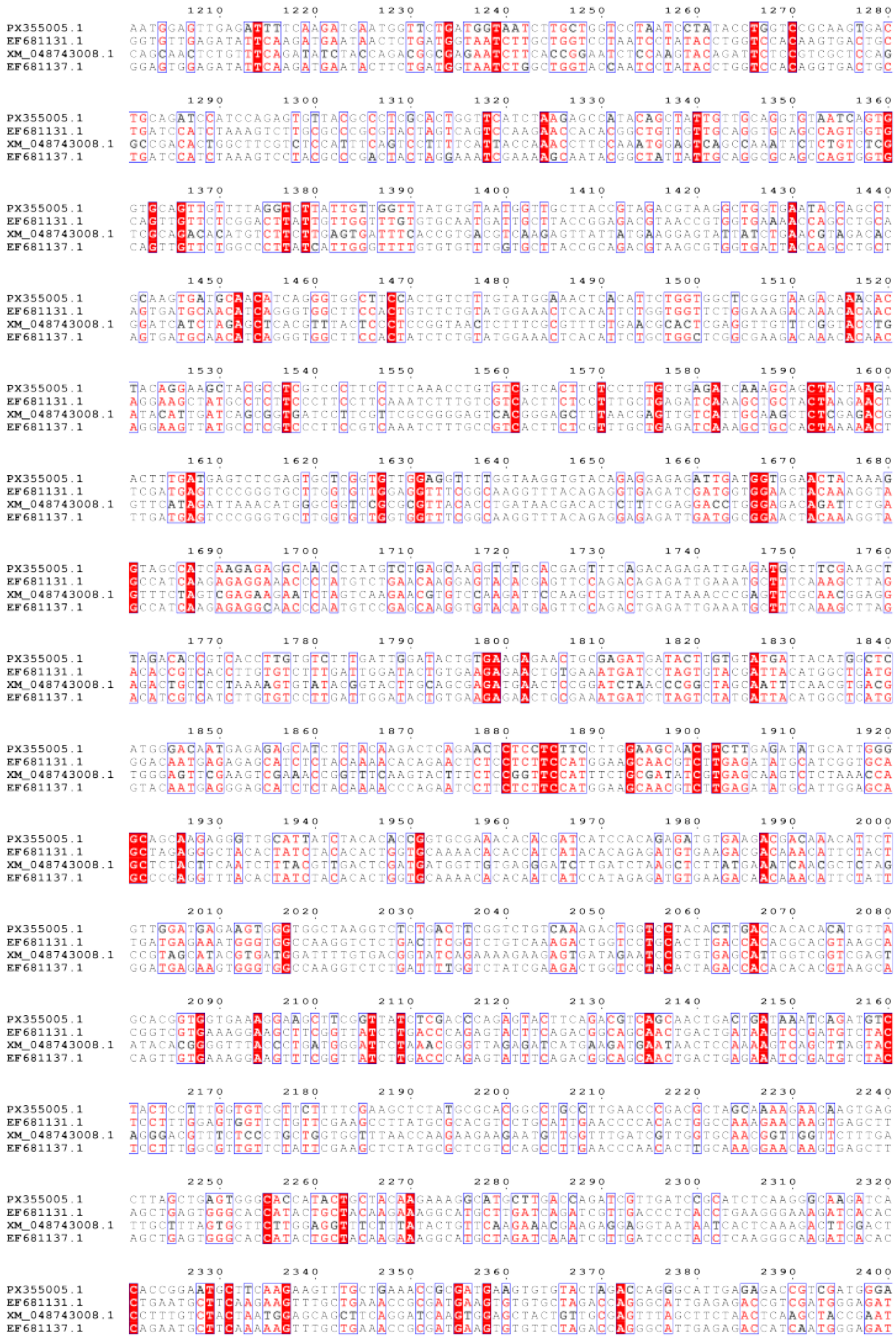


40

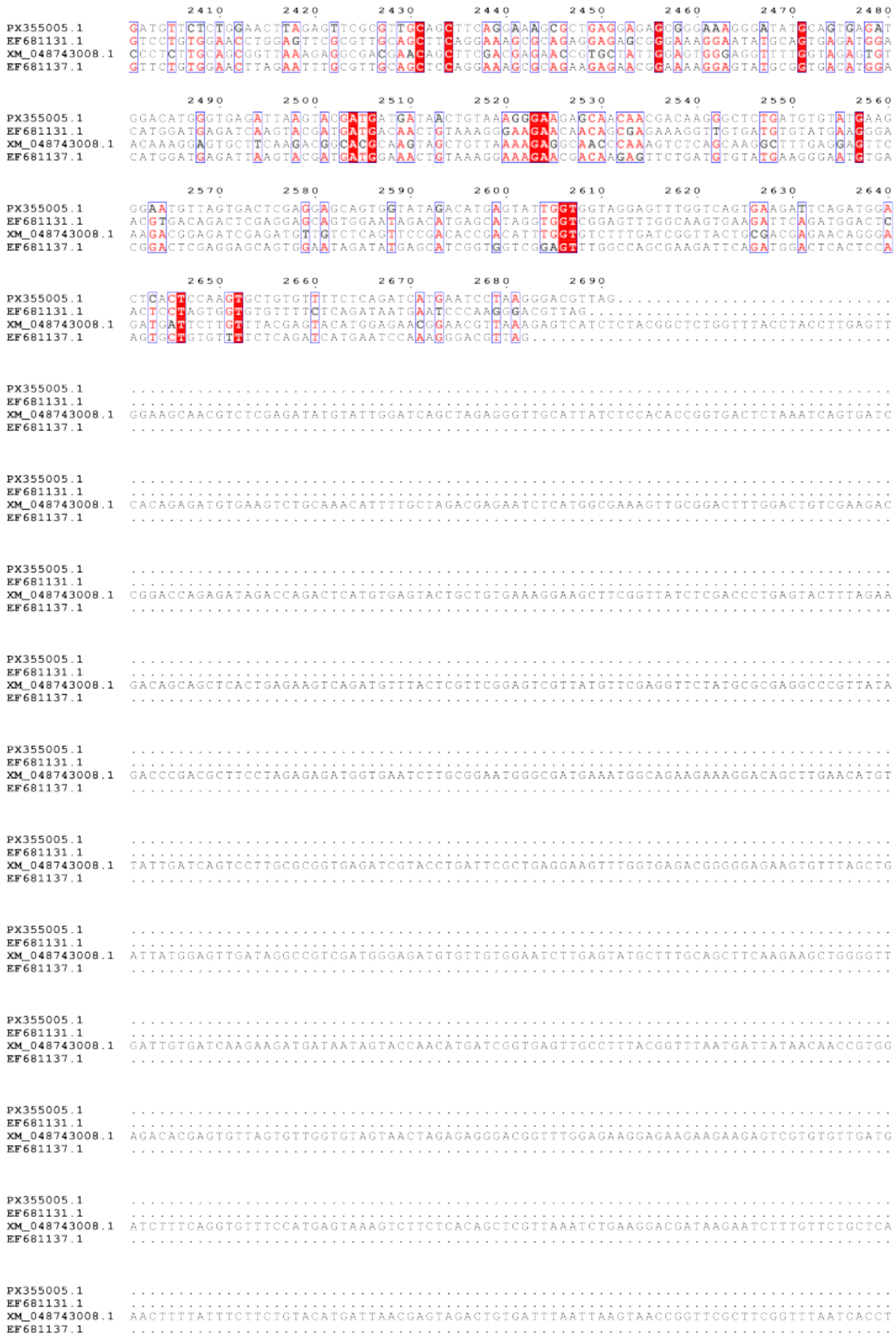
2410 2420 2430 2440 2450 2460 2470 2480
2490 2500 2510 2520 2530 2540 2550 2560
2570 2580 2590 2600 2610 2620 2630 2640
2650 2660 2670 2680 2690
PX355005.1
EF681131.1
XM_048743008.1
EF681137.1

**Multiple sequence alignment of gene sequences of *MLPK* selected for phylogenetic study**

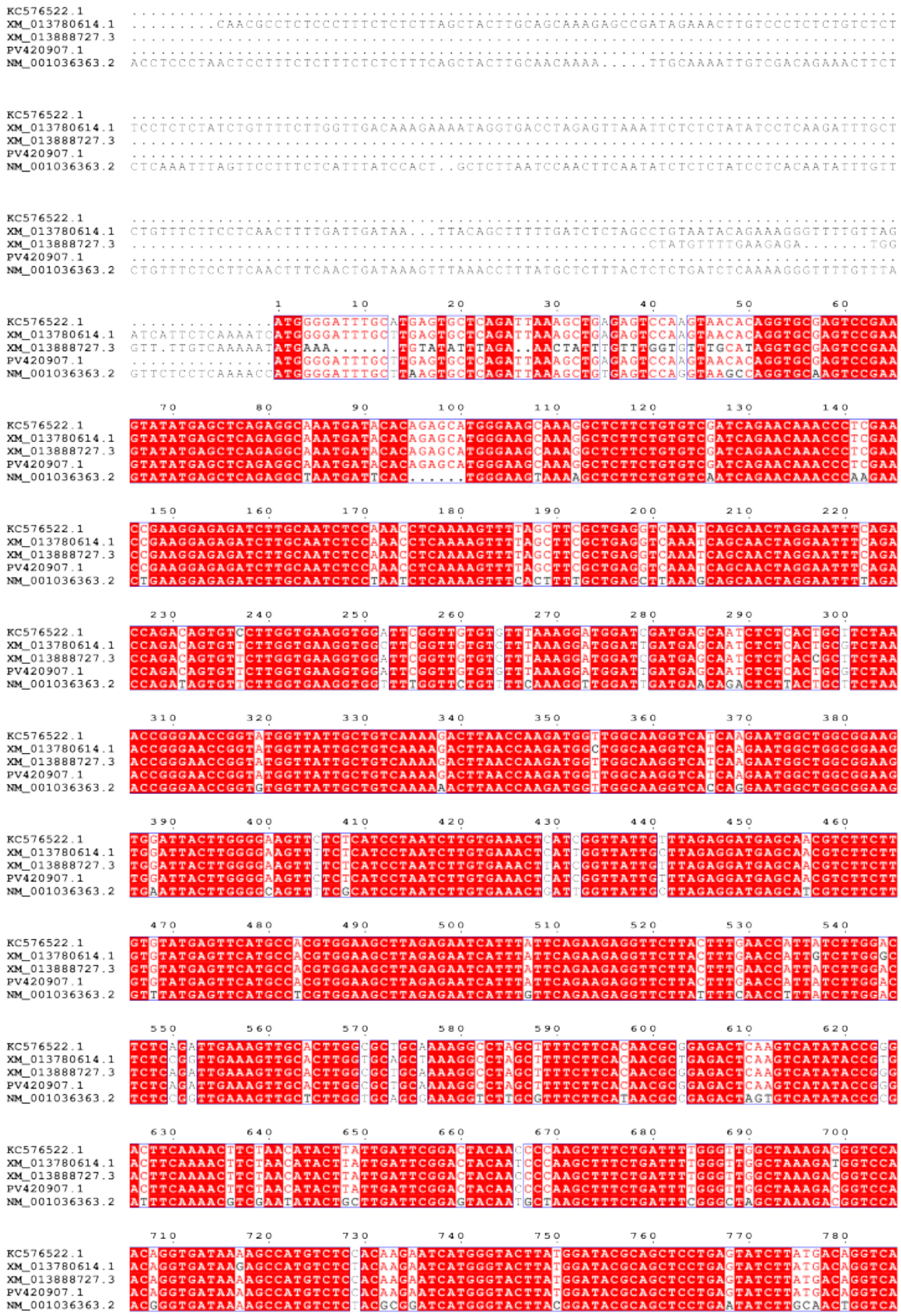

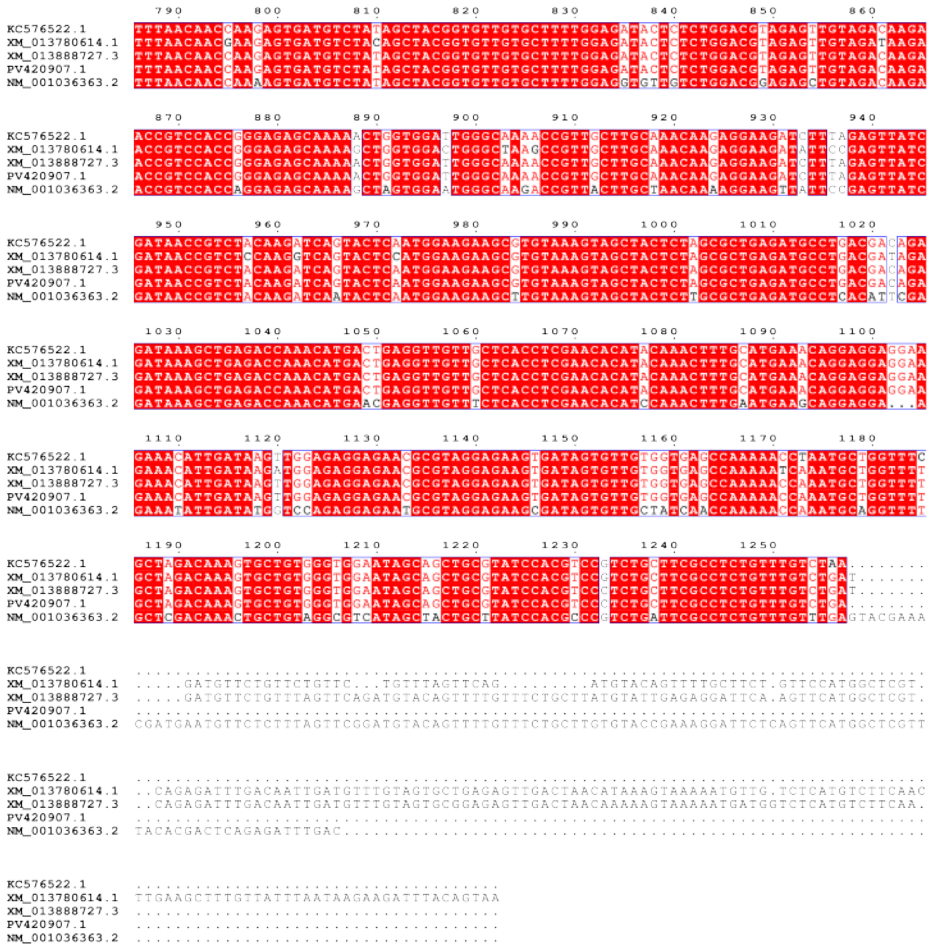

**Multiple sequence alignment of gene sequences of *ARC1* selected for phylogenetic study**

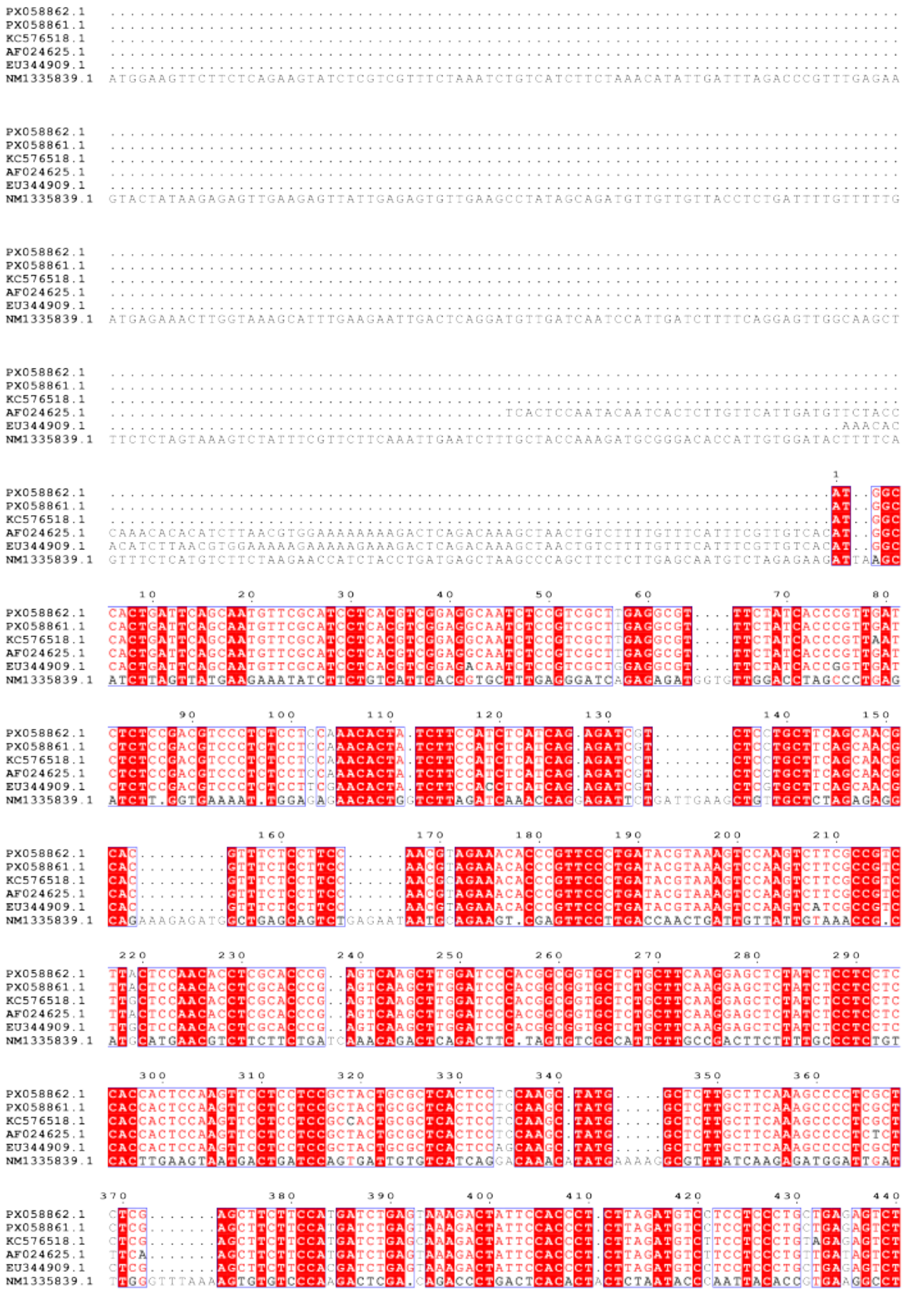

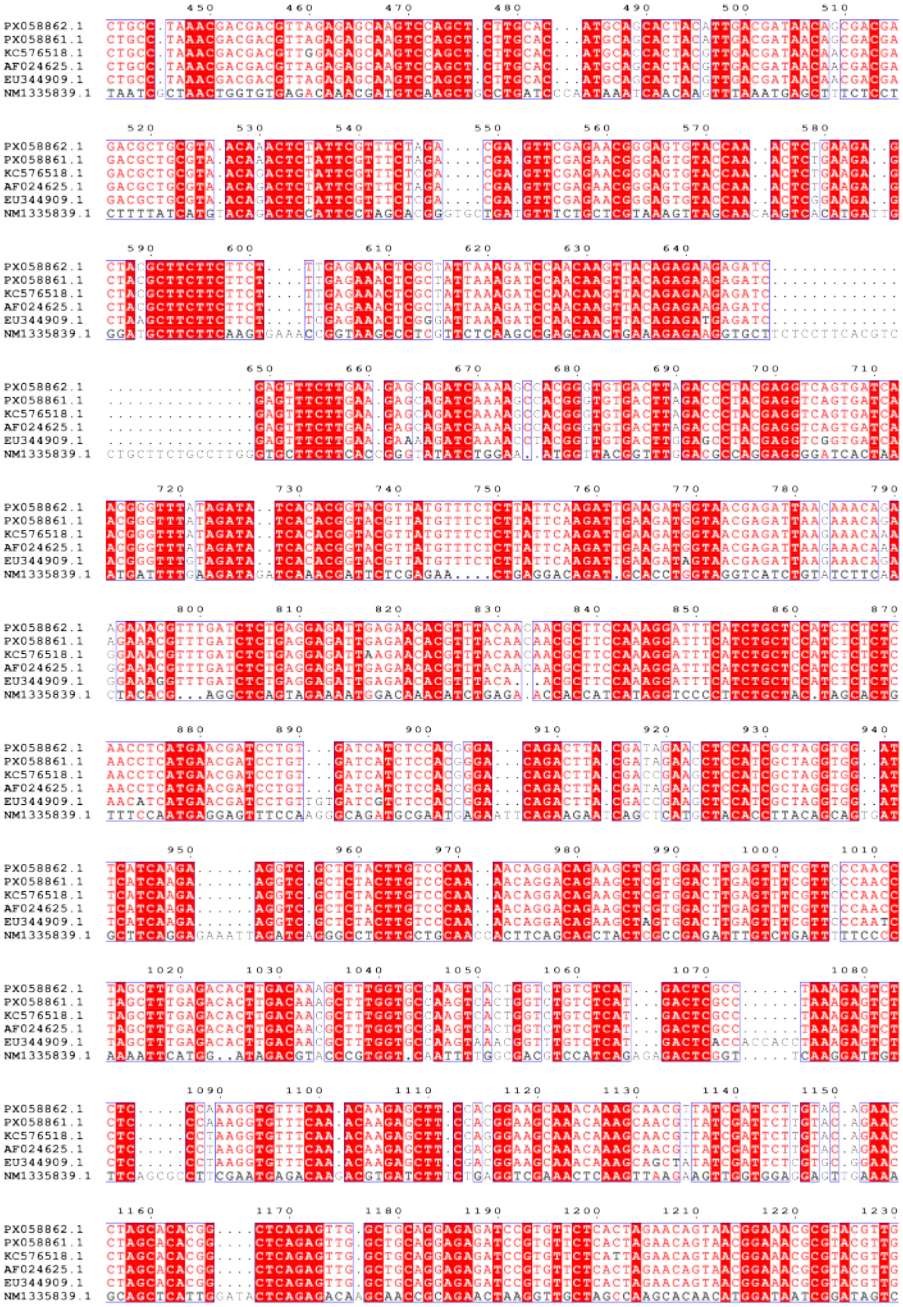
PX058862.1
PX058861.1
KC576518.1
AF024625.1
EU344909.1
NM1335839.1

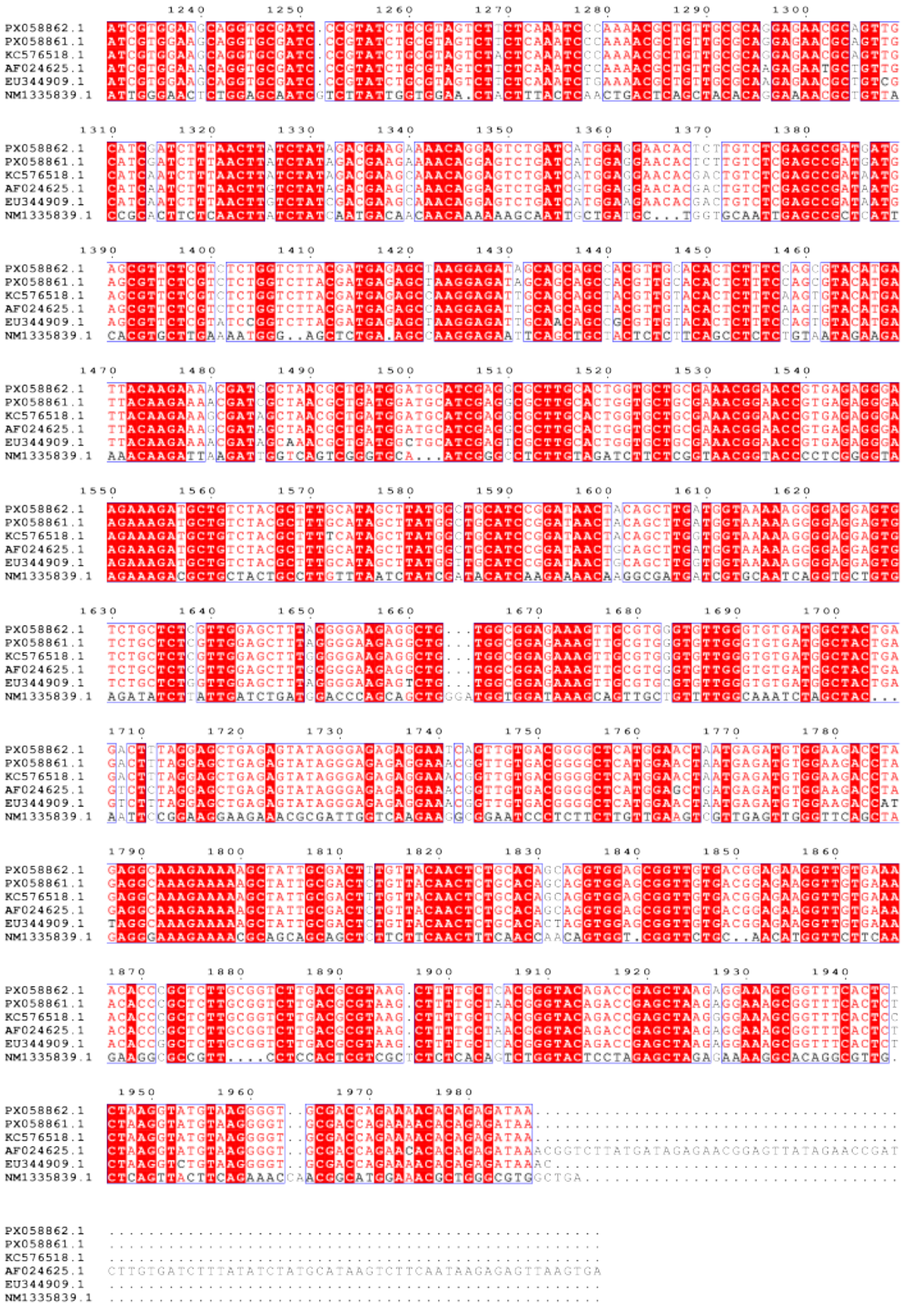

50
